# Protracted Maturation of Proactive and Reactive Systems Predicts Cognitive Stability and Psychopathology : A longitudinal multi-cohort study

**DOI:** 10.64898/2026.07.07.736457

**Authors:** Zhiyao Gao, Li Zheng, Tobias Banaschewski, Gareth J. Barker, Arun L.W. Bokde, Rüdiger Brühl, Sylvane Desrivières, Penny Gowland, Antoine Grigis, Andreas Heinz, Frauke Nees, Dimitri Papadopoulos Orfanos, Luise Poustka, Michael N. Smolka, Sarah Hohmann, Nathalie Holz, Nilakshi Vaidya, Henrik Walter, Robert Whelan, Paul Wirsching, Gunter Schumann, Hugh Garavan, Vinod Menon, Weidong Cai, the IMAGEN Consortium

## Abstract

Inhibitory control matures progressively from childhood to early adulthood, yet the neural mechanisms driving this development and their relevance to psychiatric risk remain poorly understood. Guided by the Dual Cognitive Control model, we leveraged longitudinal fMRI from two independent cohorts in the US (ABCD, ages 9–12) and Europe (IMAGEN, ages 14–22) to map the spatiotemporal dynamics of reactive and proactive control using novel single-trial modeling and representational similarity analysis. We found both reactive and proactive stopping networks stabilize after mid-adolescence, tracking the developmental patterns of inhibitory control and behavioral stability. By decoding trial-by-trial fluctuations along a speed-caution continuum, we demonstrate that brain-behavior coupling to a proactive “Safe state” tightens progressively with age. Furthermore, network-level representational coherence of this Safe state emerged as a scanner-invariant, trait-like biomarker that robustly predicted inhibitory control, behavioral stability, and transdiagnostic psychopathology across multiple developmental windows, providing a validated neural phenotype for precision psychiatry.

## Introduction

Inhibitory control, the capacity to suppress inappropriate actions, thoughts, and emotions, serves as a cornerstone of goal-directed behavior and mental well-being ^1-3^. Deficits in this domain are transdiagnostic, manifesting across a broad spectrum of neuropsychiatric conditions, including attention-deficit hyperactivity disorder (ADHD) and obsessive-compulsive disorder (OCD) ^4-6^. Although inhibitory capacity improves steadily through childhood and adolescence, a period of profound neural reorganization ^7-15^ and psychiatric vulnerability ^16-22^, the functional brain mechanisms driving this maturation remain poorly understood.

A key theoretical advance is the Dual Cognitive Control (DCC) model, which distinguishes between reactive control (a stimulus-driven, transient stopping mechanism) and proactive control (a goal-driven, anticipatory state that sustains inhibition across trials) ^23,24^. The DCC model posits that control is dynamically allocated trial-by-trial, with individuals constantly re-weighting a “speed-safety” tradeoff between rapid execution and cautious restraint. Regulating this tradeoff is thought to be the hallmark of mature cognitive control, making its developmental trajectory essential for understanding transdiagnostic psychiatric vulnerability ^25,26^.

To date, developmental research has predominantly tracked reactive control, typically indexed by the rapid shortening of the Stop-Signal Reaction Time (SSRT) in the Stop-signal Task (SST) ^8,9,27-29^, a developmental shift most pronounced during late childhood ^8,9^. In contrast, the protracted development of proactive control and behavioral stability has received far less attention ^14,25,30-33^.

Neuroimaging studies of inhibitory control have consistently implicated the joint recruitment of the Salience Network (SN), anchored in the anterior insula (AI)/inferior frontal gyrus (IFG) and anterior cingulate cortex (ACC)/pre-supplementary motor area (preSMA), and the Frontoparietal Network (FPN), anchored in the middle frontal gyrus (MFG) and posterior parietal cortex (PPC) ^34-40^. While this co-activation pattern is robust across development, prior longitudinal research has largely focused on structural markers ^7,21,41-44^ or trial-averaged functional activation ^45-48^.

Because trial-averaged mapping discards trial-to-trial neural variability, it obscures the dynamic reliability and precision of single-trial neural representations, which may serve as a much more sensitive index of functional network maturation ^48-54^. Beyond these task-positive systems, the default mode network (DMN) has also been implicated in attentional lapses, as a failure to suppress DMN activity during demanding tasks is associated with momentary inattention and inhibitory failures ^55^. Together, the SN, FPN, and DMN form a triple-network model that has been linked to cognitive control deficits across a wide range of psychopathologies ^56^; however, whether and how the coordinated maturation of these networks supports the protracted development of inhibitory control remains unknown.

To address these gaps, we leveraged large-scale longitudinal fMRI data from two complementary cohorts spanning a critical 13-year neurocognitive window (**Fig. 1a**): the Adolescent Brain Cognitive Development (ABCD) study (Release 4.0) ^57,58^, covering late childhood to early adolescence (ages 9–12; N = 1,928), and the IMAGEN study ^59,60^, extending from mid-adolescence to early adulthood (ages 14–22; N = 1,247). In both datasets, participants performed the SST, and we combined Single-Trial Modeling (STM) with Representational Similarity Analysis (RSA) to capture the spatiotemporal dynamics of neural representations underlying both reactive and proactive control (**Fig. 1b-f**), as well as behavioral stability indexed by intra-individual response variability (IIRV), a sensitive marker of attention and cognitive control efficiency ^51,61-66^.

**Fig. 1.**
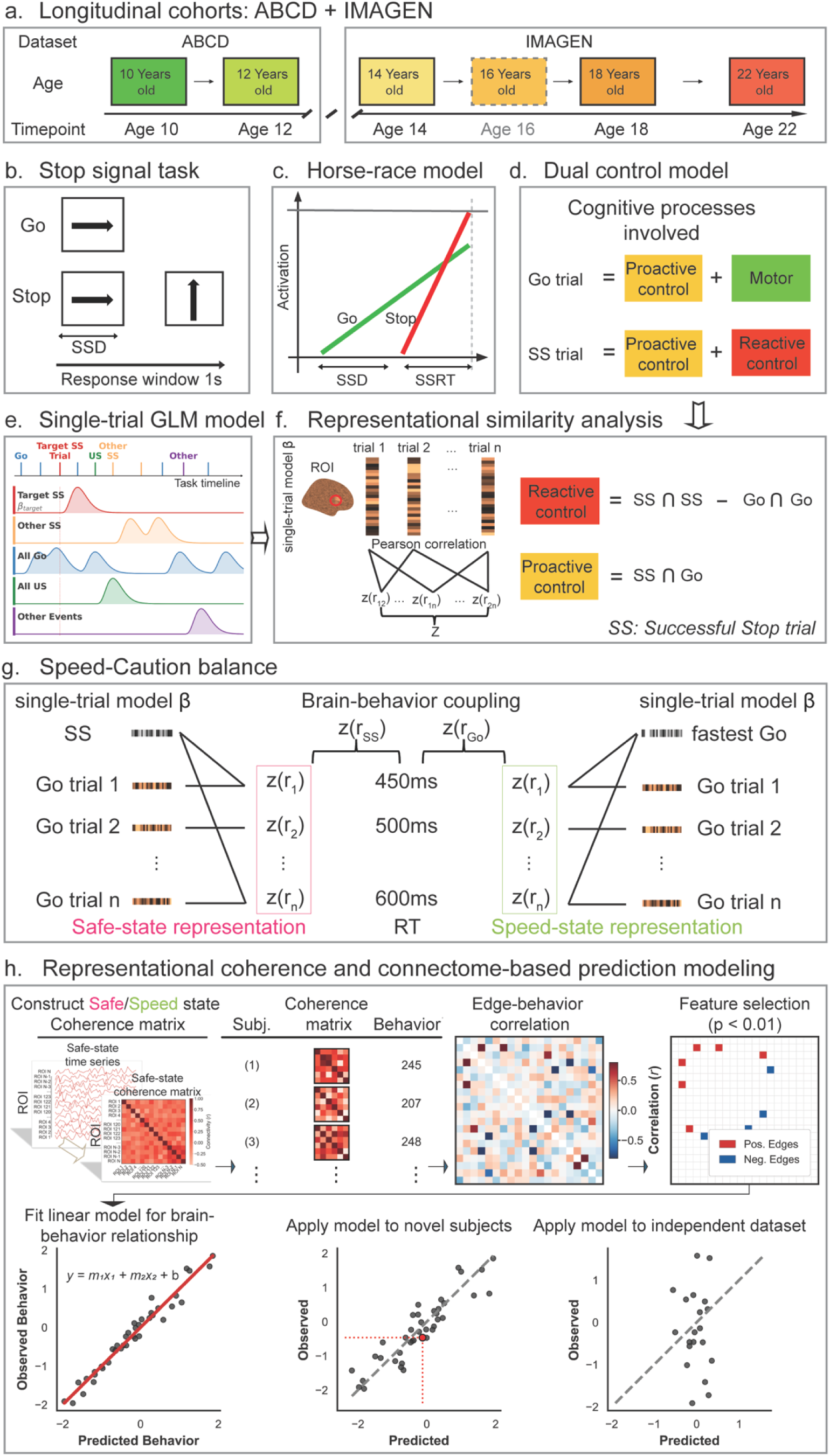
Schematic illustration of study design and analysis. **a.** Datasets. Two longitudinal cohorts spanning late childhood through early adulthood (ages 9–22) were analyzed: ABCD Release 4.0, capturing late childhood to early adolescence (Age 10, 9–10; Age 12, 11–12), and IMAGEN, extending coverage from mid-adolescence to young adulthood (Age 14; Age 18; Age 22; no neuroimaging at Age 16). b. Cognitive task. c. Horse-race model. Speed- and Safe-state compete and determine participants withhold their responses or not during the infrequent Stop-trials. d. Dual control model. Cognitive control operates in two modes: proactive and reactive, Go trials involving motor and proactive control process, Successful Stop (SS) trials involving both reactive and proactive control components. e. Single-trial modeling. The least-square single model includes one single-regressor for the target trials, other trials from the target trial condition were included as an additional regressor, other conditions were included as additional regressors. This single trial modeling procedure was applied to every trial iteratively. f. Representation similarity analysis. Pattern similarity was computed between the spatial activation pattern between the pair of trials of interest. Reactive control was measured by the within condition pattern similarity contrast between Successful Stop (SS) and Go trials. Proactive control was measured by the pattern similarity between SS and Go trials. g. Speed-caution balance. Trial-wise activity patterns were first averaged separately for the fastest Go trials (top 10%) and for SS trials to define two latent control templates. For each Go trial, pattern similarity to these templates yielded two runner time series: the Speed-state (indexing execution drive expressed on Go trials) and the Safe-state (indexing inhibitory/caution-related state expressed during stopping). Brain–behavior coupling was then quantified by relating each representation time series to Go-trial reaction time (RT), testing how trial-to-trial fluctuations in execution and caution predict behavioral speed. h. Representational coherence and connectome-based prediction modeling. We computed Safe- and Speed-state representation time series for each ROI, then constructed their specific representational coherence matrix by correlating the time series between every pair of ROIs. The resulting connectivity features were entered into connectome-based predictive models to predict SSRT and IIRV, evaluated via within-wave cross-validation and between-wave (and cross-dataset) generalization.

We structured our investigation across hierarchically organized levels of analysis anchored in the DCC framework ^23,24,67,68^. First, we assessed population-level representational stability, yielding Neural Stability Indices (NSI), to quantify the trial-to-trial consistency of reactive stopping versus proactive anticipatory networks (**Fig. 1c, d, f**). We hypothesized that reactive and proactive stability follow dissociable developmental trajectories. Second, we probed trial-level proactive dynamics by modeling moment-to-moment fluctuations between a “Speeded state” (maximal execution drive) and a “Safe state” (heightened proactive engagement). We hypothesized that Safe state processes would strengthen with age, serving as a robust, trait-like predictor of inhibitory control capacity. Finally, to resolve how distributed networks coordinate these states, we developed a novel “representational coherence” metric that captures whether distinct brain regions are co-committed to the same proactive mode. We hypothesized that Safe state representational coherence would require greater brain integration, maturing progressively to serve as a generalizable predictor of inhibitory control and psychopathology from late childhood into early adulthood.

## Results

### Cohort demographics and samples: ABCD and IMAGEN

We analyzed large-scale longitudinal fMRI data from two independent developmental cohorts: ABCD and IMAGEN. The ABCD dataset (Release 4.0) captured the transition from late childhood to early adolescence (Ages 10 and 12) ^57,58^. The IMAGEN dataset mapped mid-adolescence to young adulthood across three waves (Ages 14, 18, and 22) ^59,60^. Detailed screening criteria, demographic data, rigorous quality control criteria, and complete-case sensitivity analyses are provided in **Tables 1/2** and **Supplementary Methods**.

To ensure the data is not misinterpreted as a single continuous developmental timeline, results from these two cohorts are presented in separate sections.

### Development of inhibitory control: Late Childhood to Early Adolescence (ABCD)

We analyzed behavioral data on the SST to map the developmental trajectory of Inhibitory control function, quantified by the Stop-Signal Reaction Time (SSRT).

In the ABCD cohort, SSRT decreased significantly from Ages 10 to 12 (p<0.001), indicating rapid improvement in inhibitory control (**Fig. 2a** and **Table 1**).

**Fig. 2.**
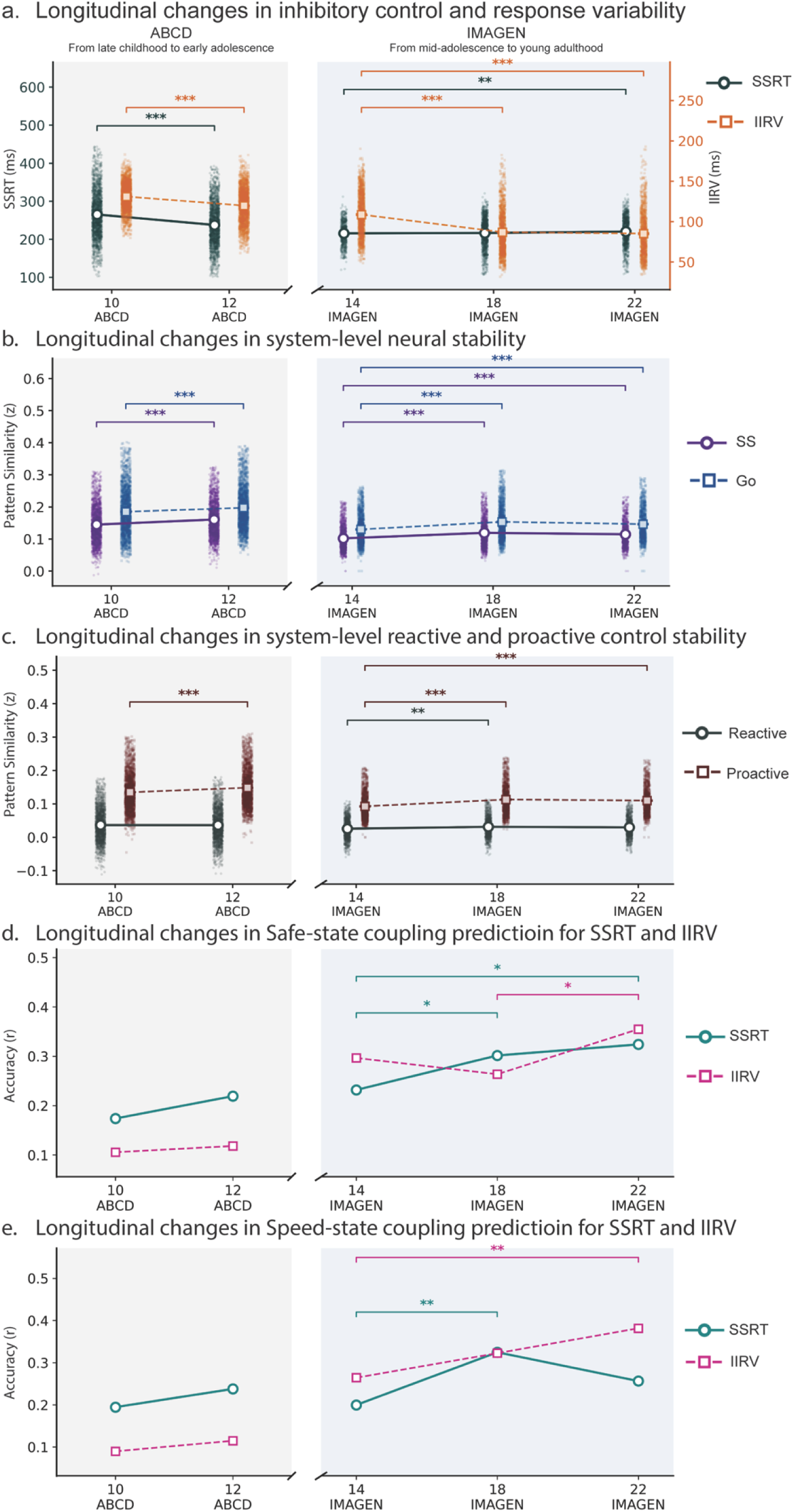
Longitudinal development in reactive suppression and intra-individual response variability (IIRV). **a.** Reactive suppression measured by stop-signal reactive time (SSRT) became faster from late childhood to early adolescence then slowdown from adolescence to early adulthood. b. System-level (whole-brain excluding sensorimotor networks) neural stability per trial type. Neural stability in SS trials and Go trials increased from late childhood to early adolescence and then increased from adolescence to early adulthood. c. System-level neural stability for reactive control and proactive control were computed using a whole-brain template excluding sensorimotor networks, reactive control remained stable from late childhood to early adolescence and then increased from adolescence to early adulthood. Proactive control increased from late childhood to early adulthood. d. Safe-state brain–behavior coupling for SSRT prediction increased continuously from late childhood through adolescence to early adulthood. In contrast, Safe-state brain-behavior coupling for IIRV prediction exhibited a non-linear trajectory, showing a modest increase from late childhood to early adolescence and then remaining relatively stable through early adulthood. e. Speed-state brain–behavior coupling (prediction accuracy) for SSRT increased from late childhood to early adolescence, rose further from mid-adolescence (≈14 years) to late-adolescence (≈18 years), then plateaued and declined by early adulthood (≈22 years). Speed-state brain-behavior coupling for IIRV prediction increased approximately linearly from late childhood through adolescence to early adulthood.

**Table 1.** Demographics and behavioral results (ABCD)

| **Demographics** | **Age 10 (1928)** | **Age 12 (1928)** | **p-value** |
| --- | --- | --- | --- |
| Age (SD) | 9.998 (0.621) | 12.047 (0.655) | 0 |
| Gender (Female/Male) | 967/961 | 967/961 |  |
| Head Motion (SD) | 0.210 (0.089) | 0.189 (0.081) | p<0.001 |
| **SST Performance** |  |  |  |
| Go Accuracy (%) | 91.4(0.050) | 93.2(0.048) | p<0.001 |
| Stop Accuracy (%) | 53.4(0.026) | 52.5(0.028) | p<0.001 |
| SSRT (ms) | 264.783 (60.875) | 236.983(54.099) | p<0.001 |
| Go RT (ms) | 529.943(75.154) | 468.336(74.514) | p<0.001 |
| Go RT STD (ms) | 131.059(16.861) | 119.810(20.935) | p<0.001 |

### Development of inhibitory control: Mid-Adolescence to Young Adulthood (IMAGEN)

In the IMAGEN cohort, SSRT trajectories were non-linear, as stopping speed did not significantly improve between Age 14 and 18, and it was significantly slower by Age 22 (p<0.05, **Fig. 2a** and **Table 2**).

**Table 2.** Demographic and behavioral results (IMAGEN behavioral sample).

| **Demographics** | **Age 14 (605)** | **Age 18 (1169)** | **Age 22 (997)** | **p-value** |
| --- | --- | --- | --- | --- |
| Age (SD) | 13.94 (0.74) | 18.02 (2.07) | 21.90(1.92) | 0 |
| Gender (Female/Male) | 331/274 | 614/555 | 525/472 | p=0.650 |
| **SST Performance** |  |  |  |  |
| Go Accuracy (%) | 93.86(4.28) | 94.08(4.56) | 93.89(4.81) | p=0.296 |
| Stop Accuracy (%) | 48.00(1.70) | 50.13(1.48) | 50.13(1.51) | p<0.001 |
| SSRT (ms) | 215.76 (30.82) | 216.64(38.92) | 220.23(37.54) | p=0.001 |
| Go RT (ms) | 416.78(54.27) | 388.96(55.58) | 393.39(61.98) | p<0.001 |
| IIRV (ms) | 96.57(22.11) | 85.58(24.81) | 83.97(26.56) | p<0.001 |

Behavioral measures included Go accuracy, Stop accuracy, Go reaction time (GoRT), , stop-signal reaction time (SSRT), and intra-individual response variability (IIRV). Linear mixed model was used to test the main effect of development on behavioral performance and head motion; p-values are reported. Sex distribution differences across visits were tested with chi-square tests.

### Development of behavioral stability: Late Childhood to Early Adolescence (ABCD)

Next, we examined the developmental trajectory of behavioral stability, operationalized as Intra-Individual Response Variability (IIRV), quantified by the standard deviation of reaction times across all correct Go trials.

In the ABCD study, IIRV decreased significantly from ages 10 to 12 (p < 0.001), indicating a rapid growth of behavioral stability (**Fig. 2a** and **Table 1**).

### Developmental trajectory of behavioral stability: *Mid-Adolescence to Young Adulthood (IMAGEN)*

In the IMAGEN cohort, IIRV at Age 22 was lower than at Age 18 (p = 0.029) and Age 14 (p < 0.001), and that Age 18 was also lower than Age 14 (p < 0.001; all p values FDR-corrected; see **Fig. 2a** and **Table 2**), suggesting that IIRV improved continuously from mid-adolescence through early adulthood.

### System-level neural stability increases with development: *Late Childhood to Early Adolescence (ABCD)*

The continuous improvement in behavioral stability suggests ongoing maturation of cognitive control systems from late childhood through early adulthood. We hypothesized that task-evoked activation within the cognitive control system becomes increasingly stable across this developmental period. Using Single-Trial Modeling (STM) and Representational Similarity Analysis (RSA) ^65,66^ (see **Supplementary Method**), we calculated a system-level neural stability index to capture the spatial consistency of task-evoked activation across trials.

In the ABCD cohort, neural stability for both Successful Stop (SS) and Go trials increased significantly from Age 10 to Age 12 (both ps < 0.001, FDR corrected, **Fig. 2b**).

### System-level neural stability increases with development: *Mid-Adolescence to Young Adulthood (IMAGEN)*

In the IMAGEN cohort, neural stability increased robustly from Age 14 to Age 18 (all *ps* < 0.001; **Fig. 2b**), before shifting to a plateau or modest decrease by Age 22 (see details in **Supplementary Results**).

### Neural Stability Trajectories of Reactive and Proactive Control: *Late Childhood to Early Adolescence (ABCD)*

Next, we examined the developmental trajectory of reactive and proactive control using system-level neural stability indices. Drawing on the dynamic DCC framework, reactive control is specifically isolated by contrasting SS with Go. Therefore, we define Reactive NSI as the difference in within-condition spatial pattern similarity between SS∩SS and Go∩Go (NSI_reactive_ = Similarity_SS∩SS_ - Similarity_Go∩Go_). Conversely, because proactive control processes are broadly engaged in anticipation of interference during both trial types, we define Proactive NSI based on the spatial pattern similarity between SS and Go trials (NSI_proactive_ = Similarity_SS∩Go_).

In the ABCD cohort, the Reactive NSI showed no significant change during this period (p = 0.237, FDR corrected). In contrast, the Proactive NSI increased significantly over this period (p < 0.001, FDR corrected; **Fig. 2c**).

### Neural Stability Trajectories of Reactive and Proactive Control: *Mid-Adolescence to Young Adulthood (IMAGEN)*

In the IMAGEN cohort, the Reactive NSI increased robustly during mid-adolescence (Age 14 to Age 18) (*p* < 0.001, FDR-corrected) but showed no further significant change from Age 18 to Age 22 (*p* = 0.163, FDR-corrected). Similarly, the Proactive NSI increased significantly from Age 14 to Age 18 (*p* < 0.001, FDR-corrected) and remained stable thereafter (*p* = 0.219, FDR-corrected; **Fig. 2c**). These system-level indices reveal that the neural stability of both control mechanisms increases during mid-adolescence, and plateaus upon entry into adulthood.

### System-level neural stability predicts inhibitory control development: *Late Childhood to Early Adolescence* (ABCD)

Next, we tested whether individual differences in the longitudinal changes of neural stability track with behavioral inhibitory control development. We computed partial Pearson’s correlations between longitudinal changes in the Reactive or Proactive NSI and changes in SSRT, while controlling for age, sex, head motion, and scanner site.

In the ABCD cohort, increases in the Reactive NSI were negatively associated with changes in SSRT (r = −0.078, p < 0.001), whereas changes in the Proactive NSI were not significantly related to SSRT changes (p = 0.651).

### System-level neural stability predicts inhibitory control development: *Mid-Adolescence to Young Adulthood (IMAGEN)*

In the IMAGEN cohort, change in the Reactive NSI exhibited a similar negative relationship with SSRT change during the later transition to young adulthood (Age 14 to Age 22: r = −0.081, p = 0.047; Age 18 to Age 22: r = −0.095, p = 0.047; FDR corrected), though not during mid-adolescence (Age 14 to Age 18: p = 0.144). Conversely, changes in the Proactive NSI were positively associated with SSRT change during mid-adolescence (Age 14 to Age 18: r = 0.109, p = 0.014; FDR corrected), but not in subsequent periods.

### Network-level neural stability of reactive and proactive control and its developmental trajectory: *Late Childhood to Early Adolescence (ABCD)*

To spatially localize the developmental changes in reactive and proactive control stability, we performed a whole-brain searchlight analysis (see **Supplementary Results and Fig. S1-4** for voxel-wise, condition-specific neural stability maps). We extracted spatial stability indices from predefined brain masks to determine whether the observed developmental effects were consistently expressed within core cognitive control systems (SN, FPN, and DMN).

In the ABCD cohort, we did not find significant developmental changes in reactive control representational patterns across core systems (**Fig. 3a**), but there was significant developmental strengthening in proactive stability within the FPN and DMN (p<0.05, FDR corrected, **Fig. 3b**).

**Fig. 3.**
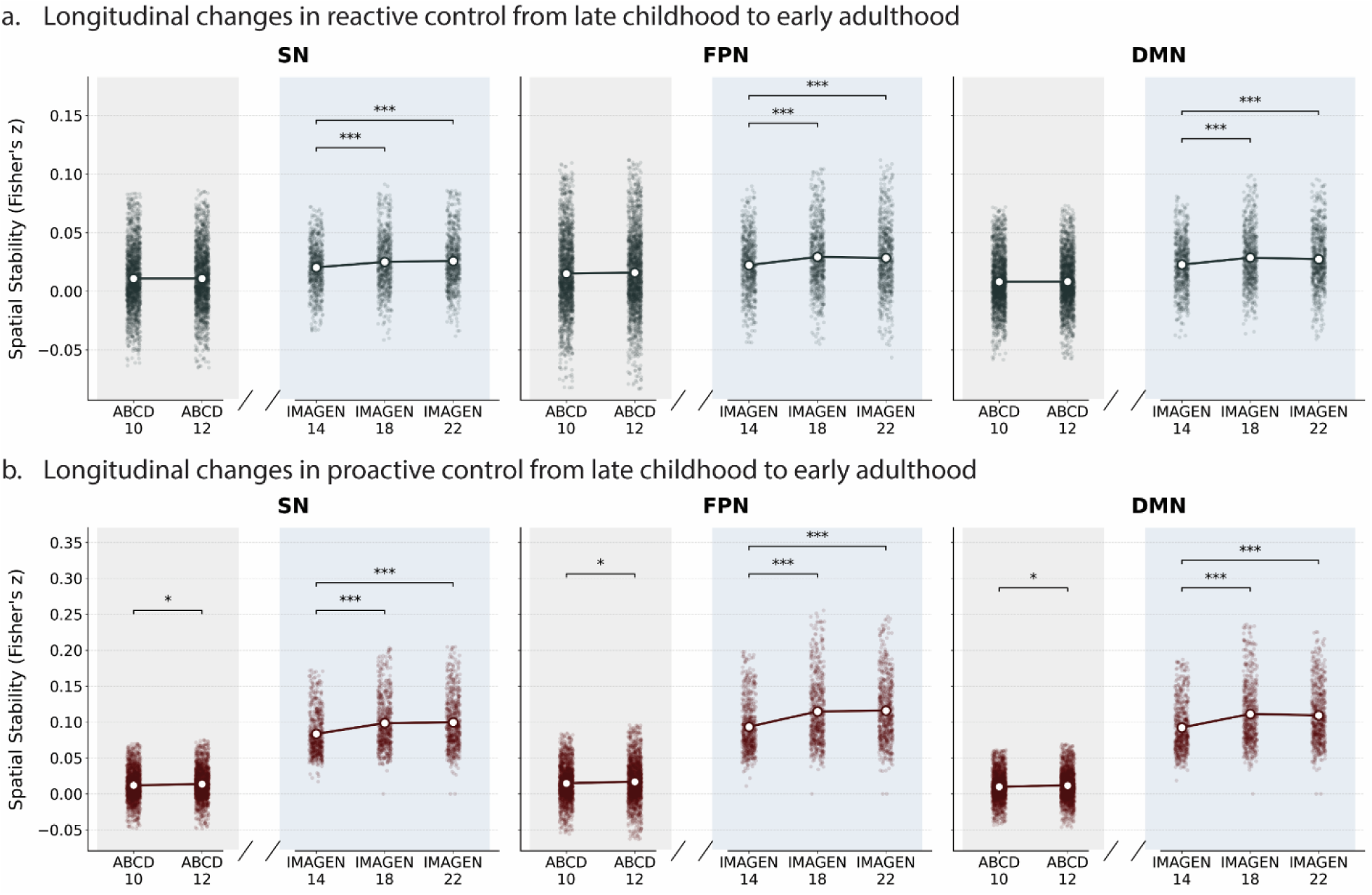
Longitudinal development in reactive and proactive control. **a.** Longitudinal changes in reactive control in the SN-FPN-DMN cognitive control system from late childhood to early adulthood. Spatial stability of reactive control remained stable from late childhood to early adolescence (10–12 years). Across all three networks, spatial stability significantly increased from mid-adolescence (≈14 years) to 18 years, followed by a plateau with no significant differences observed between 18 and early adulthood (≈22 years). Age-related developmental effects were analyzed using paired t-tests for the ABCD cohort and linear mixed-effects models for the IMAGEN cohort. b. Longitudinal changes in proactive control from late childhood to early adulthood. Spatial stability supporting proactive control increased linearly from late childhood through adolescence, plateaued around late adolescence (≈18 years), and then declined by early adulthood (≈22 years). Age-related developmental effects were analyzed using paired t-tests for the ABCD cohort and linear mixed-effects models for the IMAGEN cohort. Post-hoc pairwise comparisons were corrected using the False Discovery Rate (FDR) procedure. p < 0.05, *p < 0.01, **p < 0.001.

### Network-level neural stability of reactive and proactive control and its developmental trajectory: *Mid-Adolescence to Young Adulthood (IMAGEN)*

In the IMAGEN cohort, we observed a dynamic trajectory of strengthening followed by stabilization (see **Supplementary Results** for voxel-wise, condition-specific neural stability maps). There were significant, continuous improvements in spatial stability of reactive control across all three predefined networks (SN, FPN, and DMN). Neural stability increased continuously from Age 14 to both Age 18 and Age 22 (ps < 0.001, **Fig. 3a**).

Proactive stability increased across all three networks from Age 14 to 18. Neural stability increased significantly from Age 14 to Age 18 and Age 22 across all predefined networks (SN, FPN, DMN; ps < 0.001, **Fig. 3b**). Furthermore, stability peaked at Age 18 for the FPN and DMN, remaining significantly higher than at Age 22 (p = 0.026 and p = 0.044, respectively).

### Neural substrates of dynamic speed–caution decision-making predict trial-wise RT: *Late Childhood to Early Adolescence (ABCD)*

The dynamic DCC model posits that participants modulate their response strategies on a trial-by-trial basis, fluctuating between speed and caution. During the SST, these response tendencies are adjusted based on expectation: a bias toward speed facilitates rapid execution, whereas a bias toward safety results in strategic, cautious slowing. To characterize the neural dynamics of these opposing decision processes, we developed two distinct neural representation templates. A Speed-state template was defined as the average activation pattern from the fastest 10% of Go trials (indexing rapid action execution), while a Safe-state template was derived from SS trials (indexing inhibitory control). We then computed the representational similarity of each individual Go trial to both templates. We hypothesized that moment-to-moment fluctuations in neural state expression would track behavioral variability: greater similarity to the Speed-state template should reflect a speed bias (faster RT), whereas greater similarity to the Safe-state template should reflect a caution bias (slower RT).

In the ABCD cohort, greater similarity to the Safe-state template showed significant positive correlations with RT, particularly in the SN, FPN, and motor areas (**Fig. S5a**), whereas greater similarity to the Speed-state template exhibited a widespread negative correlation with RT across FPN, DMN, and visual regions (**Fig. S5b**). Furthermore, the Safe-minus-Speed representation difference also predicted RT fluctuations (**Fig. S5c**), confirming that the trial-wise competition between these representations drives behavioral variability.

Longitudinal analyses revealed a progressive strengthening of brain-behavior coupling across all three networks for Safe-state representations, which favor caution (**Fig. 4c**, also see **Supplementary Results**).

**Fig. 4.**
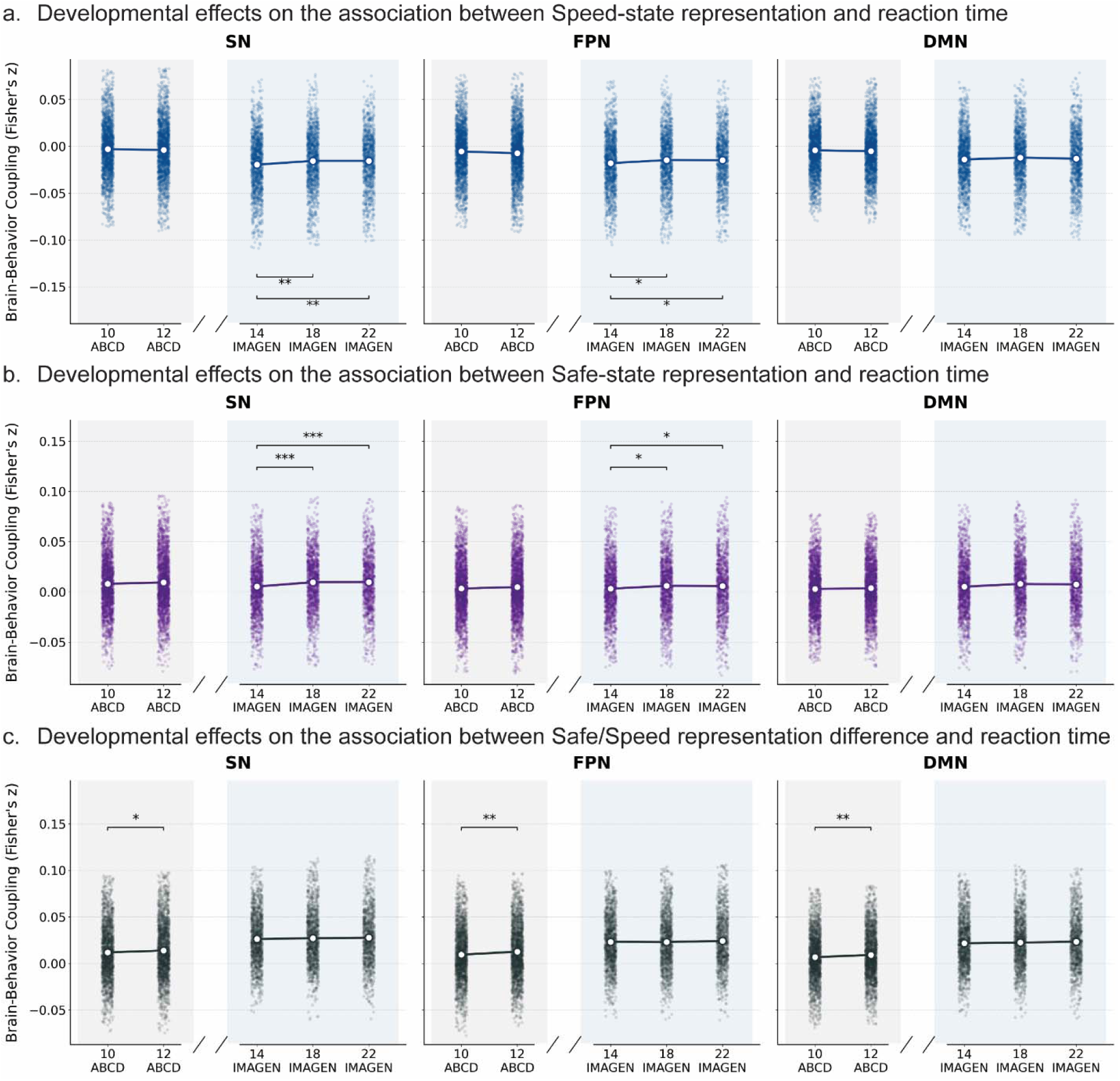
Longitudinal development in speed-caution balance from late childhood to early adulthood. **a.** The association between Speed-state representation and reaction time remained stable from late childhood to early adolescence. In both the SN and FPN, the magnitude of this negative association decreased from mid-adolescence (≈14 years) to 18 years, followed by a plateau with no significant changes through early adulthood (≈22 years). b. The association between Safe-state representation and reaction time was similarly stable from late childhood to early adolescence. In the SN and FPN, this association strengthened from mid-adolescence to 18 years, plateauing thereafter. c. The association between reaction time and the Safe-versus Speed-state representational difference significantly increased from late childhood to early adolescence across all three networks but showed no significant age-related changes from mid-adolescence to early adulthood. Age-related developmental effects were analyzed using paired t-tests for the ABCD cohort and linear mixed-effects models for the IMAGEN cohort. Post-hoc pairwise comparisons were corrected using the False Discovery Rate (FDR) procedure. p < 0.05, *p < 0.01, **p < 0.001.

### Neural substrates of dynamic speed–caution decision-making predict trial-wise RT: *Mid-Adolescence to Young Adulthood (IMAGEN)*

These findings were replicated in the IMAGEN cohort: the Speed-state consistently predicted faster RTs, while the Safe-state and the Competitive Difference score predicted slowing, with spatial patterns closely mirroring the ABCD cohort (**Fig. S5**).

Longitudinal trends paralleled those observed in childhood from the ABCD cohort, revealing a continuous strengthening of brain–behavior coupling for Safe-state representations in both SN and FPN (ps < 0.048) (**Fig. 4b**, **Supplementary Results and Fig. S6**).

### Brain-behavior coupling of Speed-state and Safe-state predict developmental change in SSRT and IIRV: *Late Childhood to Early Adolescence (ABCD)*

Next, we tested whether the strength of this coupling, governing the trade-off between fast and safe decision-making, predicts individual differences in cognitive control function (e.g., SSRT and IIRV). To do this, we constructed whole-brain ROI-level features for both Speed-state and Safe-state brain–behavior coupling using the Brainnetome atlas (excluding the sensorimotor system). We then trained ElasticNet regression models to predict SSRT and IIRV.

In the ABCD cohort, the Safe-state and Speed-state brain–behavior coupling robustly predicted individual differences in both SSRT and IIRV across Age 10 and Age 12 (ps < 0.001; **Fig. 5a-b**). SSRT prediction showed trend-level improvements from age 10 to 12 (Safe-state, p = 0.07; Speed-state, p = 0.078) (**Fig. 2d**). Importantly, longitudinal change in Safe-state coupling tracked behavioral maturation: changes in coupling strength predicted ΔSSRT (r = 0.13, p < 0.001) and ΔIIRV (r = 0.09, p < 0.001) (**Fig. 5c-d**). A similar pattern was observed for Speed-state coupling: changes in coupling strength predicted ΔSSRT (r = 0.11, p < 0.001) and ΔIIRV (r = 0.07, p = 0.004) (**Fig. 5c-d**).

**Fig. 5.**
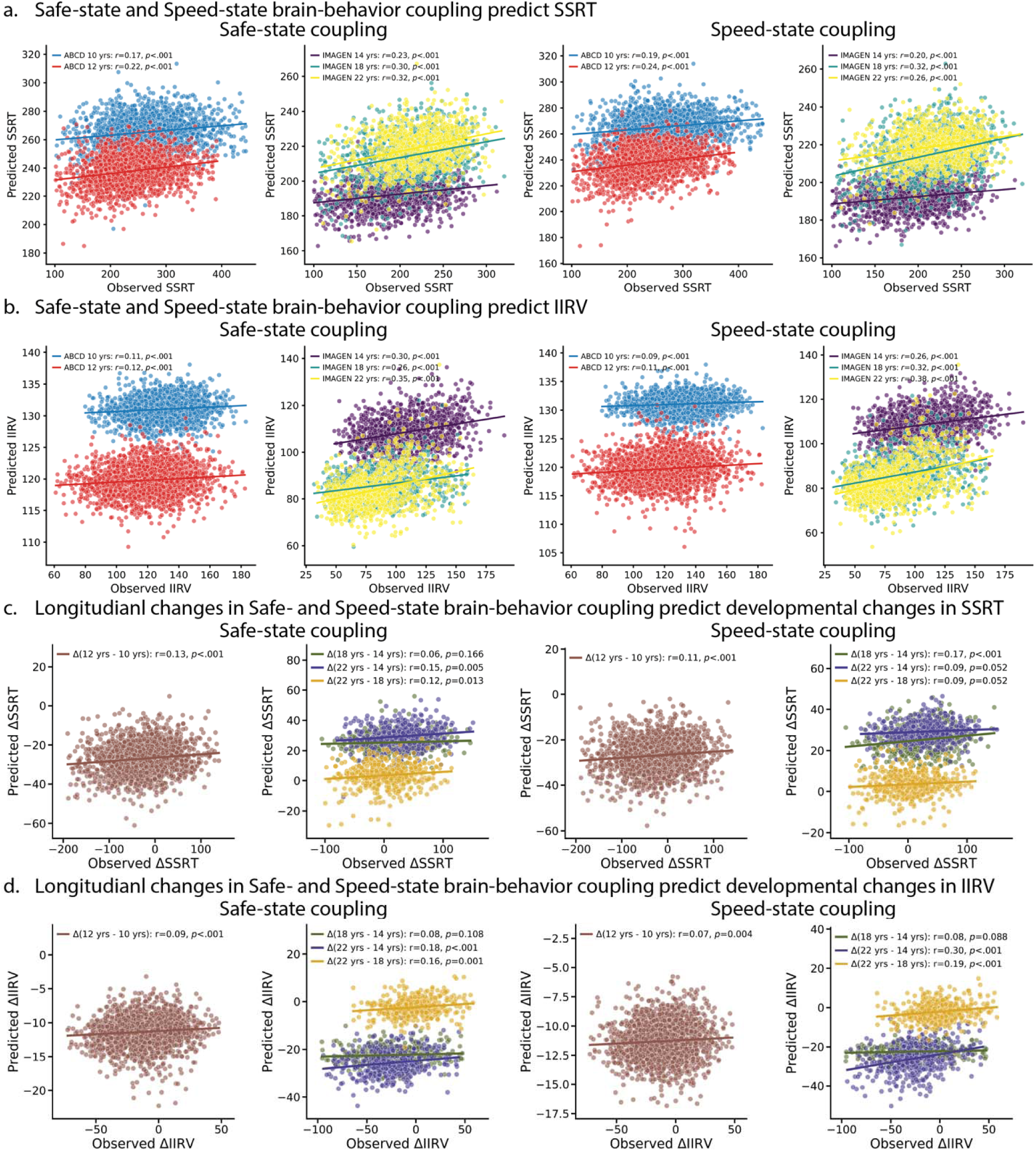
Neural substrates of speed-caution balance predict SSRT and IIRV in development. **a.** ElasticNet models trained on multi-regional Safe- or Speed-state brain-behavior coupling predict SSRT in development. b. Elastic models trained on multi-regional Safe- or Speed-state brain-behavior coupling predict IIRV in development. c. ElasticNet models trained on longitudinal changes in multi-regional Safe- or Speed-state brain-behavior coupling predicts developmental changes in SSRT. d. ElasticNet models trained on longitudinal changes in multi-regional Safe- or Speed-state brain-behavior coupling predicts developmental changes in IIRV.

### Brain-behavior coupling of Speed-state and Safe-state predict developmental change in SSRT and IIRV: *Mid-Adolescence to Early Adulthood (IMAGEN)*

Replicated in the IMAGEN cohort, both Safe-state and Speed-state coupling patterns robustly predicted SSRT and IIRV within each wave of the IMAGEN data (all ps < 0.001; **Fig. 5a-b**) as well as their longitudinal changes (all ps < 0.05; **Fig. 5c-d**). Prediction strength increased developmentally, with significant gains for SSRT (Speed-state: Age 14–18; Safe-state: Age 14–22) and IIRV (Safe-state: Age 18–22; Speed-state: Age 14–22) (all p < 0.05; **Fig. 2e**).

### Brain-behavior coupling of Speed-state and Safe-state predict developmental change in SSRT and IIRV: *Cross-wave and Cross-dataset generalization*

Elastic Net models trained within a given cohort/wave to predict SSRT and IIRV showed robust transfer across developmental waves and across cohorts (ps < 0.001, see **Supplementary Results and Fig. S10**).

### Brain-behavior coupling of Speed-state and Safe-state predict clinical symptoms: *Late Childhood to Early Adolescence (ABCD)*

Next, we investigated whether the brain-behavior coupling governing speed-safety trade-offs predicts psychopathology using Canonical Correlation Analysis (CCA).

In the ABCD cohort, Safe-state coupling changes tracked with anxiety, depression, and ADHD symptom trajectories. (r = 0.415, p = 0.003; **Fig. 6a**). No significant association was found for Speed-state (p = 0.280).

**Fig. 6.**
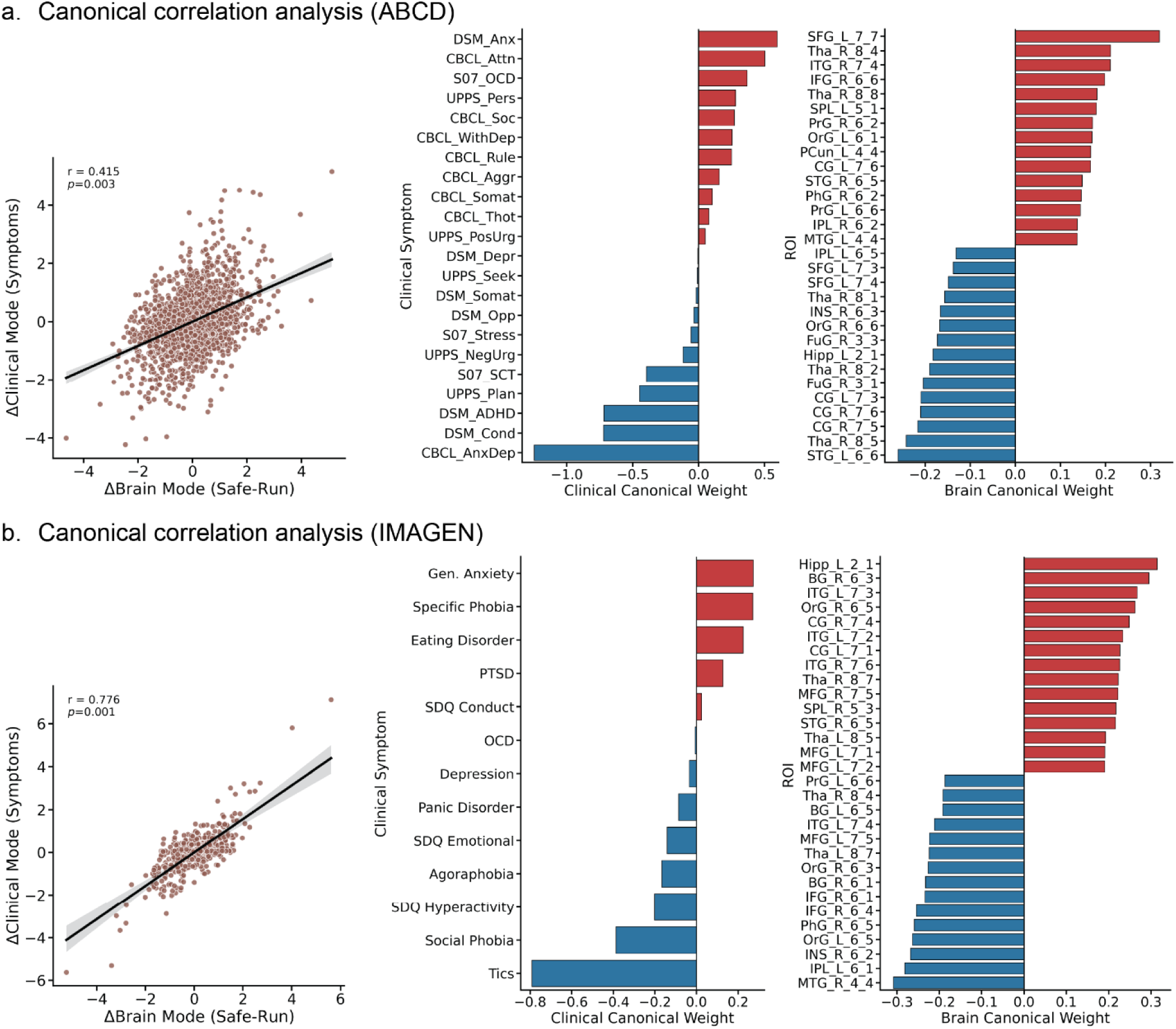
Canonical correlation analysis of the relation between longitudinal changes in Safe-state brain-behavior coupling and clinical symptoms. **a.** The longitudinal changes in multi-regional Safe-state brain-behavior coupling predict developmental changes in clinical symptoms from Age 10 to Age 12 in ABCD. The multivariate relationship was characterized by high canonical coefficients of right Insula, cingulate cortex, conduct and ADHD symptoms, and Planning function. b. The longitudinal changes in multi-regional Safe-state brain-behavior coupling predict developmental changes in clinical symptoms from Age 14 to Age 22 in IMAGEN using the ‘complete-case’ sample. The multivariate relationship was characterized by high canonical coefficients of right middle temporal gyrus (MTG), left inferior parietal lobe, and right insula, and Tics, Social Phobia, and Hyperactivity symptoms.

### Brain-behavior coupling of Speed-state and Safe-state predict clinical symptoms: *Mid-Adolescence to Early Adulthood (IMAGEN)*

Similarly, in the IMAGEN cohort, Safe-state coupling changes from Age 14 to Age 22 predicted shifts in tics, social phobia, and generalized anxiety (r = 0.776, p < 0.001, FWE corrected, **Fig. 6b**). Speed-state coupling again showed no significant effects (ps > 0.089).

### Representational coherence predicts inhibitory control and behavioral stability: *Late Childhood to Early Adolescence (ABCD)*

Next, we developed a representational coherence metric to quantify the trial-by-trial synchronization of specific proactive states across distributed brain regions (see Method). Specifically, we tested whether representational coherence strength of Speed-state and Safe-state networks could predict individual differences in inhibitory control (i.e., SSRT) and behavioral stability (i.e., IIRV), using the connectome-based predictive modeling (CPM) approach with 10-fold cross-validation.

In the ABCD study, both the Safe-state and Speed-state networks at Age 10 significantly predicted SSRT (Safe-state: r = 0.13, p < 0.001; Speed-state: r = 0.12, p < 0.001) and IIRV (Safe-state: r = 0.16, p < 0.001; Speed-state: r = 0.11, p < 0.001. **Fig. 7a**). These predictive models were successfully replicated at Age 12, yielding comparable predictive power for both SSRT (Safe-state: r = 0.14, p < 0.001; Speed-state: r = 0.14, p < 0.001) and behavioral stability (Safe-state: r = 0.17, p < 0.001; Speed-state: r = 0.14, p < 0.001. **Fig. 7a**).

**Fig. 7.**
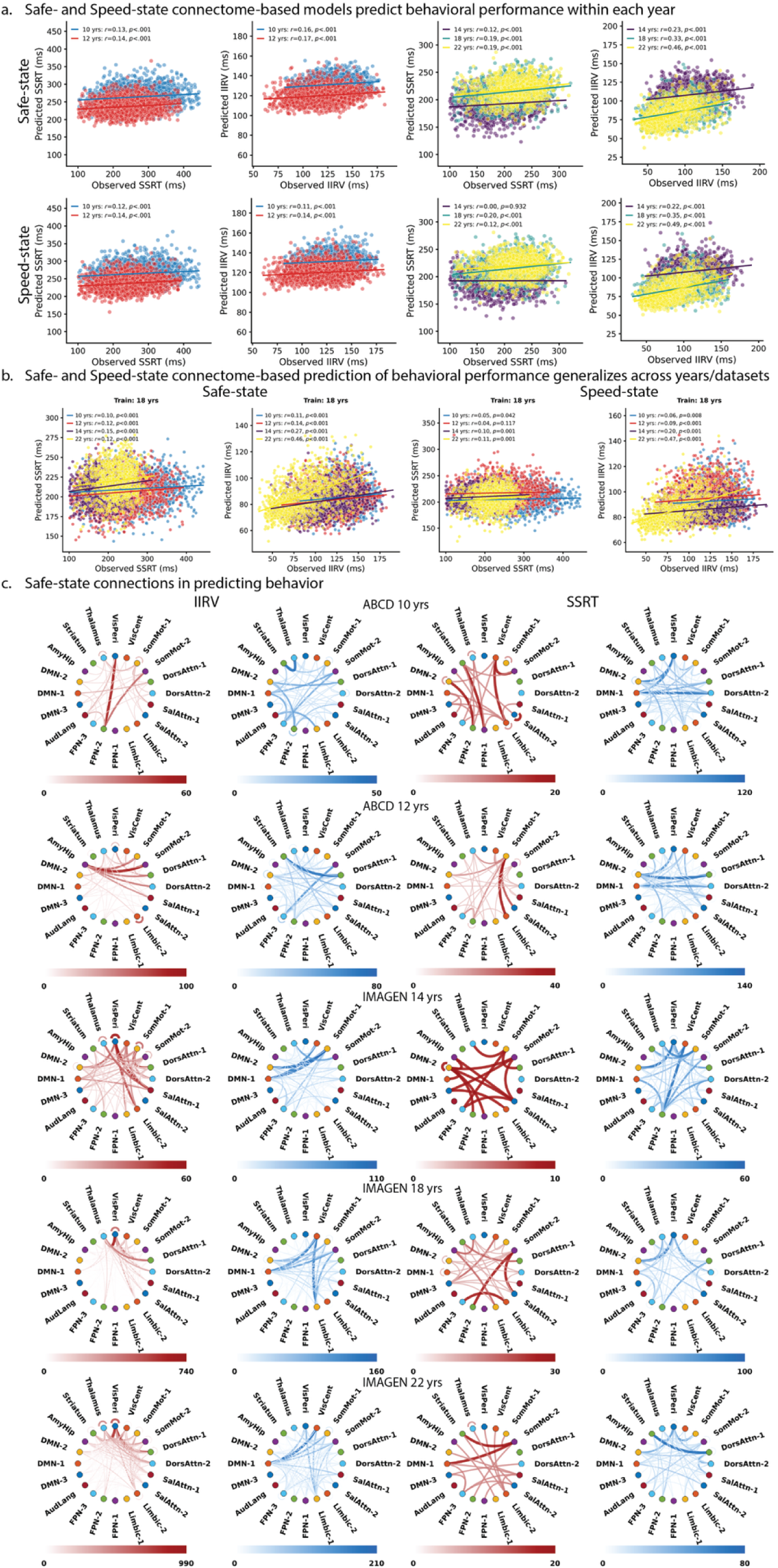
Safe-state and Speed-state connectome-based prediction models for behavioral development. **a.** Safe- and Speed-state connectome-based prediction models were able to predict individual differences in SSRT and IIRV within each wave. b. Connectome-based prediction models in predicting SSRT and IIRV were able to generalize across waves and cohorts. c. Consensus positive and negative connections of the prediction models within each wave. Individual differences in IIRV and SSRT were underlain by distinct connection patterns that also shifted across development. Stronger Safe-state connections among visual, sensorimotor, and dorsal and ventral attention networks reliably predicted greater IIRV, whereas stronger DMN–visual–sensorimotor connections predicted lower IIRV from mid-adolescence onward. Stronger connections involving the DMN, FPN, dorsal attention, and visual networks emerged as the most reliable predictors of faster SSRT.

### Representational coherence predicts inhibitory control and behavioral stability: *Mid-Adolescence to Young Adulthood (IMAGEN)*

The IMAGEN cohort replicated the results from the ABCD cohort (**Fig. 7a**). More strikingly, prediction strength increased with development: for IIRV, both Speed-state and Safe-state models showed significant increases from earlier waves to later waves (p < 0.003), whereas for SSRT, prediction increased for Speed-state from Age 14 to Age 18/22 (p < 0.01).

### Representational coherence predicts inhibitory control and behavioral stability: Generalizability of prediction models - *Late Childhood to Early Adolescence (ABCD)*

To evaluate the robustness and generalizability of our representational coherence models, we performed cross-wave prediction.

In the ABCD cohort, models trained at Age 10 successfully predicted future inhibitory control and behavioral stability at Age 12 (Safe-state: SSRT r=0.18, IIRV r=0.18; Speed-state: SSRT r=0.14, IIRV r=0.19; all p<0.001). Importantly, the reverse generalization (Train Age 12 -> Test Age 10) was equally significant (Safe-state: SSRT r=0.171, IIRV r=0.181; Speed-state: SSRT r=0.150, IIRV r=0.167; all p<0.001. See **Fig. S12-13**).

### Representational coherence predicts inhibitory control and behavioral stability: Generalizability of prediction models - *Mid-Adolescence to Young Adulthood (IMAGEN)*

Cross-wave generalizability was similarly robust in the longer-term IMAGEN cohort. Models trained at mid-adolescence (Age 14) successfully predicted outcomes at young adulthood (Age 18 and Age 22), with significant correlations ranging from r=0.10 to 0.30 (all p<0.001), with the exception of Speed-state predicting SSRT at Age 22 (p=0.071), see **Fig. S13**. Notably, models trained at Age 18 predicted Age 22 performance with high accuracy, particularly for behavioral stability (Safe-state r=0.46; Speed-state r=0.47; all p<0.001. See **Fig. 7b**).

### Representational coherence predicts inhibitory control and behavioral stability: Universal generalizability of behavioral stability

To rigorously evaluate the universality of our models, we also performed cross-cohort prediction and demonstrated remarkable cross-cohort robustness for the prediction of behavioral stability (IIRV) (**Fig. 7b**) and selective generalization of inhibitory control (**Fig. 7b**). see **Supplementary Results** and **Fig. S12-13** for details.

### Representational coherence predicts clinical symptoms: *Late Childhood to Early Adolescence (ABCD)*

Finally, we examined whether individual differences in representational coherence in the Speed-state and Safe-state states were related to psychopathology, using CCA.

In the ABCD cohort, multivariate clinical associations emerged significantly at Age 12 for both Safe-state (r = 0.437, p = 0.012) and Speed-state (r = 0.448, p = 0.008; all p values were FDR corrected, **Fig. 8a-b**). The Safe-state clinical profile was bidirectional, linking Stress, Thought Problems, and Oppositional behaviors against Aggression, Depression, and ADHD symptoms, driven neurally by Dorsal Attention-Limbic network connectivity. Conversely, the Speed-state profile contrasted Rule-breaking and Oppositional behaviors with Stress, associated with stronger Salience/Somatomotor-DMN connectivity and weaker inter-Frontoparietal Network (FPN-1 to FPN-2) coupling.

**Fig. 8.**
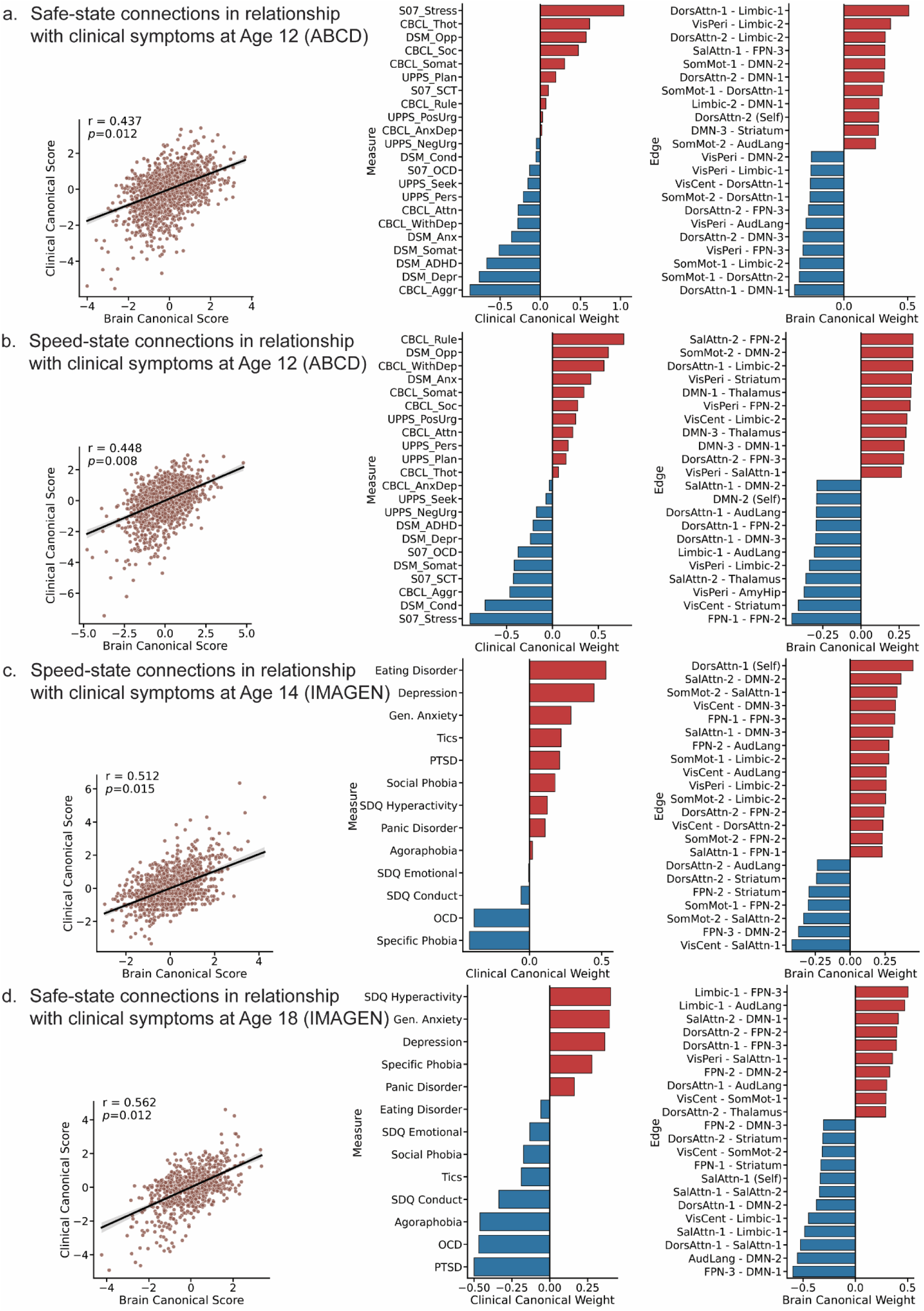
Representational coherence predicts clinical symptom profiles across development. Canonical correlation analysis (CCA) assessed multivariate associations between network-level representational coherence and clinical symptoms. (a) Safe-state coherence was significantly associated with clinical symptom profiles at ABCD Age 12 (canonical r = 0.437, p = 0.012, FDR-corrected). (b) Speed-state coherence was significantly associated with clinical symptom profiles at ABCD Age 12 (canonical r = 0.448, p = 0.008, FDR-corrected). (c) Speed-state coherence was significantly associated with clinical symptom profiles at IMAGEN Age 14 (canonical r = 0.512, p = 0.015, FDR-corrected). (d) Speed-state coherence was significantly associated with clinical symptom profiles at IMAGEN Age 18 (canonical r = 0.562, p = 0.012, FDR-corrected). For each panel, the middle plot shows canonical loadings for clinical measures, and the right plot shows the highest-contributing brain connections for the significant canonical mode.

### Representational coherence predicts clinical symptoms: *Mid-Adolescence to Young Adulthood (IMAGEN)*

In the IMAGEN cohort, Speed-state coherence linked to symptom dimensions at Age 14 (canonical r = 0.512, p = 0.015; **Fig. 8c**), revealing a bidirectional profile that contrasted positive (Hyperactivity, General Anxiety, Depression) and negative (PTSD, OCD, Agoraphobia) clinical loadings. Neurally, this mapped onto stronger Limbic-FPN and weaker FPN3-DMN1 connectivity. Conversely, Safe-state coherence showed significant association during young adulthood at Age 18 (canonical r = 0.562, p = 0.012; **Fig. 8d**), characterized by largely positive clinical loadings (Eating Disorders, Depression, Anxiety) driven by stronger Dorsal Attention and Salience-to-DMN connectivity.

## Discussion

Our study provides the most comprehensive longitudinal account to date of how the brain’s dual inhibitory control systems mature from late childhood to early adulthood. By combining single-trial modeling with representational similarity analysis across the ABCD (ages 9–12) and IMAGEN (ages 14–22) cohorts (**Fig. 1**), we bypass static, trial-averaged activation maps to capture the spatiotemporal dynamics of cognitive control. Our results reveal six convergent findings. First, inhibitory control, as indexed by SSRT, improves rapidly in childhood and plateaus in early adolescence, with no significant gains from mid-adolescence to young adulthood. In contrast, behavioral stability, indexed by IIRV, continues to improve from late childhood into early adolescence and again from mid-adolescence to young adulthood (**Fig. 2**). Second, these behavioral trajectories were mirrored by dissociable patterns of neural representations: neural stability increased globally, but reactive and proactive control networks exhibited distinct patterns of developmental change (**Fig. 2, 3**). Third, longitudinal changes in neural stability of reactive control predicted inhibitory control development. Fourth, trial-by-trial fluctuations between the Speeded and Safe states tracked behavioral variability across the full age range, and their coupling with behavior strengthened progressively with development (**Fig. 4**). Fifth, the strength of this state-behavior coupling predicted individual differences in inhibitory control and behavioral stability, with generalizability increasing across waves and independent datasets (**Fig. 5 and Fig. S10**). Finally, Safe state representational coherence, quantifying the degree to which distributed networks jointly and synchronously commit to an inhibitory control mode, emerged as a scanner-invariant, trait-like marker that generalized across cohorts to predict transdiagnostic psychopathology (**Fig. 8**). The ABCD and IMAGEN cohorts each provide internally longitudinal evidence within their respective age windows. Taken together, the findings from both cohorts establish a unified neurocognitive framework in which the progressive stabilization of proactive control networks and the maturation of speed-safety regulation together drive the transition to adult-like cognitive control.

### Development of inhibitory control and behavioral stability

Behaviorally, within the specific age span of the ABCD cohort (ages 9–12), inhibitory control (indexed by SSRT) improved significantly (**Fig. 2a**), replicating the well-established finding of rapid reactive stopping development during late childhood ^8,9,27,69,70^. Separately, in the older IMAGEN cohort (ages 14–22), SSRT was stable by mid-adolescence, with a modest slowing observed by age 22 (**Fig. 2a**), a pattern consistent with prior reports of SSRT plateauing across the adolescence-to-adulthood transition ^8,71-73^. In contrast to these distinct reactive trajectories, behavioral stability (indexed by IIRV) improved continuously within the respective timeframes of both independent datasets (**Fig. 2a**). This dissociation suggests that reactive stopping reaches a functional ceiling relatively early, while sustained proactive regulation continues to mature well into early adulthood ^74,75^.

### Dissociable neural stability maturation of reactive and proactive control

Our neural stability analyses mirrored these distinct behavioral trajectories within each separate cohort. In both studies, trial-to-trial representational fidelity increased broadly with age, regardless of trial type (**Fig. 2b**). No further change was observed after age 18 in the IMAGEN cohort, establishing a clear developmental plateau.

The Reactive Neural Stability Index (NSI), mainly localized across canonical stopping networks, showed no significant change in the ABCD cohort (ages 9–12), but increased robustly from age 14 to 18 in the IMAGEN cohort, followed by no further change between ages 18 and 22 (**Fig. 2c**).

The Proactive NSI, which localized globally across frontoparietal and default mode networks, showed significant increase from late childhood to early adolescence in the ABCD cohort (ages 9-12), as well as increase from early to middle adolescence and then stabilized in the IMAGEN cohort (age 14-22) (**Fig. 2c**).

Interestingly, longitudinal increases in Reactive, rather than Proactive NSI from late childhood to early adolescence predicted SSRT improvement in the ABCD cohort (ages 9-12), whereas increases in Proactive NSI during mid-adolescence were associated with SSRT change in the IMAGEN cohort (ages 14-22), highlighting a dynamic interplay where expanding proactive engagement modulates reactive stopping speed. Together, these parallel findings validate the DCC framework’s prediction that reactive and proactive control mature via distinct mechanisms and timelines ^23^.

### Trial-level dynamics and the Speed-Caution balance

By decoding moment-to-moment neural fluctuations along the proactive control continuum, we demonstrated that a speed-biased state (“Speed-state”) predicted faster reaction times, whereas a caution-biased state (“Safe-state”) predicted slower responses (**Fig. 4 and Fig. S5-9**). Notably, the Safe state, derived from Successful Stop trials, predicted response slowing even on Go trials across all developmental waves in both studies. This cross-trial generalization provides neural evidence for the anticipatory nature of proactive control: inhibitory representations are pre-activated to constrain execution speed even when stopping is not explicitly required ^48,76-79^. This also supports the view that proactive and reactive control recruit overlapping neural architectures ^48^, with the same circuits engaged tonically for proactive anticipation and phasically for reactive stopping.

Longitudinally, the progressive dominance of Safe over Speeded state coupling provides a neural signature of maturing cognitive regulation: not merely stronger inhibitory representations, but a tighter and increasingly dominant link between the proactive inhibitory state and its behavioral consequences. From late childhood through young adulthood, the brain–behavior coupling of the Safe state strengthened significantly in regions such as the bilateral anterior insula, pre-SMA, precuneus, and visual cortex (**Fig. S6-9**). This tightening relationship tracked behavioral maturation longitudinally within each separate cohort, establishing state-behavior coupling as a robust, trait-like neurobehavioral phenotype of cognitive control efficiency ^51^.

### Representational coherence: network-level co-commitment as a developmental marker

We introduced representational coherence to quantify whether distributed brain regions simultaneously shift toward the same cognitive state on a trial-by-trial basis. Safe state coherence revealed a globally distributed, highly integrated architecture, while Speeded state coherence was comparatively focal, restricted to sensorimotor and dorsal attention regions. This topological asymmetry, which was robust across trial subsets and replicated across all five waves in both cohorts, suggests that sustained, goal-directed regulation demands broader cross-network coordination than rapid action execution (see **Fig. S11**).

The predictive power of these coherence networks surged with age (**Fig. 7 and Fig. S12-13**). By early adulthood, models trained on Safe and Speeded state coherence achieved remarkable accuracy in predicting behavioral stability. Crucially, these predictive models demonstrated universal, bidirectional generalizability for IIRV across the ABCD and IMAGEN datasets.

Because these are independent datasets with different study designs and settings, this robust cross-cohort generalization confirms that sustained proactive regulation is instantiated in a scanner-invariant, trait-like network. SSRT prediction, by contrast, generalized more selectively across specific age windows, acting as a highly sensitive index of individual regulatory capacity.

### Triple-network coordination: integrating the default mode network into the maturation of inhibitory control

Our findings begin to resolve how the coordinated maturation of the triple network supports the protracted development of inhibitory control, extending the account beyond the task-positive salience and frontoparietal systems to the default mode network (DMN). Rather than maturation being confined to canonical control regions, the network-level stability of proactive control also strengthened within the DMN in late childhood, then increased across all three predefined networks (SN, FPN, and DMN) from mid-adolescence to early adulthood, peaking around Age 18 for the FPN and DMN before plateauing (**Fig. 3**). Also, the association between reaction time and the Safe-versus Speed-state representational difference significantly increased from late childhood to early adolescence across all three networks (**Fig. 4b**). Critically, our representational coherence metric, indexing the trial-by-trial synchronization of distributed regions to a common control state, predicted individual differences in behavioral stability and inhibitory control within and across cohorts and developmental waves (**Fig. 7**). The same coherence features were associated with a broad spectrum of transdiagnostic symptoms, with DMN couplings (e.g., salience/dorsal attention-to-DMN connectivity) among the contributing connections (**Fig. 8**). Together, these results indicate that maturing inhibitory control reflects not the isolated refinement of control regions but the progressive integration of the DMN with salience and frontoparietal systems, offering a mechanistic account of how triple-network coordination scaffolds cognitive control and why its disruption may confer transdiagnostic risk ^56^.

### Clinical relevance: safe state dynamics as a transdiagnostic marker

Inhibitory control deficits are among the most replicated transdiagnostic risk factors in psychiatry, cutting across externalizing disorders such as ADHD and conduct disorder, and internalizing disorders including anxiety and depression ^16-22^. A major contribution of this work is demonstrating that the neural phenotypes derived from both population and trial-level analyses, specifically those reflecting Safe state engagement, predicted transdiagnostic vulnerability across two independent cohorts using distinct clinical instruments.

Within the ABCD cohort, longitudinal changes in Safe state brain–behavior coupling predicted developmental changes in symptoms of anxiety, depression, and ADHD (**Fig. 6a**). Separately, in the IMAGEN cohort, Safe state coupling changes from age 14 to 22 strongly correlated with shifts in tics, social phobia, hyperactivity, and generalized anxiety (**Fig. 6b**). Across both independent groups, the right anterior insula emerged as a vital hub linking Safe state maturation to psychopathology ^56^.

Network-level representational coherence also tracked clinical symptoms, emerging significantly as children entered early adolescence (**Fig. 8**), perhaps reflecting the period when symptom profiles become more reliably expressed. The convergence of these state-behavior and coherence phenotypes across distinct study designs and clinical instruments positions Safe state representational coherence as a biologically grounded candidate biomarker for precision psychiatry.

The convergence of state-behavior coupling and representational coherence findings across two cohorts, two clinical instruments, and a broad spectrum of psychiatric dimensions suggests that the failure to consistently and efficiently engage the Safe state, whether indexed at the trial level (state-behavior coupling) or the network level (representational coherence), is a core, developmentally sensitive vulnerability factor that cuts across diagnostic boundaries. The selectivity of Safe over Speeded state associations is theoretically noteworthy: it is the proactive, anticipatory inhibitory regulatory system whose developmental maturation tracks psychiatric risk. This aligns with the DCC framework’s prediction that proactive control is the more cognitively demanding, frontally mediated component whose failure is most likely to manifest as clinical dysregulation across diagnostic categories ^23,25^. Crucially, the cross-dataset generalization of these coherence-based clinical predictions, across different scanners, continents, and age ranges, distinguishes Safe state representational coherence as a promising candidate biomarker for transdiagnostic risk stratification in precision psychiatry ^80-82^.

## Conclusion

This study establishes a unified neurocognitive framework for understanding how inhibitory control matures from childhood to adulthood. Across two large independent longitudinal cohorts we demonstrate that reactive and proactive control systems follow dissociable developmental timelines, with the progressive refinement of trial-level proactive dynamics: the strengthening Safe state coupling drives the emergence of behavioral stability and mature cognitive regulation. The Safe state representational coherence, quantifying whole-brain network co-commitment to an inhibitory control mode, emerged as a scanner-invariant, developmentally sensitive, transdiagnostic biomarker that generalizes across independent cohorts spanning different ages, scanners, and continents. These findings bridge developmental cognitive neuroscience and precision psychiatry, offering both a mechanistic account of cognitive control maturation and a validated neural phenotype for identifying transdiagnostic psychiatric risk across the critical childhood-to-adulthood transition.

## Methods

### Participant ABCD Dataset

Data for this study came from the baseline and two⍰year follow⍰up visits of the ABCD Study (ReleaseLJ4.0; CollectionLJ#2573). The ABCD study was conducted in accordance with institutional review board regulations at each site, including University of Maryland, Baltimore, University of Colorado, Boulder, University of Minnesota, the Laureate Institute for Brain Research, Oregon Health and Science University, University of Vermont, University of Pittsburgh, Virginia Commonwealth University, University of Rochester, University of Florida, Medical University of South Carolina, University of Michigan, University of Minnesota, University of Utah, SRI International, University of Wisconsin-Milwaukee, Children’s Hospital of Los Angeles, Florida International University, Washington University in St. Louis, and Yale University. Informed consent was obtained from parents and informed assent was obtained from child participants.

Of the full sample (N = 11817 at baseline; N = 7883 at two⍰year follow⍰up), we first applied the ABCD SST task⍰fMRI inclusion criteria (imgincl_sst_include == 1; N = 3546/1904 at baseline/two⍰year follow⍰up), required two SST runs of good quality (iqc_sst_total_ser = iqc_sst_good_ser = 2; N = 2784/1255), and retained only those with recorded scanner serial numbers (mri_info_deviceserialnumber; excluded 16/26). After this initial screening, 7594 participants remained at baseline and 5648 at two⍰year follow⍰up. Next, we excluded participants who: (1) did not have two SST runs in the minimally processed Release 4.0 data (N = 17/0); (2) had insufficient volumes to cover the full SST task (i.e., the last trial occurred more than two seconds after the final volume; N = 24/14); or (3) exhibited a mean framewise displacement greater than 0.5 mm in either run (as defined by Power et al.; N = 2138/1037). These exclusions resulted in 5456 participants at baseline and 4611 at follow-up.

Of these, 2671 participants had usable data at both timepoints and were subjected to behavioral quality screening. The following behavioral inclusion criteria were applied at both timepoints ^83^: (1) go-trial accuracy ≤ 75%; (2) stop-trial accuracy outside the 20–80% range; (3) mean stop-failure RT ≤ go-trial RT; (4) stop-signal reaction time (SSRT) ≤ 100 ms; and (5) to avoid familial clustering, we randomly retained one participant per family using genetic relatedness data (gen_y_pihat). The final analysis sample included 1928 participants.

### IMAGEN dataset

The IMAGEN study was approved by the local research ethics committees at each participating site, including King’s College London (London), University of Nottingham (Nottingham), Trinity College Dublin (Dublin), Central Institute of Mental Health (Mannheim), Technische Universität Dresden (Dresden), Charité – Universitätsmedizin Berlin (Berlin), Commissariat à l’Énergie Atomique et aux Énergies Alternatives (Paris), and University Medical Center at the University of Hamburg (Hamburg). Written informed consent was obtained from all participants and from a parent or legal guardian.

Participants underwent the same behavioral and fMRI quality screening procedures as described for the ABCD dataset. Each participant completed only one stop-signal task run per visit, in contrast to the ABCD dataset, which required both runs to meet quality criteria. To maximize the sample size for each visit and allow comparisons between any two visits, participants were included if they had at least one usable visit, rather than requiring complete data across all visits. From the full sample, there were 2116, 1403, and 2315 participants at baseline (mean ages 14, hereafter Age 14), first follow-up (mean ages 18, hereafter Age 18), and second follow-up (mean ages 22, hereafter Age 22), respectively.

At each visit, participants were excluded if they met any of the following criteria ^83^: (1) go-trial accuracy ≤ 75%; (2) stop-trial accuracy outside the 20–80% range; (3) mean stop-failure RT ≤ go-trial RT; (4) stop-signal reaction time (SSRT) ≤ 100 ms; (5) mean framewise displacement > 0.5 mm. After applying these behavioral and motion criteria, 1247, 1151, and 973 participants remained for Age 14, Age 18, and Age 22, respectively. Overlap across waves was 710 participants shared between Age 14 and Age 18, 600 between Age 14 and Age 22, 716 between Age 18 and Age 22, and 464 participants present in all three waves.

### Stop-signal task (SST)

In both datasets, participants performed a two-choice stop-signal task (SST) during fMRI to assess response inhibition (**Fig. 1b**). On each trial, a left- or right-pointing arrow prompted a speeded button press with the corresponding hand (go trials). On a subset of trials, the arrow was followed unpredictably by an upward-pointing stop signal instructing participants to withhold their response (stop trials).

In the ABCD dataset, participants completed two runs of 180 trials each (150 go and 30 stop trials per run; 300 go and 60 stop total). The stop-signal delay (SSD) was initially set to 50 ms and adjusted dynamically in 50 ms increments to maintain approximately 50% successful inhibition. Each trial lasted 1,000 ms, stimulus onset asynchrony (SOA) ranged ∼1,700–3,000 ms. In the IMAGEN dataset, participants completed one run of 400 go and 80 stop trials at baseline (360 total trials in the follow-up waves). Stopping difficulty was manipulated via a tracking algorithm that varied SSD in 50 ms steps (initial delay = 250 ms) to target 50% inhibition success. Stimuli were presented for up to 1,000 ms on go trials, and for 0–900 ms on stop trials depending on SSD.

### Behavioral measures

Reactive control was measured by SSRT. First, we confirmed that behavioral data did not violate the main assumption of the Race Model, that the mean RT in Unsuccessful Stop (US) trials should be shorter than the mean RT in Go trials, Go and Stop accuracy were in reasonable range (see detailed information in the participant screening section). Then, SSRT was computed using the integration method based on the Race model ^29,83^: SSRT = T−mean SSD, where T is the cutoff point where the integral of the observed distribution of Go RT in the SST equals the probability of unsuccessful stopping.

Developmental effects on behavioral performance and demographic measures were analyzed separately for the two datasets. For the ABCD dataset, paired Student’s t-tests were conducted since each participant contributed data from two time points. For the IMAGEN dataset, linear mixed-effects (LME) models with random intercepts for participants were employed to accommodate incomplete longitudinal data and thereby maximize the sample size for both within-visit effects and between-visit comparisons. The fixed effect of interest was Visit (Age 14, Age 18, Age 22), and dependent variables included Go accuracy, Stop accuracy, Go reaction time (GoRT), stop-signal reaction time (SSRT), and mean framewise displacement. The statistical significance of the Visit effect was evaluated using F-tests derived from the ANOVA of the LME models. Sex distribution, as a categorical variable, was compared across visits using chi-square tests. Descriptive statistics (mean ± SD) are reported for all measures. As a robustness check, we repeated the analyses in the subset of participants with valid data at all three IMAGEN waves to confirm that results were consistent with the primary findings.

### Clinical Assessments

#### Adolescent Brain Cognitive Development (ABCD) Study

Clinical phenotyping in ABCD combined dimensional symptom assessment with trait-level self-regulation measures, drawing on parent-report questionnaires that are well validated for youth. Child Behavior Checklist (CBCL) ^84^. Parents completed the CBCL to capture broad variation in children’s emotional and behavioral functioning across multiple symptom domains (e.g., anxious/depressed symptoms, attention problems, aggressive and rule-breaking behaviors).

UPPS-P Impulsive Behavior Scale ^85,86^. Impulsivity was assessed using the 20-item UPPS-P for Children short form, which operationalizes impulsivity as five separable facets: Negative Urgency, Positive Urgency, (Lack of) Premeditation, (Lack of) Perseverance, and Sensation Seeking. These facet scores were used to quantify trait-like individual differences in self-regulatory tendencies that are conceptually complementary to symptom measures.

#### IMAGEN Study

In IMAGEN, psychiatric symptoms were characterized using structured, computer-assisted assessments designed to align with international diagnostic criteria (DSM-IV and ICD-10). Development and Well-Being Assessment (DAWBA) ^87^. DAWBA integrates adolescent- and parent-report interview data to generate disorder-specific probability bands (0–5) indexing the likelihood of meeting diagnostic criteria. In this study, we focused on the adolescent self-report DAWBA probability bands, which are consistently available across all three waves and are considered a reliable source for longitudinal symptom tracking. We included disorder with valid data across three waves, including specific phobia, social phobia, panic disorder, agoraphobia, post-traumatic stress disorder, obsessive–compulsive disorder, generalized anxiety, depression, eating disorders, and tic disorders.

Strengths and Difficulties Questionnaire (SDQ) ^88^. The self-report version of the Strengths and Difficulties Questionnaire (SDQ) was used in order to obtain dimensional assessments of psychopathology, including conduct problems, emotional problems, and hyperactivity problems.

#### Neuroimaging data

Imaging protocols for the ABCD and IMAGEN stop-signal task (SST) datasets followed previously published, multi-site acquisition schemes (see **Supplementary Methods** for details). To control potential scanner effects, MRI site was included as a nuisance covariate in all statistical analyses.

#### Preprocessing

Details of preprocessing steps were provided in **Supplementary Methods**.

#### Single-trial estimation

The GLMs were performed separately to estimate the activation pattern for each trial using a Least Square–Separate (LS-S) approach ^66,89^, in which the trial of interest was modelled as one regressor, while all remaining trials were grouped into separate regressors based on trial type (**Fig. 1e**). Details about model setting is provided in **Supplementary Methods**.

#### Representational similarity analysis

We utilized representational similarity analysis (RSA) employing both searchlight and regions of interest (ROI) methods to examine spatial stability during task performance ^48,65,90^. For searchlight analysis, β-maps were obtained from a cubic region of interest containing 125 surrounding voxels across each participant’s whole brain ^91-94^. To calculate pattern similarity, we correlated activity vectors of any given pair of trials using Pearson correlation. Subsequently, we transformed these similarity scores into Fisher’s z-scores and compared them between conditions to quantify proactive and reactive control. ROI-based analyses were conducted similarly, except that the Pearson correlation was computed using voxels specifically selected from the chosen ROI. To minimize autocorrelation confounds, for the ABCD dataset we excluded trial pairs from the same run. For IMAGEN, we anchored on Successful Stop (SS) trial pairs with a minimum temporal lag of 30 s. Matched pairs in other conditions (e.g., Go or SS–Go) were then selected to have a temporal lag within ±2 s of the reference SS pair. If no matched pair was available, the SS anchor pair was excluded. This procedure ensured that differences in pattern similarity between conditions were not driven by residual temporal autocorrelation and is consistent with previous within-run RSA analyses. In addition, behavioral analyses confirmed that the resulting contrasts did not differ significantly in temporal distance, ensuring that pattern similarity differences were not driven by temporal confounds.

Spatial stability of trial-evoked brain responses was assessed within Go trials and within Successful Stop (SS) trials, as well as between Go and SS trials for each participant. Reactive control was quantified by contrasting SS trials with Uncertain Go (US) trials, since only SS trials involve a reactive inhibitory process. Proactive control was indexed by the pattern similarity between Go and SS trials, reflecting the extent to which inhibitory control processes are pre-activated during Go trials. We hypothesized that greater similarity between Go and SS activation patterns would indicate stronger proactive control (**Fig. 1f**).

Because baseline from IMAGEN dataset included more trials (480) than the follow-up waves (360), trial pairs at Age 14 could span longer temporal lags and potentially bias pattern-similarity estimates despite our lag-matching procedure (see Methods). To rule out this confound, we conducted additional control analysis after matching baseline scan duration to Age18/Age22 and replicated findings reported in the main text (**Fig. S4**).

Finally, between-year comparisons were performed to test whether the spatial stability of proactive and reactive control changed with maturation. The resulting contrast maps were entered into group-level analyses. Statistical significance was evaluated using FSL’s PALM ^95,96^. For contrasts testing whether effects differed significantly from zero at each data point, scanner site was modeled with dummy-coded covariates for each visit. When examining developmental effects on contrasts of interest, additional covariates included age difference between visits, gender, and head motion.

#### Tracking RT fluctuations under a speed–caution balance framework

Go trials in the stop-signal task were conceptualized as reflecting a moment-to-moment speed–caution balance: an execution drive that favors rapid responding and a cautionary braking state that prepares the system to withhold a response when stopping may be required. We asked whether trial-wise expression of these two state representations predicts fluctuations in Go-trial reaction time (RT). To do so, we constructed two representational templates—a Speed-state (speed-biased) template and a Safe-state (caution-biased) template—and, for each Go trial, quantified the similarity between the ongoing brain pattern and each template. Trial-wise similarity values were then related to RT using whole-brain searchlight analysis (**Fig. 1g**). Safe-state–like representation (caution-biased): For each participant, we defined a Safe-state template based on successful stop trials. In the within-subject approach (reported in the Supplementary Materials), an inhibitory-control template was obtained by averaging single-trial activation patterns (β maps) across all successful stop (SS) trials from one run. We then computed pattern similarity between this template and each Go-trial β map from the other run, yielding a Safe-state similarity time series. The procedure was repeated with runs swapped. Searchlight maps indexing the correlation between trial-wise Safe-state similarity and Go-trial RT were Fisher z–transformed and carried forward to group-level inference using permutation testing in FSL PALM.

Speed-state–like representation (speed-biased): Analogously, we defined a Speed-state template to capture speed-biased execution states. In the within-subject approach (Supplementary Materials), the template was constructed by averaging β maps from the fastest 10% of Go trials within one run, chosen to yield a trial count comparable to successful stop trials (given ∼50% stop success). Pattern similarity between this template and Go-trial β maps from the other run was computed to produce a trial-wise similarity time series; run direction was swapped and averaged as above. Trial-wise Speed-state similarity was then related to RT using the same searchlight, Fisher z transformation, and PALM-based permutation framework.

Mitigating temporal autocorrelation and ensuring independence (primary analysis): The IMAGEN dataset included only a single SST run, precluding cross-run template construction. Within-run temporal autocorrelation can inflate pattern similarity because trials that occur close in time often exhibit correlated activation patterns unrelated to the targeted control state. This concern is particularly salient for Speed-state templates, as the fastest Go trials may cluster within brief epochs rather than being evenly distributed across the run. Such clustering can bias similarity estimates, carry forward into subsequent between-region representational coherence analyses, and confound comparisons with Safe-state effects, especially if the magnitude or structure of autocorrelation differs across waves/datasets, complicating cross-cohort interpretation.

To mitigate this, our primary analyses used leave-one-subject-out (LOSO) group-average templates for both Speed-state and Safe-state representations. Specifically: For each participant, we computed participant-level Speed-state and Safe-state summary patterns by averaging β maps across the fastest 10% of Go trials (Speed-state) and across successful stop trials (Safe-state). For a given target participant, we formed LOSO templates by averaging the corresponding participant-level patterns from all other participants (excluding the target participant) ^48,97^. We then quantified trial-wise similarity between the target participant’s Go-trial β maps and these LOSO templates, producing Speed-state and Safe-state similarity time series that are independent of the target participant’s own data. These trial-wise similarity values were related to RT using the same searchlight and group-level permutation testing pipeline. This LOSO procedure was applied consistently in both ABCD and IMAGEN. Parallel analyses using the within-subject, cross-run templates produce convergent results and are presented in the Supplementary Materials (Supplementary **Fig. S7-9**).

#### Representational coherence analysis

To characterize how the speed–caution balance is coordinated across distributed control systems, we quantified the trial-wise expression of Speed-state and Safe-state representations within each ROI and then measured inter-regional coherence of these state expressions. ROI-level Speed-state and Safe-state time series. ROIs were defined using the Brainnetome atlas ^98^. For each participant and ROI, we computed two trial-wise representational similarity time series on Go trials: a Speed-state series indexing how strongly the ROI’s voxel wise activity pattern on each Go trial matched the Speed-state LOSO template, and a Safe-state series indexing similarity to the Safe-state LOSO template (templates constructed as described above). This yielded, for every ROI, two “runner” time series reflecting moment-to-moment fluctuations in speed-biased versus caution-biased state expression.

Constructing representational coherence matrices (**Fig. 1h**). For each participant, we built two ROI×ROI coherence matrices by correlating the runner time series between all ROI pairs across Go trials: (i) Speed-state coherence (Speed-state time series × Speed-state time series), and (ii) Safe-state coherence (Safe-state time series × Safe-state time series). Correlation coefficients were Fisher z–transformed prior to group-level analyses. In this framework, stronger within-state coherence indicates tighter synchronization of state expression across regions, coordination among regions jointly supporting an execution-biased (Speed-state) versus caution-biased (Safe-state) control mode. Direct contrasts between the Speed-state and Safe-state coherence matrices therefore provide a network-level readout of how the brain reconfigures inter-regional coordination to implement and stabilize the speed–caution balance.

#### Elastic Net Regression

To test whether distributed neural features predict individual differences in core behavioral phenotypes, we used Elastic Net regression, a regularized multivariate model well suited for high-dimensional neuroimaging data with correlated predictors. Elastic Net combines L1 (lasso) and L2 (ridge) penalties, enabling coefficient shrinkage with sparse feature selection while allowing correlated feature groups to be retained ^99^. Models were fit separately for each behavioral outcome, including stop-signal reaction time (SSRT) and Go RT variability (IIRV).

In parallel models, predictors comprised ROI-level coupling metrics derived from the speed–caution (Speed-state/Safe-state) analyses. Within each training set, all neural features were z-scored, and the same scaling parameters were applied to the held-out test set to prevent information leakage. Behavioral outcomes were standardized within the training data using the same procedure.

Elastic Net models were trained using 10-fold cross-validation. In each fold, 90% of participants were used for training and 10% were held out for testing. Hyperparameters were tuned only within the training data using nested cross-validation: the overall regularization strength (α) and the mixing parameter (l1_ratio). Optimal hyperparameters were selected by minimizing mean squared error in the inner loop. The model fit with the selected parameters was then applied to the held-out test set. Predicted scores from all folds were concatenated to yield one out-of-sample prediction per participant. Model performance was quantified as (i) the Pearson correlation between observed and predicted scores and (ii) partial correlations controlling for age, sex, head motion, and scanner site (see results in **Table S4**).

To assess temporal generalizability, models trained at one developmental wave were transferred to predict behavior at other waves and independent dataset. Feature scaling parameters and regression coefficients learned in the training wave were held fixed and applied to the target wave, providing a stringent test of stability across development.

For interpretability, we summarized the anatomical/system-level distribution of non-zero coefficients (e.g., by functional systems). These summaries were descriptive and were not used to guide model fitting or statistical inference.

#### Connectome-Based Predictive Modeling

To identify functional brain networks predictive of individual differences in inhibitory control and response variability, we applied Connectome-Based Predictive Modeling (CPM, **Fig. 1h**), a validated data-driven framework for linking whole-brain functional connectivity to behavior ^100-102^. CPM analyses were conducted separately for each behavioral measure, including stop-signal reaction time (SSRT) and Go reaction time variability (IIRV). All analyses were performed within a cross-validated framework, with strict separation of training and testing data to prevent circularity.

Feature Selection and Model Construction: We implemented a 10-fold cross-validation procedure. In each fold, participants were randomly divided into a training set (90%) and an independent testing set (10%). Feature selection was performed exclusively within the training data. For each fold, we computed the partial correlation between each edge in the whole-brain functional connectivity matrix and the behavioral measure of interest, controlling for nuisance covariates where appropriate.

Edges showing a significant association with behavior (P < 0.01, uncorrected) were retained as predictive features, consistent with established CPM implementations. Selected edges were separated into positive networks (edges positively associated with behavior) and negative networks (edges negatively associated with behavior). For each participant, network strength was calculated by summing the connectivity values of all edges within each network.

To avoid data leakage, network strengths and behavioral scores were Z-scored using parameters derived from the training set only, and the same normalization parameters were then applied to the test set. Predictive models were trained using robust linear regression to reduce sensitivity to outliers. A combined model including both positive and negative network strengths as independent predictors was fitted within each training fold.

#### Prediction and Model Evaluation

Model parameters estimated from the training data were applied to the held-out test data to generate predicted behavioral scores. This procedure was repeated across all folds, yielding a predicted score for each participant. Predictive performance was quantified using (i) the Pearson correlation between observed and predicted scores and (ii) the partial correlation between observed and predicted scores controlling for age, sex, head motion, and scanner site. The partial correlation metric assesses the variance in behavior uniquely explained by functional connectivity beyond potential confounds (see results in **Table S5**).

#### Longitudinal Generalization

To evaluate temporal stability and generalizability, we conducted cross-wave prediction analyses. Models trained at one wave (e.g., baseline) were fixed and applied directly to predict behavior at another point (e.g., follow-up), using the same feature definitions and regression coefficients. This procedure provided a stringent test of external validity across development.

#### Consensus Networks

To characterize the spatial organization of predictive networks, we defined consensus networks as edges selected in all cross-validation folds. These consensus networks were used for visualization and anatomical interpretation but were not used for model fitting or evaluation.

#### Canonical Correlation Analysis

Canonical correlation analysis (CCA) was used to examine multivariate associations between distributed brain-behavior coupling strength or interconnected representational coherence and dimensional measures of mental health. Brain variables were derived from network-level representational coherence matrices computed separately for Speed-state and Safe-state representations, or the ROI-level brain-behavior coupling strength computed by correlating Speed-state/Safe-state and trial-wise RT. Clinical variables indexed broad psychopathology and adaptive functioning, as assessed using standardized instruments (e.g., the Child Behavior Checklist (CBCL) in ABCD; cohort-specific measures in IMAGEN).

To mitigate the high dimensionality of edge-level connectivity and reduce the risk of overfitting, representational coherence values were first aggregated at the network–network level.

Specifically, coherence values for all ROI–ROI edges belonging to each pair of large-scale functional networks were averaged, yielding a compact and interpretable brain feature set for each participant. This procedure was performed separately for Speed-state and Safe-state representations and for each trial type. All brain features were standardized prior to analysis. Clinical measures were organized into dimensional symptom domains following established scoring conventions for each instrument. Clinical variables were likewise standardized to place brain and behavioral variables on comparable scales.

CCA was conducted to identify pairs of latent canonical variates that maximize the correlation between linear combinations of brain connectivity features and clinical symptom dimensions. Separate CCA models were fit for Speed-state and Safe-state representations to dissociate their contributions to psychopathology. All analyses were performed independently at each developmental time point. Importantly, we examined if the longitudinal changes in brain measurements can predict developmental effects in clinical symptoms changes.

To obtain robust statistical inference and control for multiple testing, we employed permutation-based CCA (PermCCA) ^103^. Subject labels in the clinical data were randomly permuted 10,000 times while preserving the correlation structure within the brain feature matrix. For each permutation, the maximal canonical correlation was recomputed, generating a null distribution against which the observed canonical correlation was evaluated. Statistical significance was assessed using family-wise error–corrected p-values based on this permutation distribution.

To interpret the multivariate associations, canonical loadings were examined for both brain and clinical variables. Brain loadings were used to identify the neural features contributing most strongly to each canonical variate, and clinical loadings were used to summarize the symptom dimensions most strongly aligned with the corresponding brain pattern. These analyses were conducted for descriptive interpretation only and were not used to guide model fitting or statistical inference.

## Data availability

The ABCD data are publicly available via the NIMH Data Archive (NDA). The ABCD data repository grows and changes over time, researchers with access to ABCD data will be able to download the data: (doi: 10.15154/1503209). IMAGEN data can be accessed by email at https://imagen-project.org/.

## Code availability

Functional MRI data preprocessing and statistical analyses were performed on the SPM12 and FSL6, Nilearn (version 0.10.1), and Matlab 2020. Code to analyze the data will be made available at https://osf.io/7shk5 upon publication.

## Acknowledgement

This work received support from the following sources: National Institutes of Health MH124816 (W.C.), MH121069 (V.M.), HD094623 (V.M.), and MH137325 (V.M.), Stanford Maternal and Child Health Research Institute Grant (W.C.), Stanford University Department of Psychiatry Innovator Grant (W.C.), and Arizona Alzheimer’s Consortium & State of Arizona DHS Pilot Grant (L.Z.).

The ABCD Study is supported by the National Institutes of Health and additional federal partners under award numbers U01DA041022, U01DA041028, U01DA041048, U01DA041089, U01DA041106, U01DA041117, U01DA041120, U01DA041134, U01DA041148, U01DA041156, U01DA041174, U24DA041123, U24DA041147, U01DA041093, and U01DA041025. A full list of supporters is available at https://abcdstudy.org/federal-partners.html. A listing of participating sites and a complete listing of the study investigators can be found at https://abcdstudy.org/scientists/workgroups/. ABCD consortium investigators designed and implemented the study and/or provided data but did not necessarily participate in analysis or writing of this report. This manuscript reflects the views of the authors and may not reflect the opinions or views of the NIH or ABCD consortium investigators.

Additional support was provided by the European Union-funded FP6 Integrated Project IMAGEN (Reinforcement-related behaviour in normal brain function and psychopathology) (LSHM-CT-2007-037286), the Horizon 2020 funded ERC Advanced Grant ‘STRATIFY’ (Brain network based stratification of reinforcement-related disorders) (695313), Horizon Europe ‘environMENTAL’, grant no: 101057429, UK Research and Innovation (UKRI) Horizon Europe funding guarantee (10041392 and 10038599), Human Brain Project (HBP SGA 2, 785907, and HBP SGA 3, 945539), the Chinese government via the Ministry of Science and Technology (MOST). The German Center for Mental Health (DZPG), the Bundesministerium für Bildung und Forschung (BMBF grants 01GS08152; 01EV0711; Forschungsnetz AERIAL 01EE1406A, 01EE1406B; Forschungsnetz IMAC-Mind 01GL1745B), the Deutsche Forschungsgemeinschaft (DFG project numbers 458317126 [COPE], 186318919 [FOR 1617], 178833530 [SFB 940], 386691645 [NE 1383/14-1], 402170461 [TRR 265], 454245598 [IRTG 2773]), the Medical Research Foundation and Medical Research Council (grants MR/R00465X/1 and MR/S020306/1), the National Institutes of Health (NIH) funded ENIGMA-grants 5U54EB020403-05, 1R56AG058854-01 and U54 EB020403 as well as NIH R01DA049238, the National Institutes of Health, Science Foundation Ireland (16/ERCD/3797). NSFC grant 82150710554. Further support was provided by grants from: the Eranet Neuron (Grant ANR-18-NEUR00002-01– ADORe); Agence Nationale de la Recherche (Grant ANR-12-SAMA-0004-GeBra); Assistance-Publique Hôpitaux-de-Paris and INSERM (interface grant); Paris Descartes University (Grant collaborative-project-2010); Paris Sud University (Grant IDEX-2012); Fondation de l’Avenir (Grant AP-RM-17-013); Fondation de France (Grant 00081242); Fédération pour la Recherche sur le Cerveau, and Fondation pour la Recherche Médicale (Grants DPA20140629802 and ADOLIMIS DPP20151033945); the Ile-de-France Region (Action 16700103-grant to QIM– VEAVE, n°23002745–23002747).

## Author contributions

Study design: Z.G., W.C., V.M.; Data Analysis: Z.G., W.C., L.Z.; Manuscript Drafting: Z.G., W.C.; Manuscript Editing: Z.G., H.G., V.M., W.C.; T.B., G.J.B., A.L.W.B., R.B., S.D., P.G., A.G., A.H., F.N., D.P.O., T.P., L.P., M.N.S., S.H., N.H., N.V., H.W., R.W., P.W., and G.S. are principal investigators of IMAGEN.

## References

1 Munakata, Y. et al. A unified framework for inhibitory control. Trends in cognitive sciences 15, 453–459 (2011).

2 Chambers, C. D., Garavan, H. & Bellgrove, M. A. Insights into the neural basis of response inhibition from cognitive and clinical neuroscience. Neuroscience & biobehavioral reviews 33, 631–646 (2009).

3 Wessel, J. R. & Anderson, M. C. Neural mechanisms of domain-general inhibitory control. Trends in Cognitive Sciences 28, 124–143 (2024).

4 Lipszyc, J. & Schachar, R. Inhibitory control and psychopathology: a meta-analysis of studies using the stop signal task. Journal of the International Neuropsychological Society 16, 1064–1076 (2010).

5 Yan, H. et al. Charting the neural circuits disruption in inhibitory control and its subcomponents across psychiatric disorders: A neuroimaging meta-analysis. Progress in Neuro-Psychopharmacology and Biological Psychiatry 119, 110618 (2022).

6 Mirabella, G. Inhibitory control and impulsive responses in neurodevelopmental disorders. Developmental Medicine & Child Neurology 63, 520–526 (2021).

7 Constantinidis, C. & Luna, B. Neural substrates of inhibitory control maturation in adolescence. Trends in neurosciences 42, 604–616 (2019).

8 Williams, B. R., Ponesse, J. S., Schachar, R. J., Logan, G. D. & Tannock, R. Development of inhibitory control across the life span. Developmental psychology 35, 205 (1999).

9 Bedard, A.-C. et al. The development of selective inhibitory control across the life span. Developmental neuropsychology 21, 93–111 (2002).

10 Ordaz, S. J., Foran, W., Velanova, K. & Luna, B. Longitudinal growth curves of brain function underlying inhibitory control through adolescence. Journal of Neuroscience 33, 18109–18124 (2013).

11 Vara, A. S., Pang, E. W., Vidal, J., Anagnostou, E. & Taylor, M. J. Neural mechanisms of inhibitory control continue to mature in adolescence. Developmental Cognitive Neuroscience 10, 129–139 (2014).

12 Rubia, K., Smith, A. B., Taylor, E. & Brammer, M. Linear age-correlated functional development of right inferior fronto-striato-cerebellar networks during response inhibition and anterior cingulate during error-related processes. Human brain mapping 28, 1163–1177 (2007).

13 Rubia, K. et al. Progressive increase of frontostriatal brain activation from childhood to adulthood during event-related tasks of cognitive control. Human brain mapping 27, 973–993 (2006).

14 Vink, M. et al. Frontostriatal activity and connectivity increase during proactive inhibition across adolescence and early adulthood. Human brain mapping 35, 4415–4427 (2014).

15 Luna, B., Marek, S., Larsen, B., Tervo-Clemmens, B. & Chahal, R. An integrative model of the maturation of cognitive control. Annual review of neuroscience 38, 151–170 (2015).

16 Andersen, S. L. Trajectories of brain development: point of vulnerability or window of opportunity? Neuroscience & Biobehavioral Reviews 27, 3–18 (2003).

17 Fuhrmann, D., Knoll, L. J. & Blakemore, S.-J. Adolescence as a sensitive period of brain development. Trends in cognitive sciences 19, 558–566 (2015).

18 Casey, B. J., Getz, S. & Galvan, A. The adolescent brain. Developmental review 28, 62–77 (2008).

19 Dahl, R. E. Adolescent brain development: a period of vulnerabilities and opportunities. Keynote address. Annals of the new York Academy of Sciences 1021, 1–22 (2004).

20 Shaw, P. et al. Attention-deficit/hyperactivity disorder is characterized by a delay in cortical maturation. Proceedings of the national academy of sciences 104, 19649–19654 (2007).

21 Castellanos, F. X. et al. Developmental trajectories of brain volume abnormalities in children and adolescents with attention-deficit/hyperactivity disorder. Jama 288, 1740–1748 (2002).

22 Friedman, L. A. & Rapoport, J. L. Brain development in ADHD. Current opinion in neurobiology 30, 106–111 (2015).

23 Braver, T. S. The variable nature of cognitive control: a dual mechanisms framework. Trends in cognitive sciences 16, 106–113 (2012).

24 Aron, A. R. From reactive to proactive and selective control: developing a richer model for stopping inappropriate responses. Biological psychiatry 69, e55–e68 (2011).

25 Cai, W. et al. Both reactive and proactive control are deficient in children with ADHD and predictive of clinical symptoms. Translational psychiatry 13, 179 (2023).

26 Cai, W. & Mizuno, Y. Dynamic modeling in neurocognitive frameworks of childhood ADHD: a review of inhibitory control and reward systems. Translational Psychiatry (2026).

27 Van De Laar, M. C., van den Wildenberg, W. P., van Boxtel, G. J. & van der Molen, M. W. Lifespan changes in global and selective stopping and performance adjustments. Frontiers in psychology 2, 357 (2011).

28 Luna, B., Garver, K. E., Urban, T. A., Lazar, N. A. & Sweeney, J. A. Maturation of cognitive processes from late childhood to adulthood. Child development 75, 1357–1372 (2004).

29 Logan, G. D. & Cowan, W. B. On the ability to inhibit thought and action: A theory of an act of control. Psychological review 91, 295 (1984).

30 Van Gerven, P. W., Hurks, P. P., Bovend’Eerdt, T. J. & Adam, J. J. Switch hands! Mapping proactive and reactive cognitive control across the life span. Developmental psychology 52, 960 (2016).

31 Gonthier, C., Zira, M., Colé, P. & Blaye, A. Evidencing the developmental shift from reactive to proactive control in early childhood and its relationship to working memory. Journal of experimental child psychology 177, 1–16 (2019).

32 Munakata, Y., Snyder, H. R. & Chatham, C. H. Developing cognitive control: Three key transitions. Current directions in psychological science 21, 71–77 (2012).

33 Chevalier, N., Martis, S. B., Curran, T. & Munakata, Y. Metacognitive processes in executive control development: The case of reactive and proactive control. Journal of cognitive neuroscience 27, 1125–1136 (2015).

34 Levy, B. J. & Wagner, A. D. Cognitive control and right ventrolateral prefrontal cortex: reflexive reorienting, motor inhibition, and action updating. Annals of the New York academy of sciences 1224, 40–62 (2011).

35 Swick, D., Ashley, V. & Turken, U. Are the neural correlates of stopping and not going identical? Quantitative meta-analysis of two response inhibition tasks. Neuroimage 56, 1655–1665 (2011).

36 Cai, W., Chen, T., Ide, J. S., Li, C.-S. R. & Menon, V. Dissociable fronto-operculum-insula control signals for anticipation and detection of inhibitory sensory cue. Cerebral cortex 27, 4073–4082 (2017).

37 Rushworth, M., Hadland, K., Paus, T. & Sipila, P. Role of the human medial frontal cortex in task switching: a combined fMRI and TMS study. Journal of neurophysiology 87, 2577–2592 (2002).

38 Isoda, M. & Hikosaka, O. Switching from automatic to controlled action by monkey medial frontal cortex. Nature Neuroscience 10, 240–248 (2007). 10.1038/nn1830

39 Sridharan, D., Levitin, D. J. & Menon, V. A critical role for the right fronto-insular cortex in switching between central-executive and default-mode networks. Proc Natl Acad Sci U S A 105, 12569–12574 (2008). 10.1073/pnas.0800005105

40 Ham, T., Leff, A., de Boissezon, X., Joffe, A. & Sharp, D. J. Cognitive control and the salience network: an investigation of error processing and effective connectivity. J Neurosci 33, 7091–7098 (2013). 10.1523/jneurosci.4692-12.2013

41 Achterberg, M., Peper, J. S., van Duijvenvoorde, A. C., Mandl, R. C. & Crone, E. A. Frontostriatal white matter integrity predicts development of delay of gratification: a longitudinal study. Journal of Neuroscience 36, 1954–1961 (2016).

42 Singh, M. et al. Longitudinal developmental trajectories of inhibition and white-matter maturation of the fronto-basal-ganglia circuits. Developmental Cognitive Neuroscience 58, 101171 (2022).

43 Vijayakumar, N. et al. Prefrontal structural correlates of cognitive control during adolescent development: a 4-year longitudinal study. Journal of Cognitive neuroscience 26, 1118–1130 (2014).

44 Borst, G. et al. Folding of the anterior cingulate cortex partially explains inhibitory control during childhood: a longitudinal study. Developmental cognitive neuroscience 9, 126–135 (2014).

45 Dinstein, I., Heeger, D. J. & Behrmann, M. Neural variability: friend or foe? Trends in cognitive sciences 19, 322–328 (2015).

46 Coutanche, M. N. Distinguishing multi-voxel patterns and mean activation: why, how, and what does it tell us? *Cognitive, Affective*, & Behavioral Neuroscience 13, 667–673 (2013).

47 Tsikonofilos, K., Kumar, A., Ampatzis, K., Garrett, D. D. & Månsson, K. N. The promise of investigating neural variability in psychiatric disorders. Biological Psychiatry (2025).

48 Gao, Z. et al. Reduced temporal and spatial stability of neural activity patterns predict cognitive control deficits in children with ADHD. Nature Communications 16, 2346 (2025).

49 Garrett, D. D. et al. Moment-to-moment brain signal variability: a next frontier in human brain mapping? Neuroscience & Biobehavioral Reviews 37, 610–624 (2013).

50 Armbruster-Genç, D. J., Ueltzhöffer, K. & Fiebach, C. J. Brain signal variability differentially affects cognitive flexibility and cognitive stability. Journal of Neuroscience 36, 3978–3987 (2016).

51 Cai, W. et al. Latent brain state dynamics distinguish behavioral variability, impaired decision-making, and inattention. Molecular psychiatry 26, 4944–4957 (2021).

52 Cai, W., Chen, T., Szegletes, L., Supekar, K. & Menon, V. Aberrant time-varying cross-network interactions in children with attention-deficit/hyperactivity disorder and the relation to attention deficits. Biological Psychiatry: Cognitive Neuroscience and Neuroimaging 3, 263–273 (2018).

53 Nomi, J. S., Bolt, T. S., Ezie, C. C., Uddin, L. Q. & Heller, A. S. Moment-to-moment BOLD signal variability reflects regional changes in neural flexibility across the lifespan. Journal of Neuroscience 37, 5539–5548 (2017).

54 Grady, C. L. & Garrett, D. D. Understanding variability in the BOLD signal and why it matters for aging. Brain imaging and behavior 8, 274–283 (2014).

55 Menon, V. 20 years of the default mode network: A review and synthesis. Neuron 111, 2469–2487 (2023).

56 Menon, V. Large-scale brain networks and psychopathology: a unifying triple network model. Trends in cognitive sciences 15, 483–506 (2011).

57 Casey, B. J. et al. The adolescent brain cognitive development (ABCD) study: imaging acquisition across 21 sites. Developmental cognitive neuroscience 32, 43–54 (2018).

58 Volkow, N. D. et al. The conception of the ABCD study: From substance use to a broad NIH collaboration. Developmental cognitive neuroscience 32, 4–7 (2018).

59 Mascarell Maričić, L., et al. The IMAGEN study: a decade of imaging genetics in adolescents. Molecular psychiatry 25, 2648–2671 (2020).

60 Schumann, G. et al. The IMAGEN study: reinforcement-related behaviour in normal brain function and psychopathology. Molecular psychiatry 15, 1128–1139 (2010).

61 Tamm, L. et al. Reaction time variability in ADHD: a review. Neurotherapeutics 9, 500–508 (2012).

62 Hervey, A. S. et al. Reaction time distribution analysis of neuropsychological performance in an ADHD sample. Child Neuropsychology 12, 125–140 (2006).

63 Klein, C., Wendling, K., Huettner, P., Ruder, H. & Peper, M. Intra-subject variability in attention-deficit hyperactivity disorder. Biological psychiatry 60, 1088–1097 (2006).

64 Willcutt, E. G., Doyle, A. E., Nigg, J. T., Faraone, S. V. & Pennington, B. F. Validity of the executive function theory of attention-deficit/hyperactivity disorder: a meta-analytic review. Biological psychiatry 57, 1336–1346 (2005).

65 Kriegeskorte, N., Mur, M. & Bandettini, P. A. Representational similarity analysis-connecting the branches of systems neuroscience. Frontiers in systems neuroscience 2, 249 (2008).

66 Mumford, J. A., Turner, B. O., Ashby, F. G. & Poldrack, R. A. Deconvolving BOLD activation in event-related designs for multivoxel pattern classification analyses. Neuroimage 59, 2636–2643 (2012).

67 Band, G. P., Van Der Molen, M. W. & Logan, G. D. Horse-race model simulations of the stop-signal procedure. Acta psychologica 112, 105–142 (2003).

68 Verbruggen, F. & Logan, G. D. Models of response inhibition in the stop-signal and stop-change paradigms. Neuroscience & Biobehavioral Reviews 33, 647–661 (2009).

69 Tillman, C. M., Thorell, L. B., Brocki, K. C. & Bohlin, G. Motor response inhibition and execution in the stop-signal task: development and relation to ADHD behaviors. Child Neuropsychology 14, 42–59 (2007).

70 Albert, J. et al. The development of selective stopping: Qualitative and quantitative changes from childhood to early adulthood. Developmental Science 25, e13210 (2022).

71 Schachar, R. & Logan, G. D. Impulsivity and inhibitory control in normal development and childhood psychopathology. Developmental psychology 26, 710 (1990).

72 Fosco, W. D., Hawk Jr, L. W., Colder, C. R., Meisel, S. N. & Lengua, L. J. The development of inhibitory control in adolescence and prospective relations with delinquency. Journal of Adolescence 76, 37–47 (2019).

73 Wang, H. et al. Functional connectivity predicts individual development of inhibitory control during adolescence. Cerebral Cortex 31, 2686–2700 (2021).

74 Corrigan, N. M. et al. Myelin development in cerebral gray and white matter during adolescence and late childhood. Neuroimage 227, 117678 (2021).

75 Vanes, L. D. et al. White matter tract myelin maturation and its association with general psychopathology in adolescence and early adulthood. Human brain mapping 41, 827–839 (2020).

76 Chikazoe, J. et al. Preparation to inhibit a response complements response inhibition during performance of a stop-signal task. Journal of Neuroscience 29, 15870–15877 (2009).

77 Swann, N. C. et al. Roles for the pre-supplementary motor area and the right inferior frontal gyrus in stopping action: electrophysiological responses and functional and structural connectivity. Neuroimage 59, 2860–2870 (2012).

78 Jahfari, S. et al. How preparation changes the need for top–down control of the basal ganglia when inhibiting premature actions. Journal of Neuroscience 32, 10870–10878 (2012).

79 Swann, N. C., Tandon, N., Pieters, T. A. & Aron, A. R. Intracranial electroencephalography reveals different temporal profiles for dorsal-and ventro-lateral prefrontal cortex in preparing to stop action. Cerebral cortex 23, 2479–2488 (2013).

80 Shrout, P. E. & Rodgers, J. L. Psychology, science, and knowledge construction: Broadening perspectives from the replication crisis. Annual review of psychology 69, 487–510 (2018).

81 Nosek, B. A. et al. Replicability, robustness, and reproducibility in psychological science. Annual review of psychology 73, 719–748 (2022).

82 Collaboration, O. S. Estimating the reproducibility of psychological science. Science 349, aac4716 (2015).

83 Verbruggen, F. et al. A consensus guide to capturing the ability to inhibit actions and impulsive behaviors in the stop-signal task. elife 8, e46323 (2019).

84 Achenbach, T. M. Manual for ASEBA school-age forms & profiles. *University of Vermont, Research Center for Children*, Youth & Families (2001).

85 Lynam, D. Development of a short form of the UPPS-P Impulsive Behavior Scale. Unpublished technical report (2013).

86 Lynam, D., Smith, G., Cyders, M., Fischer, S. & Whiteside, S. The UPPS-P: A multidimensional measure of risk for impulsive behavior. Unpublished technical report (2007).

87 Goodman, R., Ford, T., Richards, H., Gatward, R. & Meltzer, H. The development and well-being assessment: Description and initial validation of an integrated assessment of child and adolescent psychopathology. Journal of child psychology and psychiatry 41, 645–655 (2000).

88 Goodman, R. The Strengths and Difficulties Questionnaire: a research note. Journal of child psychology and psychiatry 38, 581–586 (1997).

89 Zeithamova, D., de Araujo Sanchez, M.-A. & Adke, A. Trial timing and pattern-information analyses of fMRI data. Neuroimage 153, 221–231 (2017).

90 Kriegeskorte, N., Goebel, R. & Bandettini, P. Information-based functional brain mapping. Proceedings of the National Academy of Sciences 103, 3863–3868 (2006).

91 Zheng, L. et al. Reduced fidelity of neural representation underlies episodic memory decline in normal aging. Cerebral Cortex 28, 2283–2296 (2018).

92 Viganò, S. & Piazza, M. Distance and direction codes underlie navigation of a novel semantic space in the human brain. Journal of Neuroscience 40, 2727–2736 (2020).

93 Gao, Z. et al. Context free and context-dependent conceptual representation in the brain. Cerebral Cortex 33, 152–166 (2023).

94 Gao, Z. et al. Distinct and common neural coding of semantic and non-semantic control demands. NeuroImage 236, 118230 (2021).

95 Winkler, A. M., Ridgway, G. R., Webster, M. A., Smith, S. M. & Nichols, T. E. Permutation inference for the general linear model. Neuroimage 92, 381–397 (2014).

96 Winkler, A. M., Ridgway, G. R., Douaud, G., Nichols, T. E. & Smith, S. M. Faster permutation inference in brain imaging. Neuroimage 141, 502–516 (2016).

97 Chen, J. et al. Shared memories reveal shared structure in neural activity across individuals. Nature neuroscience 20, 115–125 (2017).

98 Fan, L. et al. The human brainnetome atlas: a new brain atlas based on connectional architecture. Cerebral cortex 26, 3508–3526 (2016).

99 Friedman, J. H., Hastie, T. & Tibshirani, R. Regularization paths for generalized linear models via coordinate descent. Journal of statistical software 33, 1–22 (2010).

100 Shen, X. et al. Using connectome-based predictive modeling to predict individual behavior from brain connectivity. nature protocols 12, 506–518 (2017).

101 Rosenberg, M. D. et al. A neuromarker of sustained attention from whole-brain functional connectivity. Nature neuroscience 19, 165–171 (2016).

102 Yoo, K. et al. Connectome-based predictive modeling of attention: Comparing different functional connectivity features and prediction methods across datasets. Neuroimage 167, 11–22 (2018).

103 Winkler, A. M., Renaud, O., Smith, S. M. & Nichols, T. E. Permutation inference for canonical correlation analysis. Neuroimage 220, 117065 (2020).

