## Supplementary Materials for "Protracted Maturation of Proactive and Reactive Systems Predicts Cognitive Stability and Psychopathology : A longitudinal multi-cohort study"

**Supplementary Methods**

**Cohort demographics and samples**

**Cohort 1: Late Childhood to Early Adolescence (ABCD 4.0)**

The ABCD study is a large-scale longitudinal neuroimaging project that enrolled over 11,800 children at ages 9-10 (Casey et al., 2018; Volkow et al., 2018). We utilized data from Release 4.0, which comprises clinical, behavioral, and neuroimaging assessments collected at baseline (ages 9–10; hereafter 'Age 10') and the 2-year follow-up (hereafter 'Age 12'). Although partial data from the 4-year follow-up were available, we restricted our analyses to these initial two timepoints to maximize longitudinal sample retention and specifically target the developmental transition from late childhood to early adolescence. Following rigorous quality control screening for clinical, behavioral and neuroimaging quality (see *Method* for criteria), a final sample of 1,928 participants (967 females) was included in the analyses.

**Cohort 2: Mid-Adolescence to Young Adulthood (IMAGEN)**

As Release 4.0 of the ABCD study does not contain the full range of adolescent ages, we used the IMAGEN study to characterize developmental trajectories from middle adolescence to young adulthood. The IMAGEN study is also a large-scale longitudinal project designed to assess neurobiological and environmental risk factors for adolescent mental health (Mascarell Maričić et al., 2020; Schumann et al., 2010). We utilized data from three distinct waves: ages 14, ages 18, and age 22 (hereafter, Age 14, Age 18, and Age 22). This dataset was selected to complement the ABCD cohort, extending our investigation from adolescence into early adulthood. Given the eight-year span of the IMAGEN study, restricting the analysis to a 'complete-case' cohort (participants present at all three time points) would result in significant attrition and reduced statistical power. Therefore, to maximize the effective sample size and minimize selection bias, we utilized all available high-quality data at each individual wave, regardless of participation in other waves. This unbalanced longitudinal structure was explicitly handled using Linear Mixed Effects (LME) models, which provide robust and unbiased estimates of developmental trajectories even in the presence of missing temporal data. Following rigorous quality control screening for clinical, behavioral, and neuroimaging data (see *Methods* for criteria), a total of 1247 participants at Age 14 (657 females), 1151 participants at Age 18 (611 females), and 973 participants at Age 22 (512 females) were included in the analyses.

To ensure that our results were not confounded by attrition bias or missing data patterns, we conducted a sensitivity analysis using the 'complete-case' subsample (see **Table S2**. i.e., only participants with valid data across all three time points, 464 participants were included in the analyses, 258 females) to replicate our primary findings.

**Longitudinal comparison of SSRT**

We examined the distribution of stop accuracy at each wave for each cohort to assure that the comparisons were not biased by the difference of stopping performance between waves.

In the IMAGEN cohort, the preliminary analysis revealed that the distribution of stop accuracy at Age 14 deviated from the optimal 50% center observed at Age 18 and Age 22, introducing significant bias into SSRT estimation (**Table S1**, also Supplementary Methods). To ensure unbiased comparison in SSRT across study waves, we restricted this behavioral analysis to participants whose stop accuracies fall between 45% and 55%. This inclusion criteria yield samples of 605 participants at Age 14 (331 females), 1169 participants at Age 18 (614 females), and 997 participants at Age 22 (525 females).

**Neuroimaging data**

Imaging protocols for the ABCD and IMAGEN stop-signal task (SST) datasets followed previously published, multi-site acquisition schemes. In the ABCD dataset, participants completed two runs of the SST during fMRI scanning using a multiband gradient-echo EPI sequence (multiband factor = 6) on 3 T scanners (TR = 800 ms, TE = 30 ms, voxel size = 2.4 × 2.4 × 2.4 mm). High-resolution T1-weighted structural images (MPRAGE; 1 mm isotropic) were also acquired. In the IMAGEN dataset, participants completed one run of the SST during fMRI scanning using single-shot gradient-echo EPI on 3 T scanners (TR = 2200 ms, TE = 30 ms, voxel size = 3.4 × 3.4 × 3.4 mm). High-resolution T1-weighted MPRAGE images (1 mm isotropic) were collected at all sites. Both datasets were acquired across multiple MRI sites, with minor scanner-specific variations detailed in their respective protocols. To control potential scanner effects, MRI site was included as a nuisance covariate in all statistical analyses.

**Preprocessing**

For the ABCD dataset, we used the minimally processed data from release 4.0 (Collection #2573), which included distortion and motion correction. Additional preprocessing was performed using Nilearn and FSL FLIRT to achieve spatial normalization and minimal smoothing. Specifically: (1) initial volumes were removed (Siemens: 8; Philips: 8; GE DV25: 5; GE DV26 and other GE versions: 16); (2) a mean functional image was computed across time using mean_img from Nilearn; (3) this mean image was registered to an echo-planar imaging (EPI) template in MNI152 space using FSL FLIRT, and the corresponding affine transformation was estimated; (4) the full fMRI time series was spatially normalized to MNI152 space using this affine transformation; and (5) the normalized images were minimally smoothed with a Gaussian kernel of 2 mm full width at half maximum (FWHM) using Nilearn’s smooth_img function.

For the IMAGEN dataset, preprocessing followed a similar pipeline to ensure methodological consistency and comparability. Functional images underwent motion correction, followed by slice-timing correction (performed only for IMAGEN due to its longer TR of 2.2 s). The mean functional image was then registered to the MNI152 EPI template using FSL FLIRT, and the resulting affine transformation was applied to the full time series. Finally, spatial smoothing was performed with a Gaussian kernel of 2 mm FWHM.

**Single-trial estimation**

The GLMs were performed separately to estimate the activation pattern for each trial using a Least Square–Separate (LS-S) approach (Mumford et al., 2012; Zeithamova et al., 2017), in which the trial of interest was modelled as one regressor, with all other trials modelled as separate regressors. Specifically, each single-trial GLM included five regressors: (1) the single trial of interest from one condition, for instance one Successful Stop trial (SS); (2) all other SS trials; (3) all the Go trials; (4) all the US trials; (5) all other trials not being included in the previous four regressors. Six head motion parameters were included as regressors of no interest. Each event was modelled by defining the time of stimulus onset as a stick function and convolving this stick function with a canonical hemodynamic response function. This approach was applied to both the ABCD and IMAGEN datasets. To account for differences in temporal sampling, temporal autocorrelations were modeled using the FAST method for ABCD and an AR(1) model for IMAGEN. The resulting voxel-wise GLMs yielded single-trial β maps, which were used as the primary input for subsequent analyses. Crucially, to minimize the impact of head motion, we excluded trials in which framewise displacement (FD) exceeded 0.5 mm within 4–8 seconds after stimulus onset, following procedures commonly adopted in previous studies (Huffman & Ekstrom, 2019; Power et al., 2012; Zheng et al., 2021).

**System-level neural stability**

To characterize the system-level neural stability, we used the Single-Trial Model (STM) and Representational Similarity Analysis (RSA) (Kriegeskorte et al., 2008; Mumford et al., 2012). This approach allowed us to derive a metric quantifying the spatial similarity of activation patterns between pairs of trials within the same control condition (e.g. SS∩SS for comparing two Successful Stop [SS] trials or Go∩ Go for comparing two Go trials). We defined a “system-level neural stability index” by averaging spatial similarity within the key cognitive control networks, excluding sensorimotor regions. Higher values on this metric indicates greater similarity in spatial activation patterns between trial pairs, reflecting higher neural stability within the cognitive control system.

**Supplementary Results**

**Voxel-wise neural stability of reactive control and its developmental trajectory: *Late Childhood to Early Adolescence (ABCD)***

To spatially localize the developmental changes in reactive control stability, we performed a whole-brain searchlight analysis. This approach allowed us to estimate voxel-wise, condition-specific neural stability maps for each participant (see Methods for details), providing a comprehensive assessment of distributed patterns across the cortex.

We identified robust and widespread representational patterns associated with reactive control located in canonical cognitive control systems, including the frontoparietal, salience, visual, and default mode networks (**Fig. S1a**). Direct longitudinal comparisons revealed no significant developmental changes, suggesting minimal group-level developmental change in reactive control during late childhood (**Fig. S1b**).

**Voxel-wise neural stability of reactive control and its developmental trajectory: *Mid-Adolescence to Young Adulthood (IMAGEN)***

In the IMAGEN cohort, we observed a dynamic trajectory of strengthening followed by stabilization. From Age 14 to Age 18 and Age 22, there was a significant increase in the neural stability of reactive control across the bilateral anterior insula (AI), inferior and middle frontal gyri (IFG/MFG), superior frontal gyrus (SFG), pre-SMA/ACC, posterior cingulate cortex (PCC)/precuneus, as well as the bilateral visual and lateral/medial temporal cortices (**Fig. S1b**). Notably, this trajectory plateaued in early adulthood; direct comparisons between Age 18 and Age 22 revealed no significant global differences. This neural plateau mirrors the behavioral findings, where SSRT performance stabilized after age 18. Restricting analysis to the ‘complete-case’ participants yielded highly similar results (**Fig. S3**).

**Voxel-wise neural substrates of dynamic speed–caution decision-making predicts trial-wise RT: *Late Childhood to Early Adolescence (ABCD)***

The neural similarity to the Speed-state and Safe-state templates successfully tracked trial-by-trial behavioral fluctuations at both Age 10 and Age 12. Greater similarity to the Safe-state template showed significant positive correlations with RT (i.e., predicting response slowing) in the SN/Ventral Attention Network (VAN), FPN, and motor areas (**Fig. S5a**). Conversely, greater neural similarity to the Speed-state template exhibited a widespread negative correlation with RT (i.e., predicting faster responses) across frontoparietal, default mode, and visual regions (**Fig. S5b**).

To isolate the competitive dynamic between these control states, independent of shared motor preparation, we computed the Safe-minus-Speed representation difference. This differential metric also predicted RT fluctuations: a neural bias toward the "Safe-state" (positive difference) significantly predicted behavioral slowing across salience, frontoparietal, and visual regions (**Fig. S5c**), confirming that the trial-wise competition between these representations drives behavioral variability.

Longitudinal analyses revealed a significant strengthening of these brain–behavior associations for representations subserving cautious responses. The predictive power of the Safe-state (favoring caution) strengthened in the right inferior temporal lobe (**Fig. S6a**). The Speed–Caution Balance (Safe minus Speed representation) showed stronger coupling with RT in the right frontal pole, bilateral SFG, SMG, and visual cortex, indicating a developmental refinement in how neural competition dictates decision speed (**Fig. S6c**).

**Voxel-wise neural substrates of dynamic speed–caution decision-making predicts trial-wise RT: *Mid-Adolescence to Young Adulthood (IMAGEN)***

In the IMAGEN cohort, we observed robust replication of these effects. The Speed-state consistently predicted faster RTs, while the Safe-state and the Competitive Difference score predicted slowing, with spatial patterns closely mirroring the ABCD dataset (**Fig. S5**).

Longitudinal trends paralleled those observed in childhood. Relative to mid-adolescence (Age 14), young adulthood (both Age 18 and Age 22) exhibited stronger brain–behavior coupling. Specifically, the Safe-state’s association with slowing enhanced in the visual cortex at both Age 18 and Age 22, with additional clusters in bilateral AI, pre-SMA, M/SFG, SMG, and Precuneus at Age 18 (**Fig. S6a**). The speed–caution balance showed similarly stronger coupling in the visual cortex at both Age 18 and Age 22, with additional clusters emerging at Age 22 in the IFG, MFG, insula, ACC, postcentral gyrus, SPL, PCC/precuneus, and subcortical regions including the thalamus, caudate, and putamen (**Fig. S5c**). While the magnitude of Speed-state’s negative association strength was attenuated at Age 18 than BL in the pre-SMA, IFG, MFG/Frontal pole, and SMG. Crucially, no significant differences were observed between Age 18 and Age 22, indicating that the neural mechanisms translating control states into behavioral speed stabilize by late adolescence (**Fig. S6b**). We also conducted additional control analyses using group-wise templates (**Fig. S7-8**) and restricted to complete-case participants, which both yielded highly similar results (**Fig. S9**).

**Brain-behavior coupling of Speed-state and Safe-state states predicts developmental change in SSRT and IIRV: *Cross-wave prediction (ABCD)***

The Safe-state and Speed-state brain–behavior coupling robustly predicted individual differences in both SSRT and IIRV across Age 10 and Age 12 (ps < 0.001). Prediction of SSRT showed trend-level improvements from age 10 to 12 using both Safe-state (p = 0.07) and Speed-state (p = 0.078) representations (**Fig. 2d**). Moreover, these models demonstrate robust generalizability with cross-wave validation (i.e. model trained on Age 10 and tested on Age 12 or vice versa) (all ps < 0.001, **Fig. S10**).

**Brain-behavior coupling of Speed-state and Safe-state states predicts developmental change in SSRT and IIRV: *Cross-wave prediction (IMAGEN)***

Similar to the findings in the ABCD data, both Safe-state and Speed-state coupling patterns robustly predicted SSRT and IIRV within each wave of the IMAGEN data (all ps < 0.001; Fig. 6a-b). Trained models demonstrated reliable between-wave generalization across all train–test pairs (all ps < 0.001, **Fig. S10**). These findings remained robust and significant when accounting for confounding factors including age, sex, head motion, and scanner site, see **Table S4**.

**Brain-behavior coupling of Speed-state and Safe-state states predicts developmental change in SSRT and IIRV: *Cross-cohort prediction***

Elastic Net models trained within a given cohort/wave to predict SSRT and IIRV showed robust transfer across developmental waves and across cohorts (all ps < 0.05, **Fig. S10** and **Table S4**).

**Characterizing representational coherence of Speed-state and Safe-state states**

We mapped the differential topology of Speed- and Safe-state representations by directly contrasting their connectivity strength. This analysis revealed distinct organizational principles: Safe-state coherence showed widespread, high-strength connectivity across the brain, consistent with a globally integrated network architecture. In contrast, Speed-state coherence was comparatively spatially restricted, with stronger connectivity concentrated within focal edges linking sensorimotor, dorsal attention, and striatal regions. These topological distinctions were robust and highly consistent across trial subsets (Go, SS, US trials) and persisted when all trials were analyzed together (**Fig. S11**). The differential connectivity pattern between Speed- and Safe-state states replicated across all five waves in both the ABCD and IMAGEN datasets. Accordingly, subsequent coherence analyses were conducted using all trials.

**Representational coherence predicts inhibitory control and behavioral stability: *Universal generalizability of behavioral stability***

We found remarkable cross-dataset robustness for the prediction of behavioral stability (IIRV). Models trained on ABCD significantly predicted IIRV in all IMAGEN waves (Age 14, 18, 22), with the Safe-state network achieving r = 0.11–0.22 (p < 0.001) and the Speed-state network r = 0.10–0.15 (p < 0.01). This generalization was bidirectional: models trained on IMAGEN successfully predicted IIRV in ABCD waves, with Safe-state r = 0.11–0.15 (p < 0.001) and Speed-state r = 0.09–0.11 (p < 0.05), see **Fig. S12.** And these effects remained significant after adjusting for confounding factors, see **Table S5**. These results indicate that the representation coherence of Safe-state and Speed-state states capture a scanner-invariant, trait-like signature of performance variability that persists across population and developmental stages.

Selective generalization of inhibitory control: In contrast, the prediction of inhibitory control (SSRT) was more specific. The Safe-state network trained on ABCD Age 10 significantly predicted IMAGEN Age 14 (r = 0.06, p = 0.037) and Age 18 (r = 0.09, p = 0.003), while the ABCD Age 12 model predicted IMAGEN Age 18 (r = 0.09, p = 0.001) but failed to predict Age 22 (p > 0.05) (**Fig. S12** and **Table S5**). The Safe-state network trained on IMAGEN Age 18 predicted SSRT in ABCD waves (r = 0.10-012, p < 0.001, Fig. 9b). However, other IMAGEN waves did not generalize consistently.

Robustness check: We also performed additional control analyses in which representational coherence matrices were computed using Go trials only, and the results were highly consistent with the primary findings (**Fig. S13**).

**Replication of developmental effects in spatial stability of reactive and proactive control using the IMAGEN complete-case sample**

To maximize statistical power, we tested developmental effects using a pairwise-available approach for each longitudinal contrast. For each comparison (Age 14 to Age 18, Age 18 to Age 22, and Age 14 to Age 22), we included all participants with data at both time points, regardless of their availability at the third wave; participants missing either time point were excluded from that contrast. To ensure that our results were not confounded by attrition bias or missing data patterns, we conducted a sensitivity analysis using the 'complete-case' subsample, restricting to participants with data at all three waves, and obtained highly similar results (**Fig. S5**), indicating that the findings were robust and not driven by the sample inclusion strategy.

**Replicating the development effects in spatial stability underlying reactive and proactive control when matching the scan length between waves for IMAGEN**

Because Age 14 from IMAGEN dataset included more trials (480) than the follow-up waves (360), trial pairs at baseline could span longer temporal lags and potentially bias pattern-similarity estimates despite our lag-matching procedure (see Methods). To rule out this confound, we conducted additional control analysis after matching baseline scan duration to Age18/Age22 and obtained highly similar findings (**Fig. S6**).

**Replication of developmental effects on the neural substrates of the Speed–Caution balance using the group-wise templates**

To assess the robustness of the Speed–Caution balance findings and controlling potential confounding factors of temporal autocorrelation, we repeated the primary analyses using leave-one-subject-out (LOSO) group-average templates for both Speed-state and Safe-state representations. Specifically: For each participant, we computed participant-level Speed-state and Safe-state summary patterns by averaging β maps across the fastest 10% of Go trials (Speed-state) and across successful stop trials (Safe-state). For a given target participant, we formed LOSO templates by averaging the corresponding participant-level patterns from all other participants (excluding the target participant) (Chen et al., 2017; Gao et al., 2025). We then quantified trial-wise similarity between the target participant’s Go-trial β maps and these LOSO templates, producing Speed-state and Safe-state similarity time series that are independent of the target participant’s own data. These trial-wise similarity values were related to RT using the same searchlight and group-level permutation testing pipeline. The LOSO procedure was applied consistently in both ABCD and IMAGEN. Results closely replicated the main findings reported in the main text (**Fig. S7–S9**).

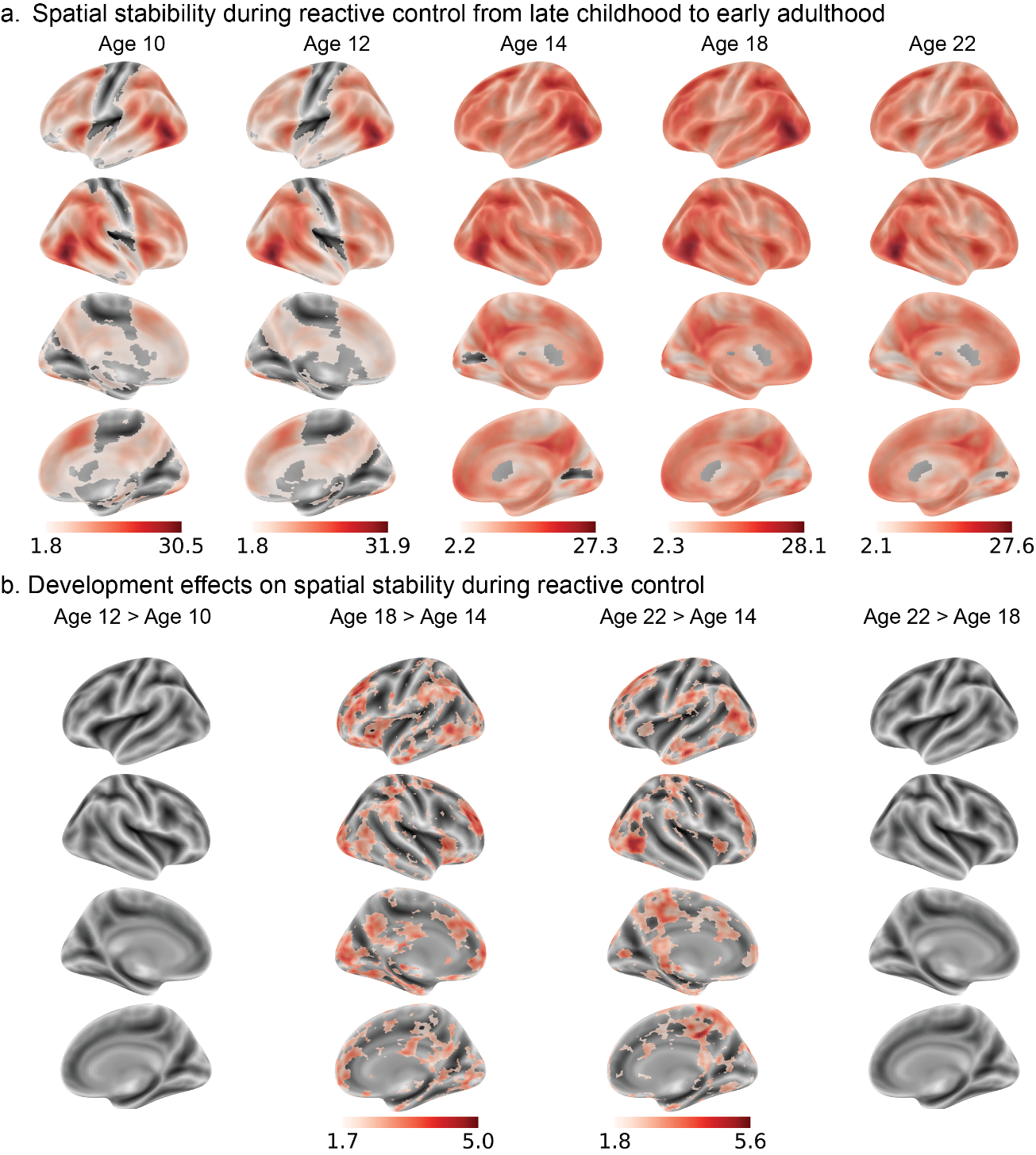

**Fig. S1.** Longitudinal development of reactive control from late childhood to early adulthood. (a) Significant effects of spatial stability underlying reactive control in each wave. (b). Spatial stability supporting reactive control increased from mid-adolescence (≈14 years) to 18 years and to early adulthood (≈22 years). No significant difference was observed between 18 and 22, nor between late childhood and early adolescence. Statistical maps were corrected using TFCE (p < 0.05). Whole-brain plots display voxel-wise t-values (rather than p-values) to avoid visually uniform maps after correction.

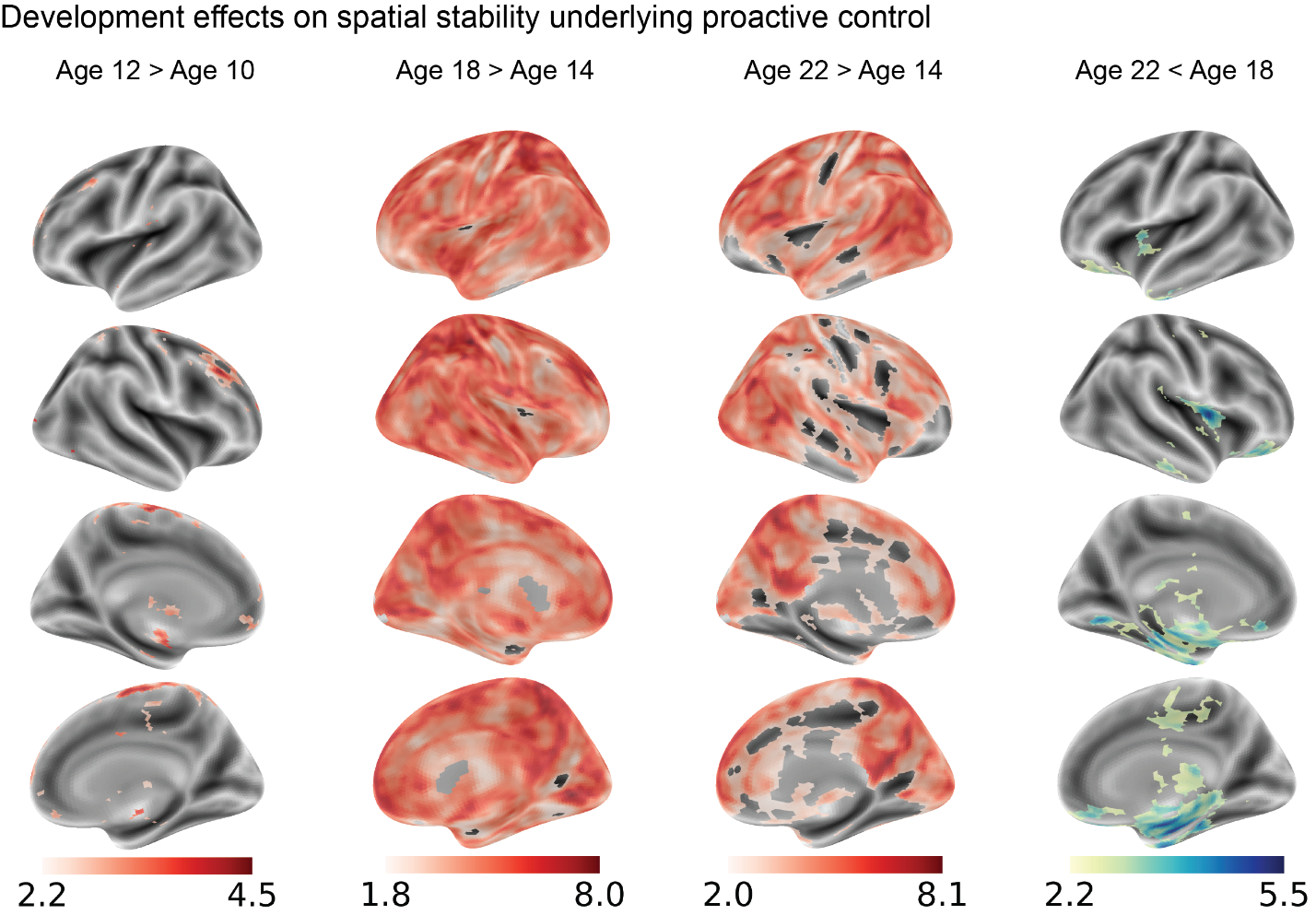

**Fig. S2.** Longitudinal development of proactive control from late childhood to early adulthood. Spatial stability supporting proactive control increased linearly from late childhood through adolescence, plateaued around late adolescence (≈18 years), and then declined by early adulthood (≈22 years). Statistical maps were corrected using TFCE (p < 0.05). Whole-brain plots display voxel-wise t-values (rather than p-values) to avoid visually uniform maps after correction.

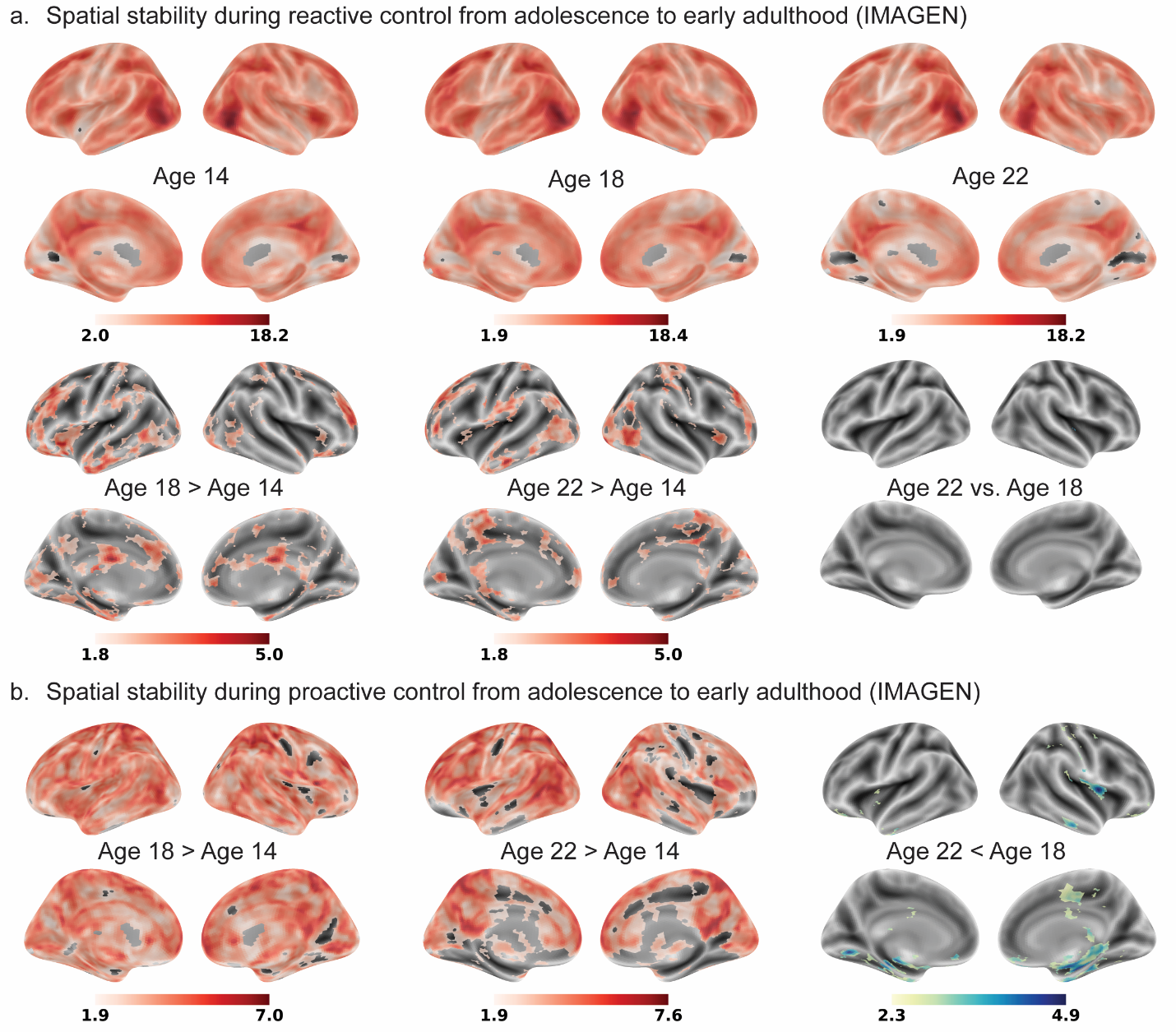

**Fig. S3.** Longitudinal development of reactive (a) and proactive (b) control from adolescence to early adulthood based on a ‘complete-case’ sample from the IMAGEN dataset.

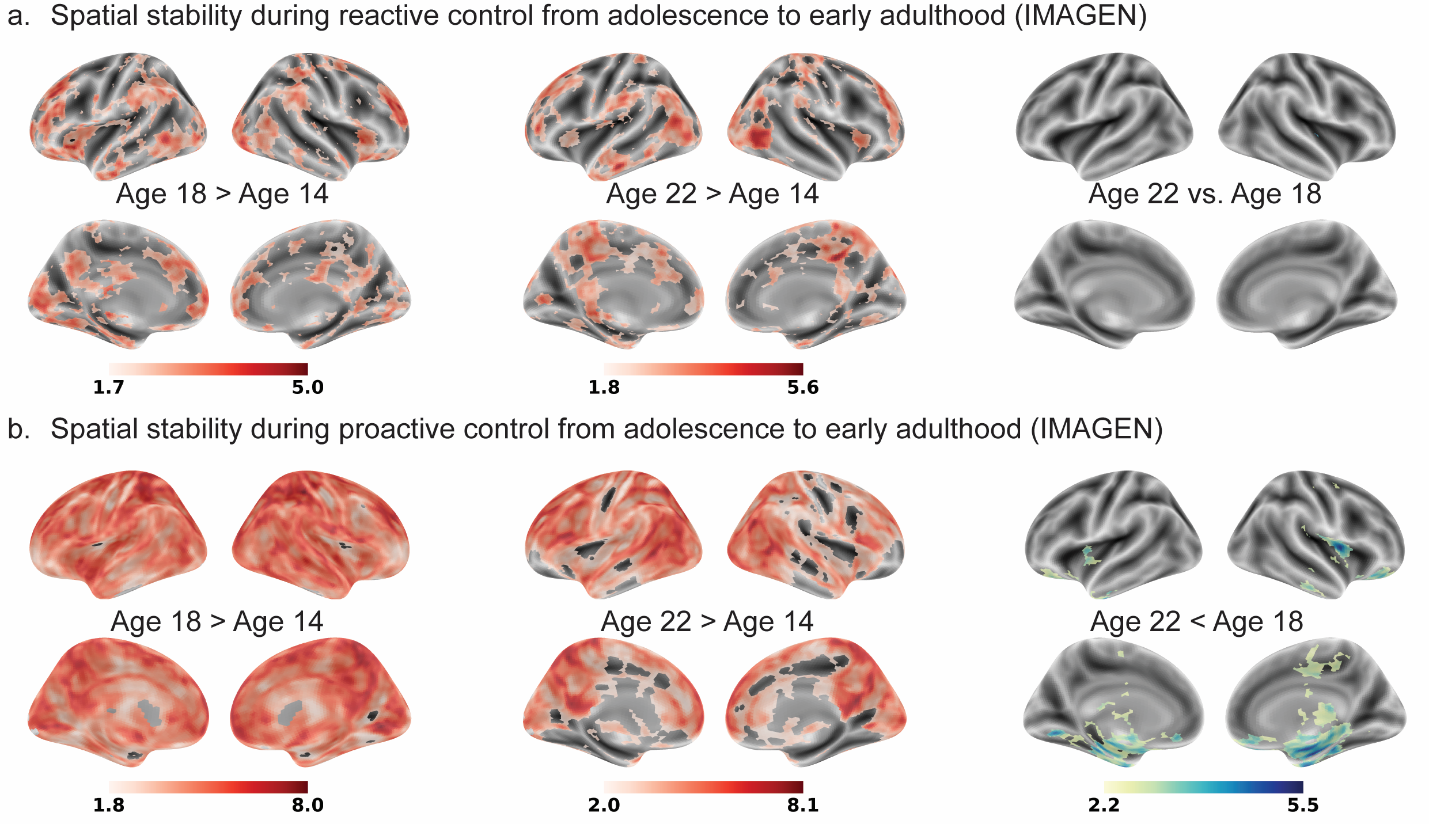

**Fig. S4.** Longitudinal development of reactive (a) and proactive (b) control from adolescence to early adulthood when matching run length across waves in the IMAGEN dataset. Statistical maps were corrected using TFCE (p < 0.05). Whole-brain plots display voxel-wise t-values (rather than p-values) to avoid visually uniform maps after correction.

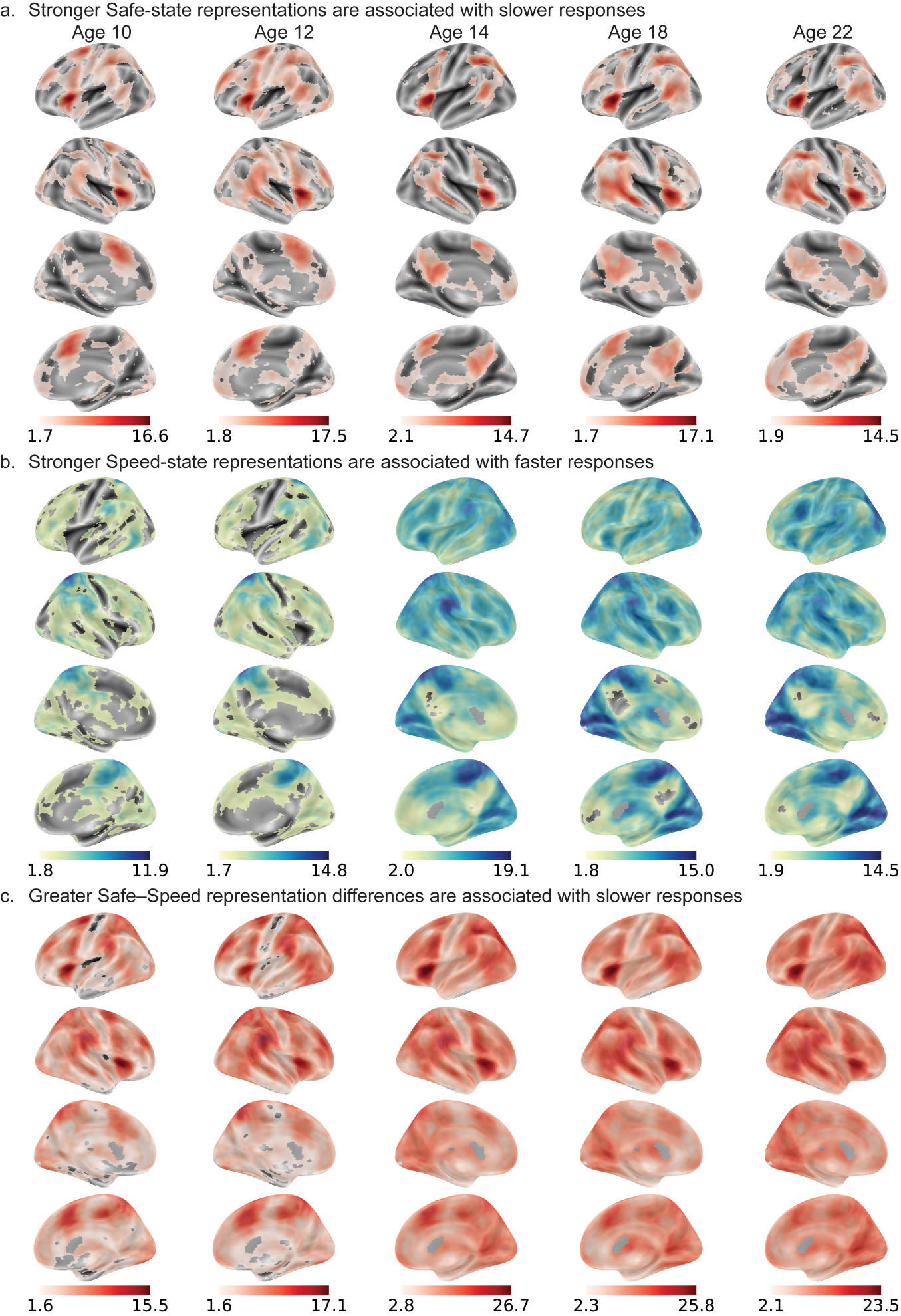

**Fig. S5**. Neural substrates of speed-caution balance. a. Safe-state representation was associated with slower responses. b. Speed-state representation was associated with faster responses. c. The Safe-state and Speed-state representation difference was associated with slower responses. Statistical maps were corrected using TFCE (p < 0.05). Whole-brain plots display voxel-wise t-values (rather than p-values) to avoid visually uniform maps after correction.

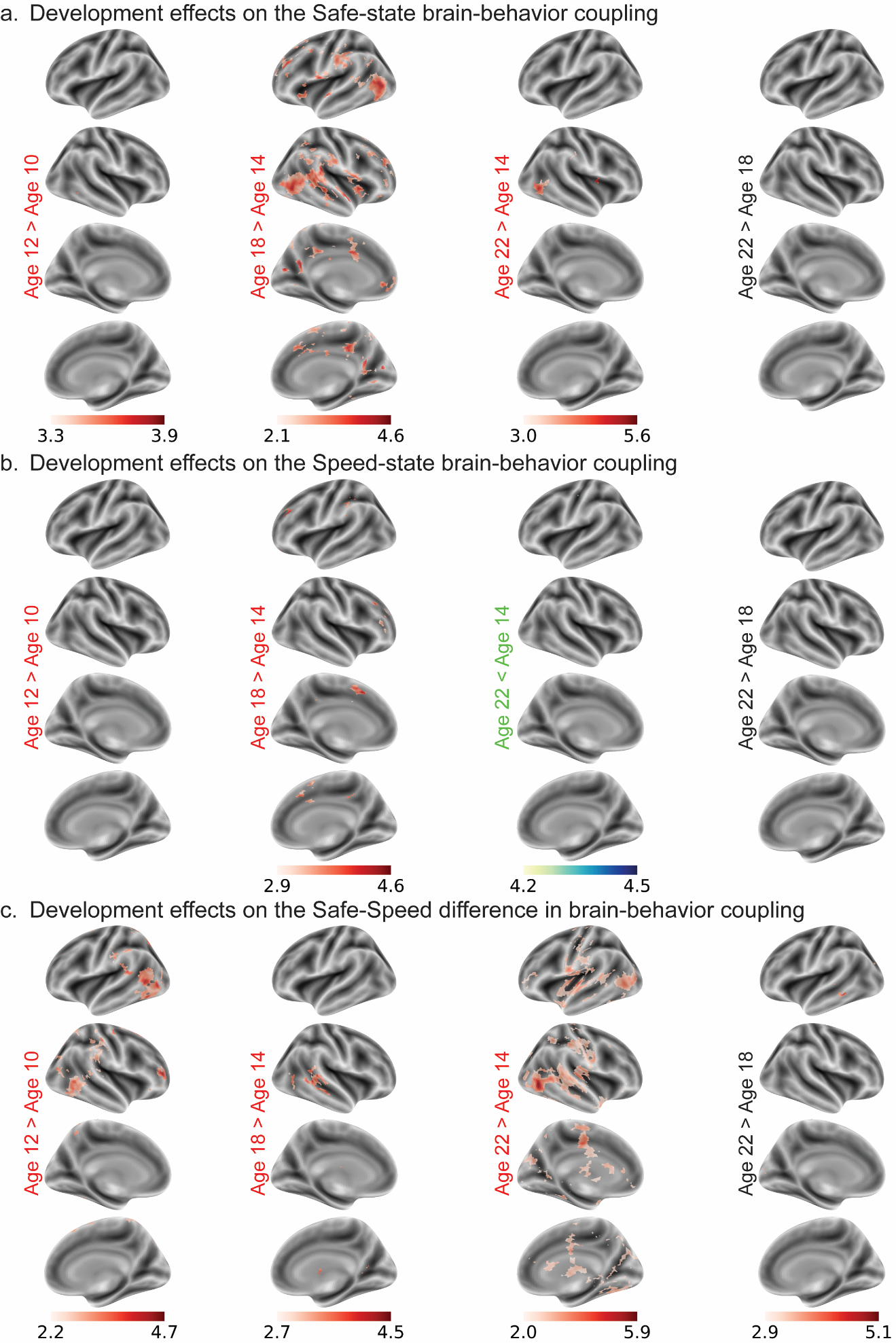

**Fig. S6**. Development effects on Speed-Caution brain-behavior coupling. a. Safe-state brain–behavior coupling strength increased from late childhood through adolescence, plateaued around 18 years, and remained stable into early adulthood (≈22 years). b. Speed-state brain–behavior coupling strength, a negative association with response speed, attenuated from mid-adolescence (≈14 years) to late adolescence (≈18 years), with no significant change from late childhood to early adolescence or from adolescence to early adulthood (≈22 years). c. The Safe–Speed difference in brain–behavior coupling increased linearly from late childhood through adolescence into early adulthood. Statistical maps were corrected using TFCE (p < 0.05). Whole-brain plots display voxel-wise t-values (rather than p-values) to avoid visually uniform maps after correction.

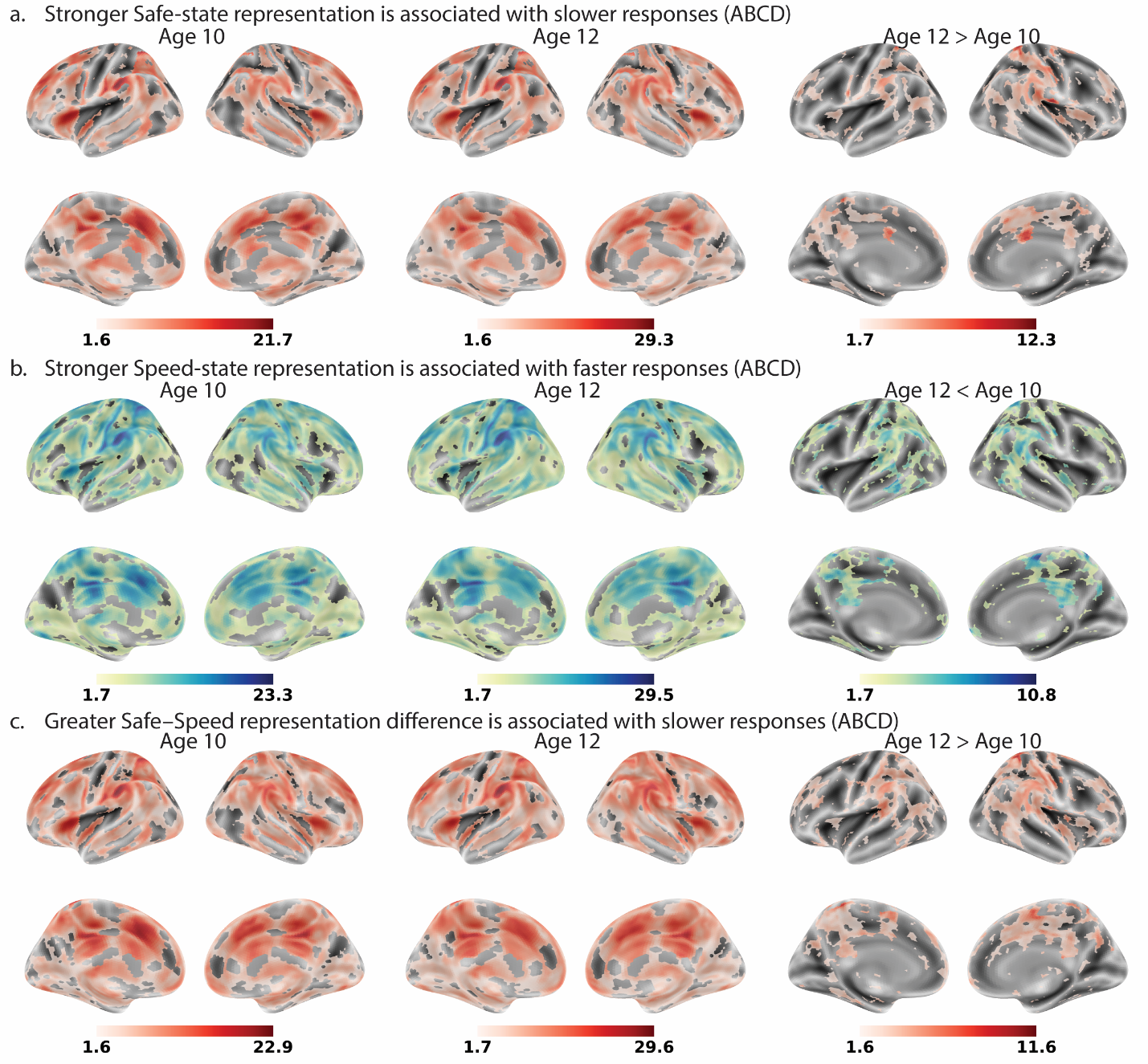

**Fig. S7**. Neural substrates of speed-caution balance based on the group-wise templates in ABCD. a. Safe-state representation was associated with slower responses. b. Speed-state representation was associated with faster responses. c. The Safe-state and Speed-state representation difference was associated with slower responses. Statistical maps were corrected using TFCE (p < 0.05). Whole-brain plots display voxel-wise t-values (rather than p-values) to avoid visually uniform maps after correction.

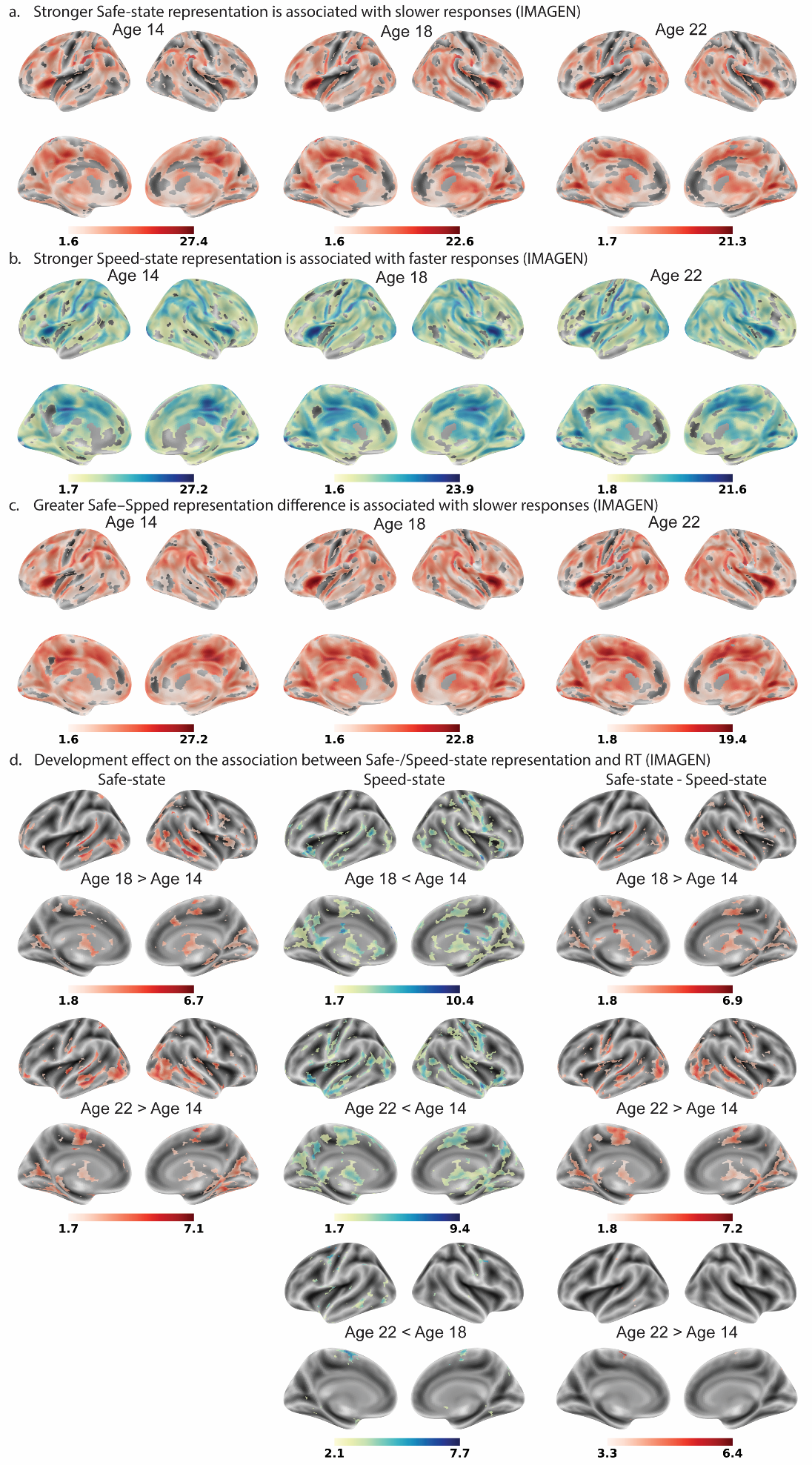

**Fig. S8**. Neural substrates of speed-caution balance based on the group-wise templates in IMAGEN. a. Safe-state representation was associated with slower responses. b. Speed-state representation was associated with faster responses. c. The Safe-state and Speed-state representation difference was associated with slower responses. d. Longitudinal development of brain-behavior coupling with trial-wise RT for Safe-state, Speed-state, and Safe-state and Speed-state difference. Statistical maps were corrected using TFCE (p < 0.05). Whole-brain plots display voxel-wise t-values (rather than p-values) to avoid visually uniform maps after correction.

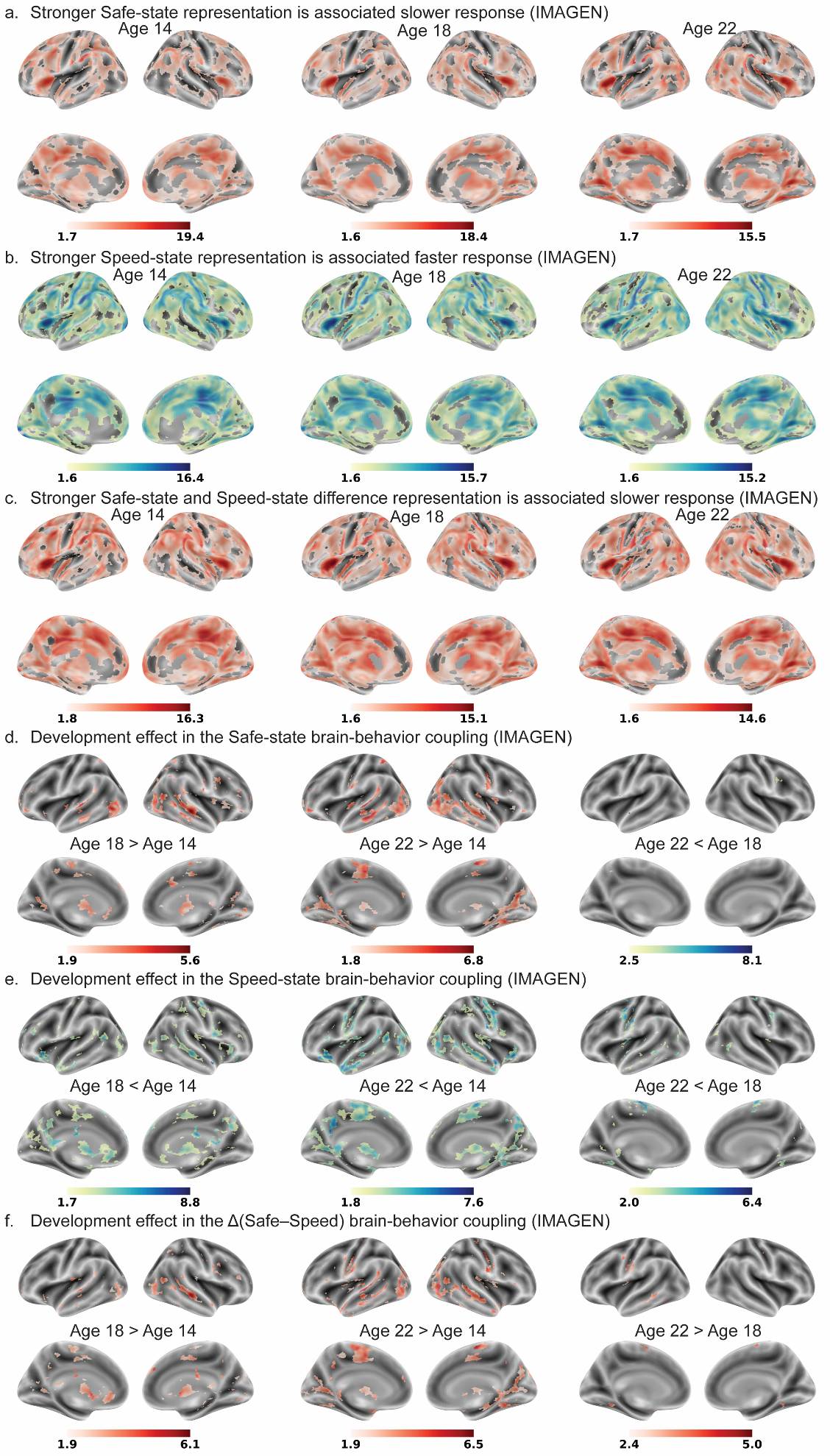

**Fig. S9**. Neural substrates of speed-caution balance based on the group-wise templates and a ‘complete-case’ sample in IMAGEN. a. Safe-state representation was associated with slower responses. b. Speed-state representation was associated with faster responses. c. The Safe-state and Speed-state representation difference was associated with slower responses. d-f. Longitudinal development of brain-behavior coupling with trial-wise RT for Safe-state, Speed-state, and Safe-state and Speed-state difference. Statistical maps were corrected using TFCE (p < 0.05). Whole-brain plots display voxel-wise t-values (rather than p-values) to avoid visually uniform maps after correction.

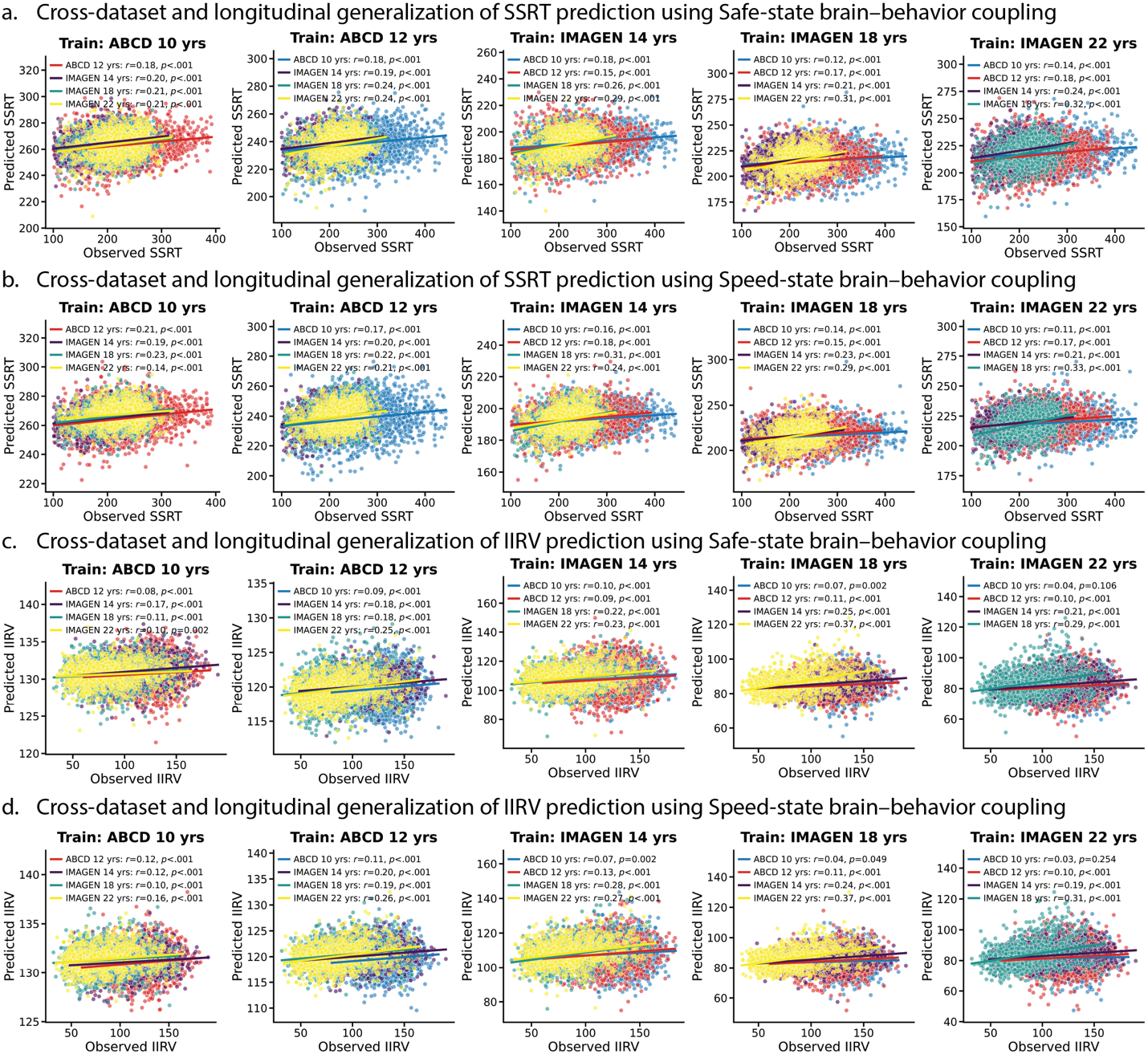

**Fig. S10.** Longitudinal generalization of behavior performance prediction using multi-regional Safe-state/Speed-state brain-behavior coupling. a. Cross-waves and datasets longitudinal generalization of SSRT prediction using Safe-state brain-behavior coupling. b. Cross-waves and datasets longitudinal generalization of SSRT prediction using Speed-state brain-behavior coupling. c. Cross-waves and datasets longitudinal generalization of IIRV prediction using Safe-state brain-behavior coupling. d. Cross-waves and datasets longitudinal generalization of IIRV prediction using Speed-state brain-behavior coupling.

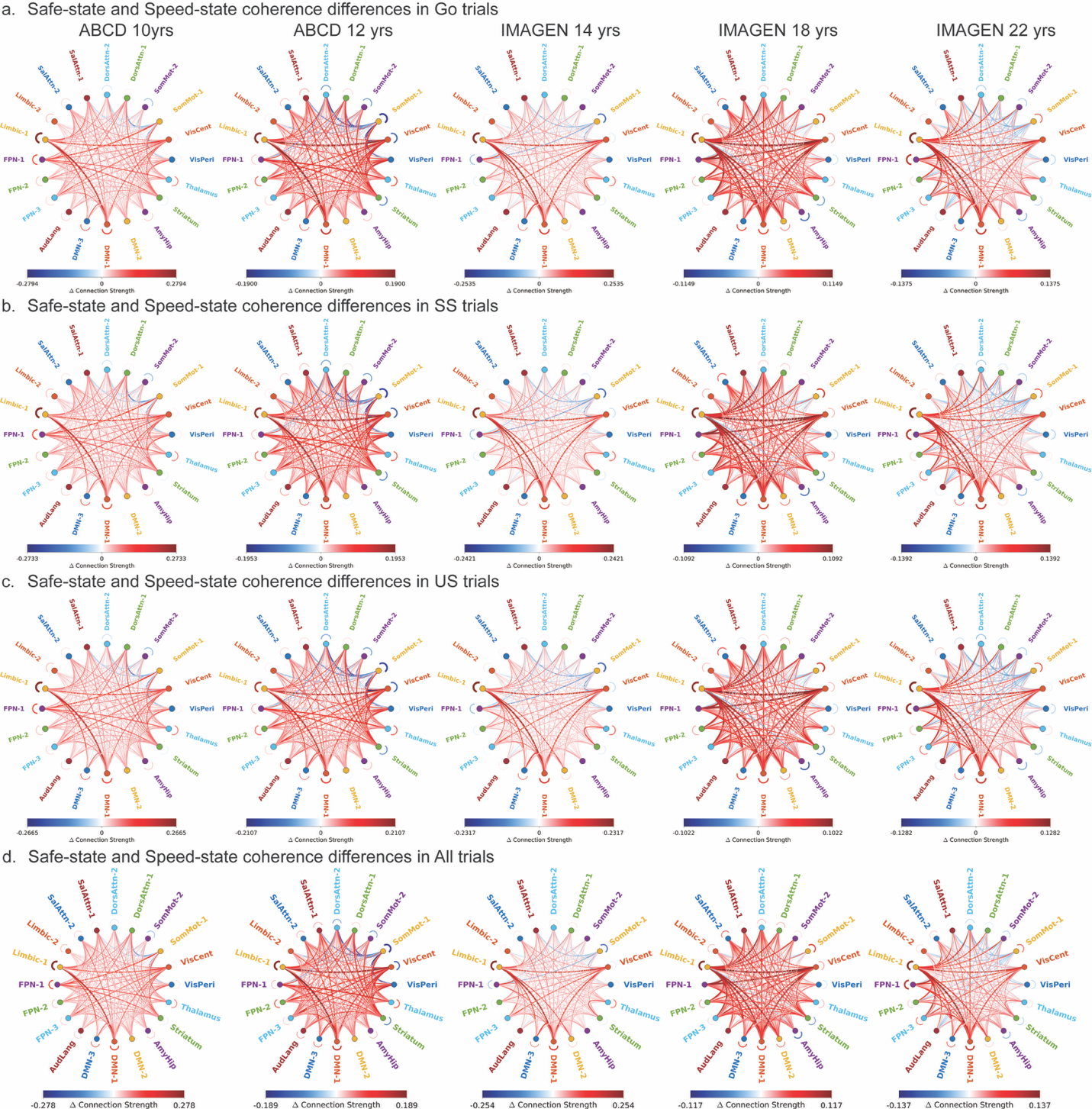

**Fig. S11.** Representational coherence differences between Safe- and Speed-state states across Go (a), SS (b), US (c), and All (d) trials.

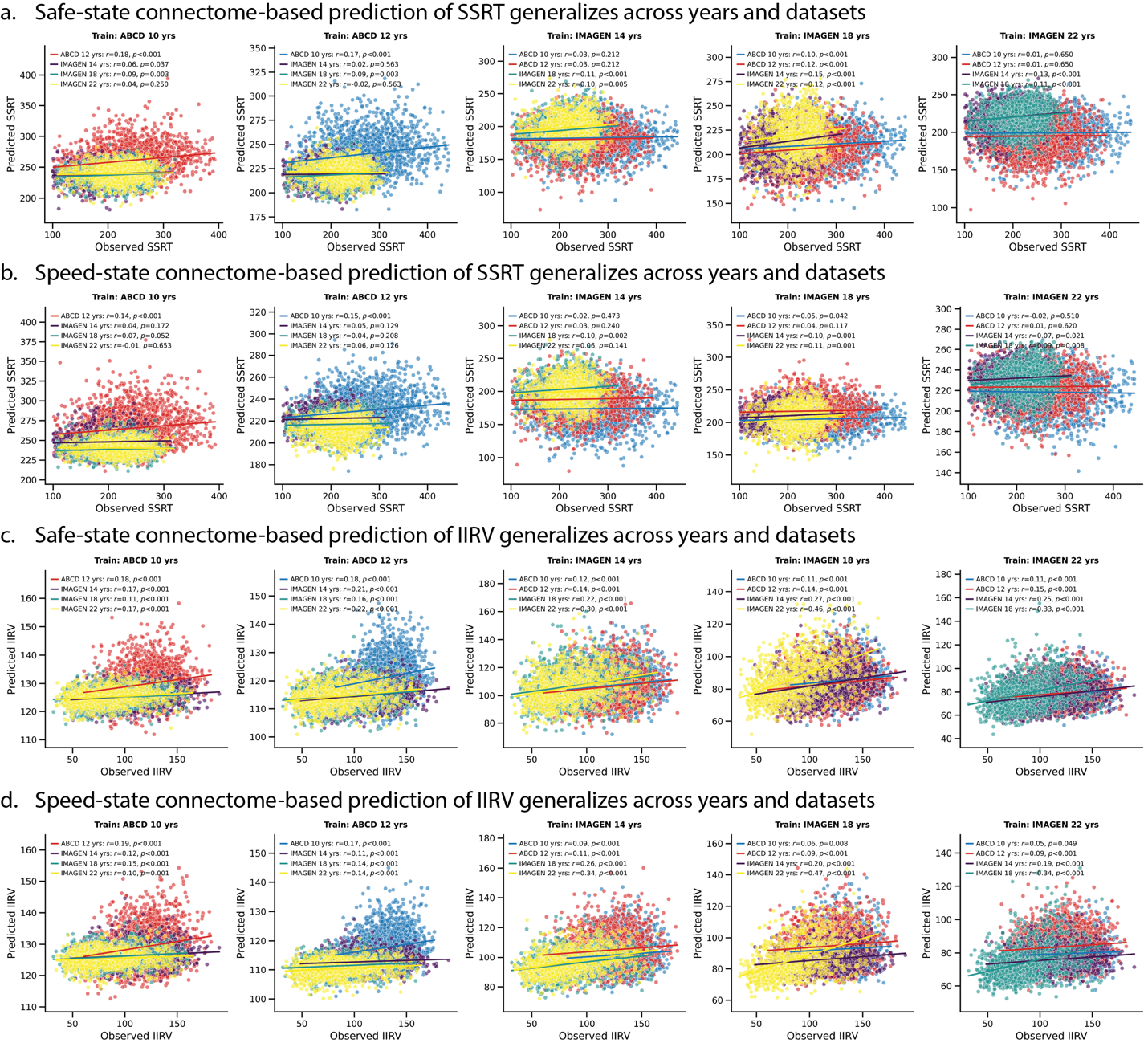

**Fig. S12.** Safe-state/Speed-state connectome-based prediction of behavioral performance generalizes across waves and datasets using all trials. a. Safe-state connectome-based prediction of SSRT generalizes across waves and datasets. b. Speed-state connectome-based prediction of SSRT generalizes across waves and datasets. c. Safe-state connectome-based prediction of IIRV generalizes across waves and datasets. d. Speed-state connectome-based prediction of IIRV generalizes across waves and datasets. All p values were FDR-corrected.

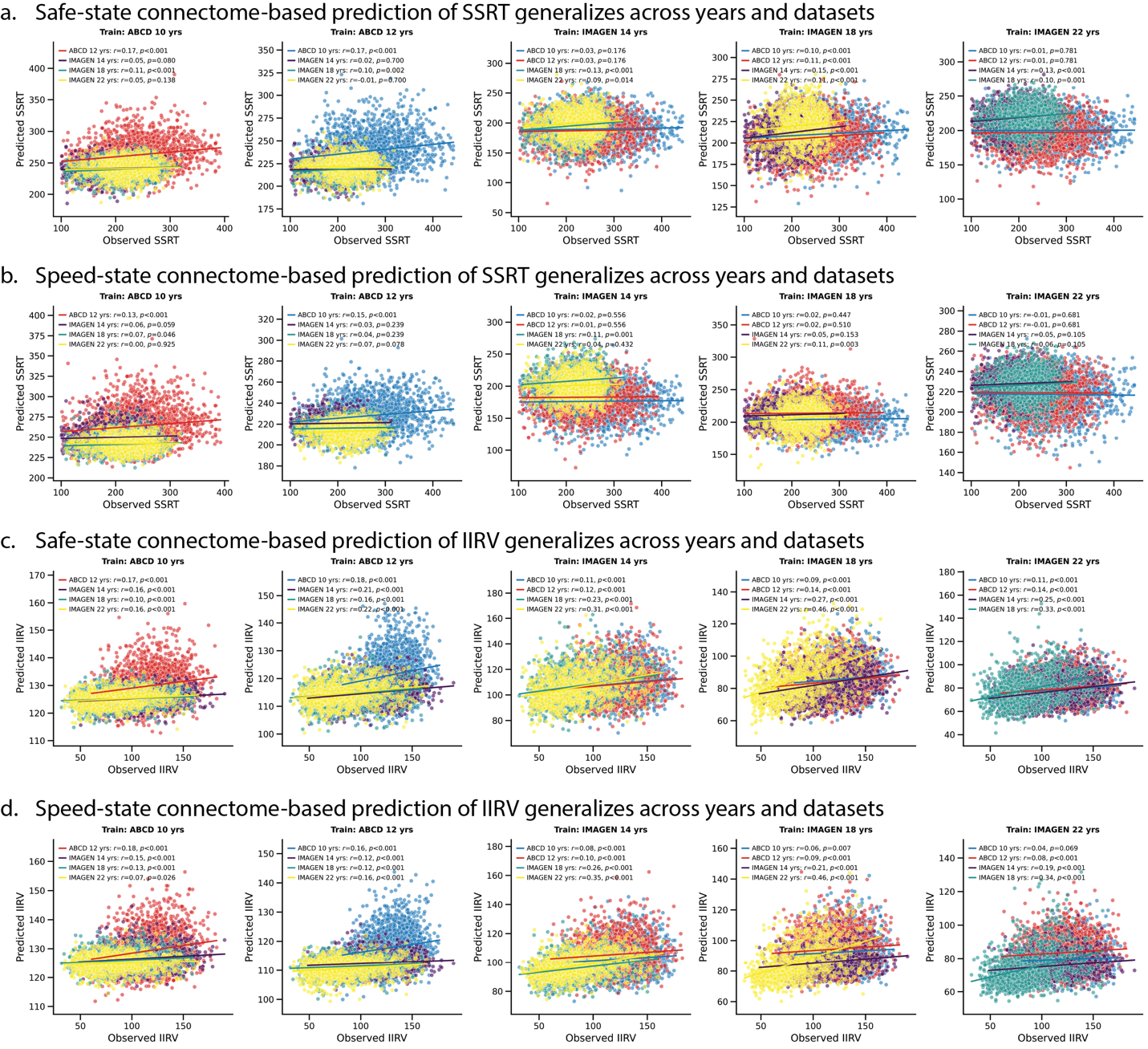

**Fig. S13.** Safe-state/Speed-state connectome-based prediction of behavioral performance generalizes across waves and datasets using only Go trials. a. Safe-state connectome-based prediction of SSRT generalizes across waves and datasets. b. Speed-state connectome-based prediction of SSRT generalizes across waves and datasets. c. Safe-state connectome-based prediction of IIRV generalizes across waves and datasets. d. Speed-state connectome-based prediction of IIRV generalizes across waves and datasets. All p values were FDR-corrected.

**Table S1. Demographics and behavioral results (IMAGEN)**

| Demographics | Age 14 (N=1247) | Age 18  (N = 1151) | Age 22  (N = 973) | p-value |
| --- | --- | --- | --- | --- |
| Age (SD) | 13.97 (2.55) | 18.05 (2.04) | 21.90(1.93) | 0 |
| Gender (Female/Male) | 657/590 | 611/540 | 512/461 | p = 0.972 |
| Head Motion (SD) | 0.11(0.08) | 0.08 (0.06) | 0.07(0.05) | p<0.001 |
| SST Performance |  |  |  |  |
| Go Accuracy (%) | 93.74(4.08) | 94.4(4.08) | 94.3(4.22) | p<0.001 |
| Stop Accuracy (%) | 44.52(4.00) | 50.24(1.69) | 50.22(1.66) | p<0.001 |
| SSRT (ms) | 192.04 (38.86) | 214.72(40.31) | 218.88(38.50) | p<0.001 |
| Go RT (ms) | 444.15(66.42) | 392.49(61.30) | 396.35(66.93) | p<0.001 |
| Go RT STD (ms) | 108.72(26.12) | 86.91(26.93) | 85.20(28.32) | p<0.001 |

Behavioral measures included Go accuracy, Stop accuracy, Go reaction time (GoRT), and stop-signal reaction time (SSRT). Head motion was quantified using mean framewise displacement (FD). Because not all participants had data at every visit, linear mixed-effects models with random intercepts for participants were used to test the main effect of development on behavioral performance and head motion; p-values are reported. Sex distribution differences across visits were tested with chi-square tests.

**Table S2. Demographic and behavioral characteristics of the IMAGEN participants with complete data across BL, FU2, and FU3.**

| Demographics | Age 14 (N=464) | Age 18  (N = 464) | Age 22  (N = 464) | p-value |
| --- | --- | --- | --- | --- |
| Age (SD) | 13.90 (0.84) | 17.82 (2.27) | 21.79(2.06) | 0 |
| Gender (Female/Male) | 258/206 | 258/206 | 258/206 |  |
| Head Motion (SD) | 0.10(0.08) | 0.07(0.05) | 0.07(0.04) | p<0.001 |
| SST Performance |  |  |  |  |
| Go Accuracy (%) | 94.53(3.52) | 94.59(3.93) | 94.29(4.15) | P=0.196 |
| Stop Accuracy (%) | 44.91(3.93) | 50.10(1.53) | 50.03(1.60) | p<0.001 |
| SSRT (ms) | 193.82 (35.97) | 219.11(37.21) | 222.75(37.29) | p<0.001 |
| Go RT (ms) | 439.74(64.99) | 385.79(56.72) | 385.03(57.46) | p<0.001 |
| Go RT STD (ms) | 104.89(25.04) | 82.81(24.90) | 80.48(25.57) | p<0.001 |

Behavioral measures included Go accuracy, Stop accuracy, Go reaction time (GoRT), and stop-signal reaction time (SSRT). Head motion was quantified using mean framewise displacement (FD). Repeated ANOVAs were used to test the main effect of development on behavioral performance and head motion; p-values are reported.

**Table S3. Demographic and behavioral results (IMAGEN behavioral complete-case sample).**

| Demographics | Age 14 (N=256) | Age 18  (N = 256) | Age 22  (N = 256) | p-value |
| --- | --- | --- | --- | --- |
| Age (SD) | 13.89(0.99) | 17.95(2.06) | 21.68(2.52) | 0 |
| Gender (Female/Male) | 144/112 | 144/112 | 144/112 |  |
| SST Performance |  |  |  |  |
| Go Accuracy (%) | 94.37(3.89) | 94.32(4.14) | 93.73(4.64) | P=0.067 |
| Stop Accuracy (%) | 47.95(1.68) | 49.89(1.40) | 49.86(1.61) | p<0.001 |
| SSRT (ms) | 214.60 (29.14) | 224.74(36.29) | 230.94(34.50) | p<0.001 |
| Go RT (ms) | 418.16(53.92) | 380.66(52.19) | 377.89(51.58) | p<0.001 |
| IIRV (ms) | 96.04(22.12) | 80.19(22.24) | 78.57(23.86) | p<0.001 |

Behavioral measures included Go accuracy, Stop accuracy, Go reaction time (GoRT), stop-signal reaction time (SSRT), and intra-individual response variability (IIRV). Head motion was quantified using mean framewise displacement (FD). Linear mixed model was used to test the main effect of development on behavioral performance; p-values are reported.

**Table S4. Longitudinal generalization of behavior performance prediction using multi-regional Safe-state/Speed-state brain-behavior coupling (adjusting for confounding factors)**

| Neural | Behavior | Train | Test | R | P | Partial_R | Partial_P |
| --- | --- | --- | --- | --- | --- | --- | --- |
| Safe-state | SSRT | Age 10 | Age 12 | 0.183424 | 6.38E-16 | 0.176356 | 8.55E-15 |
| Safe-state | SSRT | Age 10 | Age 14 | 0.203908 | 3.60E-13 | 0.20526 | 2.67E-13 |
| Safe-state | SSRT | Age 10 | Age 18 | 0.208151 | 1.06E-12 | 0.209571 | 7.87E-13 |
| Safe-state | SSRT | Age 10 | Age 22 | 0.206385 | 8.24E-11 | 0.208366 | 5.76E-11 |
| Safe-state | SSRT | Age 12 | Age 10 | 0.182835 | 8.64E-16 | 0.178651 | 4.11E-15 |
| Safe-state | SSRT | Age 12 | Age 14 | 0.193831 | 5.08E-12 | 0.191568 | 9.55E-12 |
| Safe-state | SSRT | Age 12 | Age 18 | 0.235369 | 6.48E-16 | 0.230054 | 3.22E-15 |
| Safe-state | SSRT | Age 12 | Age 22 | 0.240506 | 2.95E-14 | 0.240527 | 3.22E-14 |
| Safe-state | SSRT | Age 14 | Age 10 | 0.176012 | 9.96E-15 | 0.166626 | 2.57E-13 |
| Safe-state | SSRT | Age 14 | Age 12 | 0.150086 | 4.28E-11 | 0.143229 | 3.29E-10 |
| Safe-state | SSRT | Age 14 | Age 18 | 0.261843 | 1.88E-19 | 0.259162 | 4.98E-19 |
| Safe-state | SSRT | Age 14 | Age 22 | 0.286238 | 8.73E-20 | 0.287038 | 7.79E-20 |
| Safe-state | SSRT | Age 18 | Age 10 | 0.119345 | 1.74E-07 | 0.110642 | 1.31E-06 |
| Safe-state | SSRT | Age 18 | Age 12 | 0.16575 | 3.06E-13 | 0.154948 | 1.01E-11 |
| Safe-state | SSRT | Age 18 | Age 14 | 0.206709 | 1.68E-13 | 0.20757 | 1.42E-13 |
| Safe-state | SSRT | Age 18 | Age 22 | 0.306894 | 1.21E-22 | 0.309154 | 6.62E-23 |
| Safe-state | SSRT | Age 22 | Age 10 | 0.142716 | 3.88E-10 | 0.136297 | 2.38E-09 |
| Safe-state | SSRT | Age 22 | Age 12 | 0.176756 | 7.07E-15 | 0.17096 | 5.55E-14 |
| Safe-state | SSRT | Age 22 | Age 14 | 0.241102 | 5.97E-18 | 0.24346 | 3.05E-18 |
| Safe-state | SSRT | Age 22 | Age 18 | 0.321206 | 5.81E-29 | 0.32005 | 1.10E-28 |
| Safe-state | IIRV | Age 10 | Age 12 | 0.084645 | 0.000202 | 0.065452 | 0.004106 |
| Safe-state | IIRV | Age 10 | Age 14 | 0.166961 | 3.00E-09 | 0.163207 | 7.06E-09 |
| Safe-state | IIRV | Age 10 | Age 18 | 0.109711 | 0.00021 | 0.104635 | 0.000417 |
| Safe-state | IIRV | Age 10 | Age 22 | 0.100284 | 0.001855 | 0.095155 | 0.003198 |
| Safe-state | IIRV | Age 12 | Age 10 | 0.094197 | 3.48E-05 | 0.081856 | 0.000328 |
| Safe-state | IIRV | Age 12 | Age 14 | 0.177727 | 2.61E-10 | 0.182803 | 8.26E-11 |
| Safe-state | IIRV | Age 12 | Age 18 | 0.175211 | 2.72E-09 | 0.178275 | 1.49E-09 |
| Safe-state | IIRV | Age 12 | Age 22 | 0.246993 | 8.01E-15 | 0.244635 | 1.60E-14 |
| Safe-state | IIRV | Age 14 | Age 10 | 0.102286 | 6.90E-06 | 0.08299 | 0.00027 |
| Safe-state | IIRV | Age 14 | Age 12 | 0.091825 | 5.49E-05 | 0.082971 | 0.000272 |
| Safe-state | IIRV | Age 14 | Age 18 | 0.224674 | 1.78E-14 | 0.226773 | 1.08E-14 |
| Safe-state | IIRV | Age 14 | Age 22 | 0.231529 | 3.67E-13 | 0.221439 | 4.17E-12 |
| Safe-state | IIRV | Age 18 | Age 10 | 0.068912 | 0.002485 | 0.053239 | 0.019588 |
| Safe-state | IIRV | Age 18 | Age 12 | 0.106308 | 2.96E-06 | 0.107551 | 2.31E-06 |
| Safe-state | IIRV | Age 18 | Age 14 | 0.249499 | 3.77E-19 | 0.23182 | 1.22E-16 |
| Safe-state | IIRV | Age 18 | Age 22 | 0.366237 | 7.13E-32 | 0.356435 | 4.47E-30 |
| Safe-state | IIRV | Age 22 | Age 10 | 0.036831 | 0.106212 | 0.031058 | 0.173506 |
| Safe-state | IIRV | Age 22 | Age 12 | 0.099467 | 1.24E-05 | 0.107026 | 2.58E-06 |
| Safe-state | IIRV | Age 22 | Age 14 | 0.206908 | 1.59E-13 | 0.20376 | 4.00E-13 |
| Safe-state | IIRV | Age 22 | Age 18 | 0.290435 | 1.54E-23 | 0.286813 | 6.49E-23 |
| Speed-state | SSRT | Age 10 | Age 12 | 0.21077 | 1.25E-20 | 0.208014 | 4.28E-20 |
| Speed-state | SSRT | Age 10 | Age 14 | 0.190123 | 1.30E-11 | 0.191144 | 1.06E-11 |
| Speed-state | SSRT | Age 10 | Age 18 | 0.225654 | 1.02E-14 | 0.222428 | 2.66E-14 |
| Speed-state | SSRT | Age 10 | Age 22 | 0.140175 | 1.15E-05 | 0.141412 | 9.94E-06 |
| Speed-state | SSRT | Age 12 | Age 10 | 0.167514 | 1.83E-13 | 0.160404 | 1.94E-12 |
| Speed-state | SSRT | Age 12 | Age 14 | 0.203656 | 3.85E-13 | 0.204453 | 3.31E-13 |
| Speed-state | SSRT | Age 12 | Age 18 | 0.219354 | 5.66E-14 | 0.209089 | 8.90E-13 |
| Speed-state | SSRT | Age 12 | Age 22 | 0.206272 | 8.44E-11 | 0.206101 | 9.35E-11 |
| Speed-state | SSRT | Age 14 | Age 10 | 0.162337 | 1.00E-12 | 0.154235 | 1.34E-11 |
| Speed-state | SSRT | Age 14 | Age 12 | 0.183157 | 7.04E-16 | 0.18094 | 1.66E-15 |
| Speed-state | SSRT | Age 14 | Age 18 | 0.312281 | 2.19E-27 | 0.30814 | 1.31E-26 |
| Speed-state | SSRT | Age 14 | Age 22 | 0.242221 | 1.91E-14 | 0.241882 | 2.29E-14 |
| Speed-state | SSRT | Age 18 | Age 10 | 0.135941 | 2.54E-09 | 0.128311 | 1.94E-08 |
| Speed-state | SSRT | Age 18 | Age 12 | 0.151699 | 2.63E-11 | 0.149354 | 5.51E-11 |
| Speed-state | SSRT | Age 18 | Age 14 | 0.228994 | 2.67E-16 | 0.231046 | 1.55E-16 |
| Speed-state | SSRT | Age 18 | Age 22 | 0.291009 | 2.00E-20 | 0.291475 | 1.98E-20 |
| Speed-state | SSRT | Age 22 | Age 10 | 0.106384 | 3.25E-06 | 0.0995 | 1.37E-05 |
| Speed-state | SSRT | Age 22 | Age 12 | 0.168229 | 1.34E-13 | 0.166664 | 2.36E-13 |
| Speed-state | SSRT | Age 22 | Age 14 | 0.205928 | 2.08E-13 | 0.20661 | 1.85E-13 |
| Speed-state | SSRT | Age 22 | Age 18 | 0.331195 | 8.63E-31 | 0.33179 | 7.98E-31 |
| Speed-state | IIRV | Age 10 | Age 12 | 0.123198 | 5.93E-08 | 0.116884 | 2.79E-07 |
| Speed-state | IIRV | Age 10 | Age 14 | 0.120473 | 1.99E-05 | 0.121268 | 1.80E-05 |
| Speed-state | IIRV | Age 10 | Age 18 | 0.101022 | 0.000646 | 0.09691 | 0.001085 |
| Speed-state | IIRV | Age 10 | Age 22 | 0.159984 | 6.19E-07 | 0.150565 | 2.86E-06 |
| Speed-state | IIRV | Age 12 | Age 10 | 0.106895 | 2.60E-06 | 0.097214 | 1.96E-05 |
| Speed-state | IIRV | Age 12 | Age 14 | 0.195286 | 3.49E-12 | 0.193358 | 6.07E-12 |
| Speed-state | IIRV | Age 12 | Age 18 | 0.186157 | 2.52E-10 | 0.185533 | 3.06E-10 |
| Speed-state | IIRV | Age 12 | Age 22 | 0.264264 | 8.08E-17 | 0.263408 | 1.14E-16 |
| Speed-state | IIRV | Age 14 | Age 10 | 0.06998 | 0.002125 | 0.065319 | 0.004172 |
| Speed-state | IIRV | Age 14 | Age 12 | 0.133974 | 3.65E-09 | 0.136476 | 1.90E-09 |
| Speed-state | IIRV | Age 14 | Age 18 | 0.280236 | 5.82E-22 | 0.277556 | 1.66E-21 |
| Speed-state | IIRV | Age 14 | Age 22 | 0.265809 | 5.27E-17 | 0.237482 | 9.50E-14 |
| Speed-state | IIRV | Age 18 | Age 10 | 0.044887 | 0.048941 | 0.035034 | 0.124692 |
| Speed-state | IIRV | Age 18 | Age 12 | 0.11209 | 8.26E-07 | 0.114887 | 4.44E-07 |
| Speed-state | IIRV | Age 18 | Age 14 | 0.238251 | 1.49E-17 | 0.220154 | 4.04E-15 |
| Speed-state | IIRV | Age 18 | Age 22 | 0.371168 | 9.35E-33 | 0.361262 | 6.60E-31 |
| Speed-state | IIRV | Age 22 | Age 10 | 0.026007 | 0.254078 | 0.021406 | 0.348271 |
| Speed-state | IIRV | Age 22 | Age 12 | 0.09554 | 2.70E-05 | 0.106674 | 2.79E-06 |
| Speed-state | IIRV | Age 22 | Age 14 | 0.193648 | 5.32E-12 | 0.192265 | 8.01E-12 |
| Speed-state | IIRV | Age 22 | Age 18 | 0.309374 | 1.21E-26 | 0.305602 | 6.10E-26 |

**Table S5. Safe-state/Speed-state connectome-based prediction of behavioral performance generalizes across waves and datasets (adjusting for confounding factors)**

| Neural | Beh | Train | Test | R | P | R_Partial | P_Partial |
| --- | --- | --- | --- | --- | --- | --- | --- |
| Speed-state | SSRT | Age 10 | Age 12 | 0.139958 | 8.27E-10 | 0.106581 | 3.70E-06 |
| Speed-state | SSRT | Age 10 | Age 14 | 0.042986 | 0.129228 | 0.056403 | 0.047333 |
| Speed-state | SSRT | Age 10 | Age 18 | 0.065615 | 0.026205 | 0.026715 | 0.367914 |
| Speed-state | SSRT | Age 10 | Age 22 | -0.01445 | 0.653131 | 0.019513 | 0.545934 |
| Safe-state | SSRT | Age 10 | Age 12 | 0.181203 | 1.51E-15 | 0.138019 | 1.92E-09 |
| Safe-state | SSRT | Age 10 | Age 14 | 0.062429 | 0.02749 | 0.058065 | 0.041164 |
| Safe-state | SSRT | Age 10 | Age 18 | 0.092729 | 0.00166 | 0.082881 | 0.005147 |
| Safe-state | SSRT | Age 10 | Age 22 | 0.036949 | 0.250274 | 0.052974 | 0.100932 |
| Speed-state | IIRV | Age 10 | Age 12 | 0.189134 | 6.28E-17 | 0.127914 | 2.40E-08 |
| Speed-state | IIRV | Age 10 | Age 14 | 0.117976 | 2.96E-05 | 0.040868 | 0.150859 |
| Speed-state | IIRV | Age 10 | Age 18 | 0.148926 | 4.55E-07 | 0.104478 | 0.000443 |
| Speed-state | IIRV | Age 10 | Age 22 | 0.103035 | 0.001398 | 0.056288 | 0.083082 |
| Safe-state | IIRV | Age 10 | Age 12 | 0.180334 | 1.66E-15 | 0.105429 | 4.36E-06 |
| Safe-state | IIRV | Age 10 | Age 14 | 0.172964 | 7.84E-10 | 0.088507 | 0.001834 |
| Safe-state | IIRV | Age 10 | Age 18 | 0.111672 | 0.000161 | 0.048221 | 0.105673 |
| Safe-state | IIRV | Age 10 | Age 22 | 0.167988 | 1.67E-07 | 0.084985 | 0.00881 |
| Speed-state | SSRT | Age 12 | Age 10 | 0.150347 | 4.50E-11 | 0.081951 | 0.00039 |
| Speed-state | SSRT | Age 12 | Age 14 | 0.047006 | 0.097082 | 0.051416 | 0.070652 |
| Speed-state | SSRT | Age 12 | Age 18 | 0.037196 | 0.207906 | 0.034273 | 0.247988 |
| Speed-state | SSRT | Age 12 | Age 22 | 0.059716 | 0.063011 | 0.079303 | 0.013979 |
| Safe-state | SSRT | Age 12 | Age 10 | 0.171441 | 5.32E-14 | 0.099437 | 1.66E-05 |
| Safe-state | SSRT | Age 12 | Age 14 | 0.016426 | 0.56225 | 0.016441 | 0.563468 |
| Safe-state | SSRT | Age 12 | Age 18 | 0.094052 | 0.001421 | 0.078222 | 0.008293 |
| Safe-state | SSRT | Age 12 | Age 22 | -0.01858 | 0.563191 | 0.002616 | 0.935495 |
| Speed-state | IIRV | Age 12 | Age 10 | 0.167354 | 1.65E-13 | 0.133397 | 6.05E-09 |
| Speed-state | IIRV | Age 12 | Age 14 | 0.106735 | 0.000159 | 0.034074 | 0.231091 |
| Speed-state | IIRV | Age 12 | Age 18 | 0.138752 | 2.65E-06 | 0.098689 | 0.000908 |
| Speed-state | IIRV | Age 12 | Age 22 | 0.143023 | 8.74E-06 | 0.098017 | 0.002505 |
| Safe-state | IIRV | Age 12 | Age 10 | 0.181079 | 1.36E-15 | 0.131281 | 1.05E-08 |
| Safe-state | IIRV | Age 12 | Age 14 | 0.214229 | 2.07E-14 | 0.148395 | 1.58E-07 |
| Safe-state | IIRV | Age 12 | Age 18 | 0.157822 | 8.83E-08 | 0.104808 | 0.000425 |
| Safe-state | IIRV | Age 12 | Age 22 | 0.221408 | 4.10E-12 | 0.163382 | 4.18E-07 |
| Speed-state | SSRT | Age 14 | Age 10 | 0.016464 | 0.473241 | 0.025567 | 0.269268 |
| Speed-state | SSRT | Age 14 | Age 12 | 0.030726 | 0.179733 | 0.029378 | 0.2033 |
| Speed-state | SSRT | Age 14 | Age 18 | 0.102405 | 0.000511 | 0.096436 | 0.001125 |
| Speed-state | SSRT | Age 14 | Age 22 | 0.058077 | 0.070609 | 0.085162 | 0.00829 |
| Safe-state | SSRT | Age 14 | Age 10 | 0.028624 | 0.212345 | 0.057657 | 0.012666 |
| Safe-state | SSRT | Age 14 | Age 12 | 0.030947 | 0.176623 | 0.047628 | 0.039088 |
| Safe-state | SSRT | Age 14 | Age 18 | 0.113274 | 0.00012 | 0.156552 | 1.11E-07 |
| Safe-state | SSRT | Age 14 | Age 22 | 0.097276 | 0.002422 | 0.108891 | 0.000726 |
| Speed-state | IIRV | Age 14 | Age 10 | 0.087351 | 0.000128 | 0.063038 | 0.006171 |
| Speed-state | IIRV | Age 14 | Age 12 | 0.106737 | 2.75E-06 | 0.059151 | 0.010108 |
| Speed-state | IIRV | Age 14 | Age 18 | 0.258132 | 9.17E-19 | 0.221276 | 5.76E-14 |
| Speed-state | IIRV | Age 14 | Age 22 | 0.343267 | 6.59E-28 | 0.289101 | 9.99E-20 |
| Safe-state | IIRV | Age 14 | Age 10 | 0.119494 | 1.54E-07 | 0.064019 | 0.005415 |
| Safe-state | IIRV | Age 14 | Age 12 | 0.141032 | 5.35E-10 | 0.079537 | 0.000538 |
| Safe-state | IIRV | Age 14 | Age 18 | 0.222337 | 3.35E-14 | 0.179971 | 1.17E-09 |
| Safe-state | IIRV | Age 14 | Age 22 | 0.297328 | 5.00E-21 | 0.219983 | 7.29E-12 |
| Speed-state | SSRT | Age 18 | Age 10 | 0.049244 | 0.031843 | 0.066606 | 0.003967 |
| Speed-state | SSRT | Age 18 | Age 12 | 0.035913 | 0.116838 | 0.031856 | 0.167714 |
| Speed-state | SSRT | Age 18 | Age 14 | 0.095451 | 0.000738 | 0.110665 | 9.61E-05 |
| Speed-state | SSRT | Age 18 | Age 22 | 0.112464 | 0.000449 | 0.20664 | 1.02E-10 |
| Safe-state | SSRT | Age 18 | Age 10 | 0.098115 | 1.83E-05 | 0.101691 | 1.06E-05 |
| Safe-state | SSRT | Age 18 | Age 12 | 0.118784 | 1.95E-07 | 0.117458 | 3.34E-07 |
| Safe-state | SSRT | Age 18 | Age 14 | 0.15451 | 4.17E-08 | 0.16618 | 4.11E-09 |
| Safe-state | SSRT | Age 18 | Age 22 | 0.116909 | 0.000263 | 0.226741 | 1.17E-12 |
| Speed-state | IIRV | Age 18 | Age 10 | 0.060197 | 0.008381 | 0.035878 | 0.119333 |
| Speed-state | IIRV | Age 18 | Age 12 | 0.094186 | 3.56E-05 | 0.063167 | 0.006013 |
| Speed-state | IIRV | Age 18 | Age 14 | 0.203965 | 3.54E-13 | 0.154038 | 5.19E-08 |
| Speed-state | IIRV | Age 18 | Age 22 | 0.46818 | 2.10E-53 | 0.453773 | 2.21E-49 |
| Safe-state | IIRV | Age 18 | Age 10 | 0.105547 | 3.64E-06 | 0.056515 | 0.0141 |
| Safe-state | IIRV | Age 18 | Age 12 | 0.141892 | 4.19E-10 | 0.093654 | 4.55E-05 |
| Safe-state | IIRV | Age 18 | Age 14 | 0.265788 | 1.31E-21 | 0.209429 | 9.99E-14 |
| Safe-state | IIRV | Age 18 | Age 22 | 0.455647 | 2.47E-50 | 0.434352 | 6.04E-45 |
| Speed-state | SSRT | Age 22 | Age 10 | -0.02004 | 0.382559 | 0.019738 | 0.393759 |
| Speed-state | SSRT | Age 22 | Age 12 | 0.011369 | 0.619695 | 0.033726 | 0.144121 |
| Speed-state | SSRT | Age 22 | Age 14 | 0.072372 | 0.010574 | 0.068161 | 0.016501 |
| Speed-state | SSRT | Age 22 | Age 18 | 0.090898 | 0.00205 | 0.15808 | 8.30E-08 |
| Safe-state | SSRT | Age 22 | Age 10 | 0.010413 | 0.650113 | 0.07091 | 0.002159 |
| Safe-state | SSRT | Age 22 | Age 12 | 0.011313 | 0.621408 | 0.03811 | 0.09882 |
| Safe-state | SSRT | Age 22 | Age 14 | 0.131369 | 3.25E-06 | 0.139163 | 8.95E-07 |
| Safe-state | SSRT | Age 22 | Age 18 | 0.108964 | 0.000217 | 0.234366 | 1.15E-15 |
| Speed-state | IIRV | Age 22 | Age 10 | 0.045051 | 0.048588 | 0.039123 | 0.089405 |
| Speed-state | IIRV | Age 22 | Age 12 | 0.090031 | 7.77E-05 | 0.067445 | 0.003351 |
| Speed-state | IIRV | Age 22 | Age 14 | 0.186359 | 3.30E-11 | 0.140529 | 6.97E-07 |
| Speed-state | IIRV | Age 22 | Age 18 | 0.338545 | 6.92E-32 | 0.321774 | 1.46E-28 |
| Safe-state | IIRV | Age 22 | Age 10 | 0.106873 | 2.74E-06 | 0.06316 | 0.006073 |
| Safe-state | IIRV | Age 22 | Age 12 | 0.147859 | 7.41E-11 | 0.096359 | 2.72E-05 |
| Safe-state | IIRV | Age 22 | Age 14 | 0.249143 | 4.25E-19 | 0.202157 | 7.12E-13 |
| Safe-state | IIRV | Age 22 | Age 18 | 0.332002 | 1.17E-30 | 0.326043 | 2.54E-29 |

**References:**

Casey, B. J., Cannonier, T., Conley, M. I., Cohen, A. O., Barch, D. M., Heitzeg, M. M., Soules, M. E., Teslovich, T., Dellarco, D. V., & Garavan, H. (2018). The adolescent brain cognitive development (ABCD) study: imaging acquisition across 21 sites. *Developmental Cognitive Neuroscience*, *32*, 43–54.

Chen, J., Leong, Y. C., Honey, C. J., Yong, C. H., Norman, K. A., & Hasson, U. (2017). Shared memories reveal shared structure in neural activity across individuals. *Nature Neuroscience*, *20*(1), 115–125.

Gao, Z., Duberg, K., Warren, S. L., Zheng, L., Hinshaw, S. P., Menon, V., & Cai, W. (2025). Reduced temporal and spatial stability of neural activity patterns predict cognitive control deficits in children with ADHD. *Nature Communications*, *16*(1), 2346.

Huffman, D. J., & Ekstrom, A. D. (2019). A modality-independent network underlies the retrieval of large-scale spatial environments in the human brain. *Neuron*, *104*(3), 611–622. e617.

Kriegeskorte, N., Mur, M., & Bandettini, P. A. (2008). Representational similarity analysis-connecting the branches of systems neuroscience. *Frontiers in systems neuroscience*, *2*, 249.

Mascarell Maričić, L., Walter, H., Rosenthal, A., Ripke, S., Quinlan, E. B., Banaschewski, T., Barker, G. J., Bokde, A. L., Bromberg, U., & Büchel, C. (2020). The IMAGEN study: a decade of imaging genetics in adolescents. *Molecular psychiatry*, *25*(11), 2648–2671.

Mumford, J. A., Turner, B. O., Ashby, F. G., & Poldrack, R. A. (2012). Deconvolving BOLD activation in event-related designs for multivoxel pattern classification analyses. *Neuroimage*, *59*(3), 2636–2643.

Power, J. D., Barnes, K. A., Snyder, A. Z., Schlaggar, B. L., & Petersen, S. E. (2012). Spurious but systematic correlations in functional connectivity MRI networks arise from subject motion. *Neuroimage*, *59*(3), 2142–2154.

Schumann, G., Loth, E., Banaschewski, T., Barbot, A., Barker, G., Büchel, C., Conrod, P. J., Dalley, J., Flor, H., & Gallinat, J. (2010). The IMAGEN study: reinforcement-related behaviour in normal brain function and psychopathology. *Molecular psychiatry*, *15*(12), 1128–1139.

Volkow, N. D., Koob, G. F., Croyle, R. T., Bianchi, D. W., Gordon, J. A., Koroshetz, W. J., Pérez-Stable, E. J., Riley, W. T., Bloch, M. H., & Conway, K. (2018). The conception of the ABCD study: From substance use to a broad NIH collaboration. *Developmental Cognitive Neuroscience*, *32*, 4–7.

Zeithamova, D., de Araujo Sanchez, M.-A., & Adke, A. (2017). Trial timing and pattern-information analyses of fMRI data. *Neuroimage*, *153*, 221–231.

Zheng, L., Gao, Z., McAvan, A. S., Isham, E. A., & Ekstrom, A. D. (2021). Partially overlapping spatial environments trigger reinstatement in hippocampus and schema representations in prefrontal cortex. *Nature Communications*, *12*(1), 6231.
